# Characterization of exosome release and extracellular vesicle-associated miRNAs for human bronchial epithelial cells irradiated with high charge and energy ions

**DOI:** 10.1101/2020.07.15.204909

**Authors:** Zhentian Li, Kishore K. Jella, Lahcen Jaafar, Carlos S. Moreno, William S. Dynan

## Abstract

Exosomes are extracellular vesicles that mediate transport of nucleic acids, proteins, and other molecules. Prior work has implicated exosomes in the transmission of radiation nontargeted effects. Here we investigate the ability of energetic heavy ions, representative of species found in galactic cosmic rays, to stimulate exosome release from human bronchial epithelial cells *in vitro*. Immortalized human bronchial epithelial cells (HBEC3-KT F25F) were irradiated with 1.0 Gy of high linear energy transfer (LET) ^48^Ti, ^28^Si, or ^16^O ions, or with 10 Gy of low-LET reference γ-rays, and extracellular vesicles were collected from conditioned media. Preparations were characterized by single particle tracking analysis, transmission electron microscopy, and immunoblotting for the exosomal marker, TSG101. Irradiation with high-LET ions, but not γ-rays, stimulated exosome release by about 3-fold, relative to mock-irradiated controls. The exosome-enriched vesicle preparations contained pro-inflammatory damage-associated molecular patterns, including HSP70 and calreticulin. Additionally, miRNA profiling was performed for vesicular RNAs using NanoString technology. The miRNA profile was skewed toward a small number of species that have previously been shown to be involved in cancer initiation and progression, including miR-1246, miR-1290, miR-23a, and miR-205. Additionally, a set of 24 miRNAs was defined as modestly over-represented in preparations from HZE ion-irradiated versus other cells. Gene set enrichment analysis based on the over-represented miRNAs showed highly significant association with nonsmall cell lung and other cancers.

## 1. INTRODUCTION

HZE ions are a form of high-LET radiation. Passage through cells and tissues generates densely ionizing radiation tracks, which result in complex DNA damage that is repaired slowly or not at all (Asaithamby et al., 2008; Kumar et al., 2018; Werner et al., 2017). HZE ions are an important component of galactic cosmic rays (reviewed in (Nelson, 2016)) Exposure is linked to cancer and other diseases and is thus a limiting factor for deep space travel (Cucinotta et al., 2017). Cancer risks of HZE ion exposure arise both from targeted effects in cells that are directly traversed by radiation tracks and from non-targeted effects, defined as effects that arise via communication between irradiated cells and their non-irradiated neighbors (Cucinotta and Cacao, 2017).

Exosomes are extracellular vesicles that originate within multivesicular bodies (reviewed in (Hessvik and Llorente, 2018; Juan and Furthauer, 2018)). They can influence tissue physiology in various ways, including by presentation of ligands on their extraluminal surface or by internalization to recipient cells, with release of proteins, nucleic acids, and other cargo to the cell interior. Cellular stress, including exposure to genotoxic agents, stimulates exosome release (Arscott et al., 2013; Beer et al., 2015; Jella et al., 2014; Lehmann et al., 2008). Prior work has shown that exosomes are capable of transmitting non-targeted radiation effects *in vitro*, including genomic and telomeric instability (Al-Mayah et al., 2015; Al-Mayah et al., 2017; Al-Mayah et al., 2012; Arscott et al., 2013; Jella et al., 2014), reviewed in (Du et al., 2020).

High-LET radiation is a potent inducer of non-targeted effects, and we hypothesized that it might be particularly effective in promoting exosome release. In the present study, we exposed human bronchial epithelial cells to HZE ions, prepared extracellular vesicles, and characterized the resulting exosome-enriched preparations. We investigated the presence of DAMPs, HSP70 and calreticulin, which have previously been shown to contribute to pro-inflammatory effects of exosomes derived from cells exposed to DNA damaging agents (Elsner et al., 2007; Gross et al., 2003).

We also performed miRNA profiling on vesicular preparations. Evidence in a variety of models implicates exosomal miRNA in the regulation of gene expression in recipient cells (reviewed in (Mills et al., 2019; Yu et al., 2016)). Consistent with a role for miRNAs, prior work has shown that RNase treatment abrogates the exosome-mediated transmission of radiation-induced bystander effects (Al-Mayah et al., 2017; Al-Mayah et al., 2012). Our profiling data showed a miRNA distribution that was highly skewed toward a few abundant, cancer-related miRNAs. Analysis also defined a set of RNAs that were enriched in preparations from HZE ion-irradiated versus other cells. Work thus establishes a framework for understanding the potential role of exosomes in mediating effects of HZE ion irradiation.

## 2. Materials and Methods

### 2.1 Cell culture

HBEC3-KT F25F cells were previously derived from HBEC3-KT line (Ramirez et al., 2004) by selection for growth on soft agar following ^56^Fe ion irradiation (Li et al., 2018b). They were cultured using Keratinocyte-SFM medium with 50 µg/mL bovine pituitary extract and 5 ng/mL epidermal growth factor (ThermoFisher Scientific) in a humidified incubator at 37°C with 5% CO_2_.

### 2.2 Irradiation

Irradiation with HZE particles was performed at the NASA Space Radiation Laboratory, Upton, NY. Irradiation was performed in T-150 flasks containing 20 ml medium. These were flipped to a vertical position, orthogonal to the beam, for irradiation. Beam energies and calculated LET values were as follows: ^48^Ti, 1000 MeV/u (LET=108 keV/μm) or 230 MeV/u (LET=200 keV/μm); ^28^Si, 148 MeV/u (LET=100 keV/μm) or 65 MeV/u (LET=200 keV/μm); ^16^O, 35 MeV/u (LET=100 keV/μm). LET values were estimated using the Galactic Cosmic Radiation Event-based Risk Model code (GERMcode v1.1 2000) (Cucinotta et al., 2011). Reference ^137^Cs γ-ray irradiation was performed at Brookhaven National Laboratory using a J.L. Sheppard & Associates MK I-68A irradiator at a dose rate of 1.5 Gy/min. Mock irradiations were performed in parallel with HZE ion or γ-ray irradiations except that flasks were not brought into the target room (HZE ions) or placed in the irradiation chamber (γ-rays).

### 2.3 Extracellular vesicle isolation

Conditioned media was harvested at 24 h post irradiation, passed through a 0.22 μm filter, frozen at - 80°C, and returned to the home laboratory. For extracellular vesicle isolation, 60 ml of conditioned media was thawed, and all subsequent steps were performed at 4 1°C. Media was centrifuged at 10,000*g* for 30 min, and the supernatant was concentrated using 10 kDa cutoff Amicon Ultra-15 centrifugal filter units (MilliporeSigma, Burlington, MA., catalog no. UFC901024). The retentate (2 ml) was subjected to centrifugation at 100,000*g* for 2 h in an Optima MAX-XP benchtop ultracentrifuge with a TLA-100.3 rotor (Beckman Coulter, Inc., Brea, CA). Pellets were resuspended in 100 µl PBS and stored at −80 °C.

### 2.4 Immunoblotting

Extracellular vesicle pellets were resuspended in 100 μl of PBS. For immunoblotting, a 10 μl aliquot was mixed with an equal volume of 2X radioimmunoprecipitation assay buffer and incubated on ice for 30 min. SDS-PAGE sample buffer was added and samples were incubated at 95 °C for 5 min. Proteins were resolved by 4-15% SDS-PAGE and transferred to a Immobilon-FL PVDF membrane (MilliporeSigma, catalog no. IPFL20200). The membrane was blocked with a solution of 5% BSA in TBST (50 mM Tris-HCl, pH 7.5, 150 mM NaCl, 0.1% Tween 20) for 1 h, then incubated overnight at 4 °C with primary antibodies diluted in a solution of 1% BSA in TBST. Primary antibodies were as follows: anti-TSG101 (Abcam, Cambridge, MA; catalog no. ab83, 1:1000), anti-calreticulin (Abcam, catalog no. ab2907, 1:1000), and anti-HSP70 (Enzo Life Sciences, Inc., Farmingdale, NY, cat no. ADI-SPA-810-D, 1:1000). Membranes were washed with TBST, then incubated with a solution of 1% BSA in TBST containing a 1:15,000 dilution of IR Dye 680RD-conjugated goat anti-mouse IgG (LI-COR Biosciences, Lincoln, NE, catalog no. 925-68070). Immune complexes were imaged using a LI-COR Odyssey system and quantified using Image Studio v4.0 (LI-COR Biosciences).

### 2.5 Transmission electron microscopy

A 10 μl aliquot of extracellular vesicles was deposited onto a 400-mesh carbon-coated copper grid, which was washed with distilled water, stained with 1% phosphotungstic acid (pH 6.5), and dried prior to CCD imaging. The imaging was performed using a JEOL JEM-1400 instrument (JEOL, Tokyo, Japan).

### 2.6 Single-particle tracking analysis

A 10 μl aliquot of extracellular vesicles was diluted 1:1000 in PBS and analyzed in triplicate using a NanoSight instrument (Malvern Instruments, Amesbury, UK). Data were analyzed using Nanoparticle tracking analysis software version 2.2.

### 2.7 miRNA profiling

Extracellular vesicles were prepared from 80 ml of conditioned media (4 X T-150 flasks) as in Section 2.3. The vesicular pellet was resuspended in 700 μl of Qiazol lysis buffer and RNA was isolated using a miRNeasy Mini Kit (QIAGEN Inc., Germantown MD, Cat #217004). Preparations were analyzed using an Agilent 2100 Bioanalyzer and Agilent Small RNA kit (Agilent Technologies, Santa Clara CA).

Analysis was performed using the NanoString Human v3 miRNA Assay. Briefly, miRNAs were annealed to target-specific miRtag sequences via a bridge oligonucleotide. Following ligation, excess tag oligonucleotides were removed enzymatically. Purified miRNA-miRtag complexes were incubated for 18 h at 65 ^°^C with tag-specific, biotin-labeled capture probes and fluorescently-labeled reporter probes. Samples were injected into a NanoString SPRINT cartridge and loaded onto the SPRINT instrument where excess capture probe and reporter probe were removed and hybridized and miRNA complexes were immobilized for imaging. Following image acquisition, miRNA counts were extracted from nCounter data files using NanoString nSolver analysis software v3.0.

Statistical analysis was performed using a Significance Analysis of Microarray (SAM) approach (Tusher et al., 2001), considering preparations from HZE ion-irradiated cultures as one group and preparations from γ-ray-or non-irradiated cultures as the other. A set 24 miRNAs, modestly enriched in HZE ion-irradiated cells, was used for gene set enrichment analysis with the TAM 2.0 tool (http://www.scse.hebut.edu.cn/tam/) (Li et al., 2018a).

## 3. Results

### 3.1 HZE ion irradiation stimulates exosome secretion

Studies here were performed *in vitro* using the HBEC3-KT F25F line of human bronchial epithelial cells, which are immortalized and transformed to anchorage-independent growth, but are non-tumorigenic. We have previously used them as a model for precancerous lung cells (Li et al., 2018b). Cultures were irradiated with 1 Gy of ^48^Ti, ^28^Si, or ^16^O ions at the NASA Space Radiation Laboratory. Parallel cultures were mock-irradiated or irradiated with reference γ-rays, using a 10 Gy dose that we separately determined to be optimal for stimulation of exosome release in a tumor cell model (Jella et al., 2020). Media was collected at 24 h post-irradiation. Extracellular vesicles were concentrated by ultrafiltration, isolated by differential centrifugation, and analyzed.

To determine relative exosome content in the extracellular vesicle preparations, we performed immunoblotting for the exosomal marker, TSG101, which is a component of the ESCRT machinery required for exosome biogenesis (Juan and Furthauer, 2018). Irradiation with ^48^Ti increased TSG101 levels by 3.4-fold, relative to a mock-irradiated control (Fig. 1A). Irradiation with ^28^Si and ^16^O, which was performed independently (in a separate NSRL campaign), increased TSG101 levels by an average of 2.8-fold (Fig. 1B). There appeared to be little difference between ^28^Si ions delivered at different energies (calculated to provide LET values of 100 and 200 keV/μm) or between ^28^Si and ^16^O ions. There also appeared to be little difference between γ-ray irradiation and mock-irradiation. For statistical analysis, we therefore considered the HZE ion-irradiated samples as one group and the γ-ray and mock-irradiated samples as another group. The difference in exosome content in preparations from the two groups, as indicated by levels of the TSG101 marker, was statistically significant (P<0.01).

**Fig 1.**
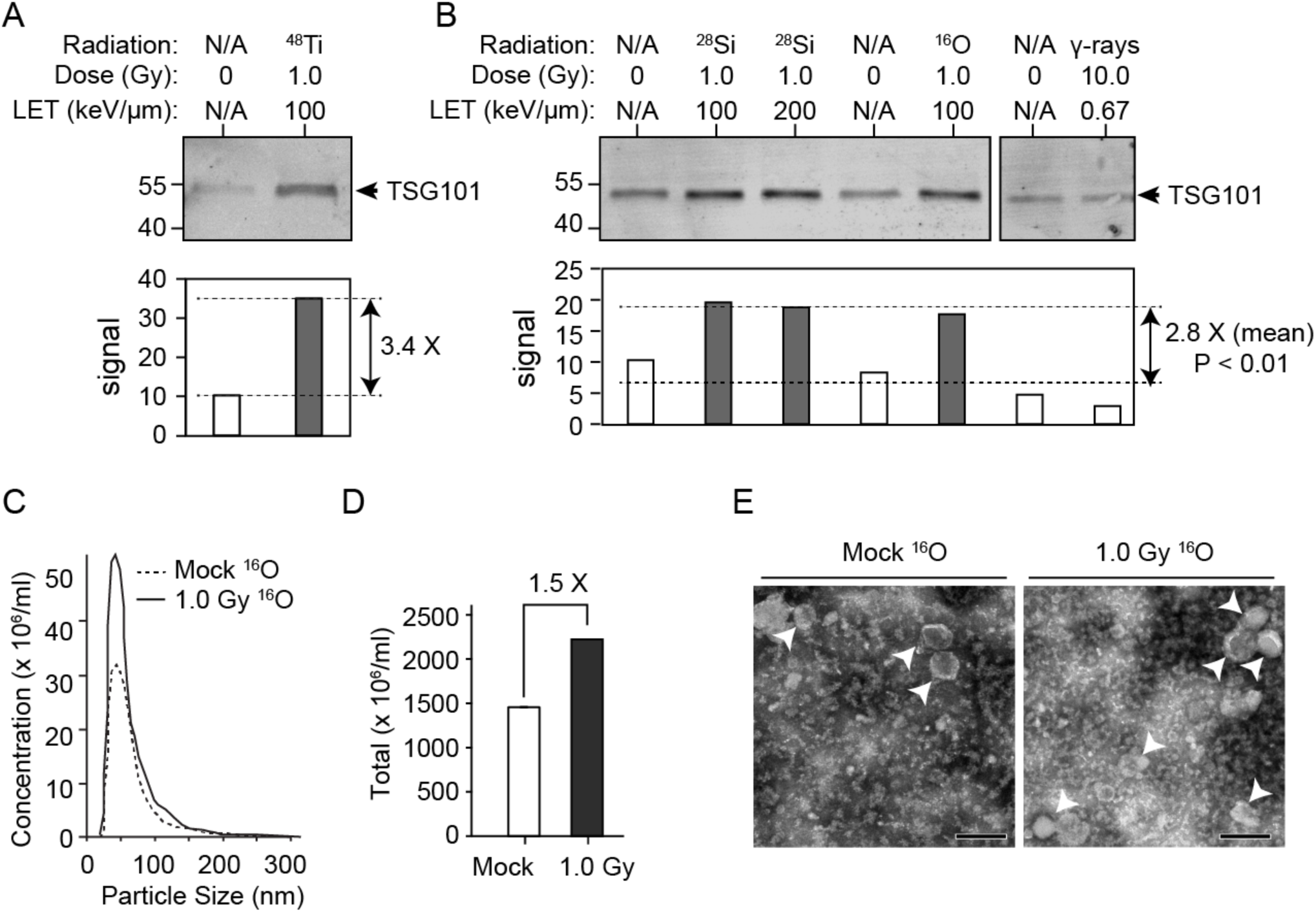
HZE ion irradiation stimulates exosome secretion. A. Effect of ^48^Ti ion irradiation on exosome release. Human bronchial epithelial cells (HBEC-3KT F25F) were irradiated at the indicated dose and LET values. Exosomes were prepared from conditioned medium as described in Materials and Methods and exosomal proteins were analyzed by immunoblotting using anti-TSG101 primary antibody and fluorescent secondary antibody. Imaging was performed using a LI-COR Odyssey system. Upper panel, immunoblot image with positions of molecular weight standards indicated at left (kDa) and TSG101 indicated by arrow at right. Lower panel, relative fluorescence signal intensity (arbitrary units). B. Effect of ^28^Si, ^16^O, and γ-ray irradiation on exosome release. Samples were prepared and analyzed as in Panel A. Upper panel, immunoblot image. Note that all lanes are from the same gel. Lower panel, relative fluorescence signal intensity, with dashed lines indicating mean intensity for HZE ion-irradiated groups and γ-ray/mock-irradiated groups. Statistical analysis was performed using a two-tailed Student’s t-test with P value indicated. C. Particle size distribution in the indicated extracellular vesicle preparations determined by NanoSight analysis. D. Quantification of results in Panel C. E. Transmission electron micrograph of indicated extracellular vesicle preparations. Arrowheads denote lipid-bounded vesicles that likely correspond to exosomes. Scale bar, 200 nm.

Selected samples were also analyzed by single particle tracking and electron microscopy. Consistent with the TSG101 immunoblotting results, single particle tracking analysis showed an increase in 30-100 nm vesicles in preparations derived from a ^16^O ion-irradiated culture relative to preparations from a mock-irradiated control culture (Fig. 1C, 1D). The peak of the vesicle size distribution was 44 nm, similar to what was seen in a prior study of irradiated keratinocytes (Jella et al., 2014). The result suggests that preparations were composed primarily of exosomes, rather than vesicles derived by budding from the plasma membrane or from apoptotic bodies, which are typically larger (Simpson and Mathivanan, 2012). Transmission electron microscopy confirmed the presence of numerous membrane-bounded vesicles in the exosome size range (Fig. 1E). We shall refer to preparations from HZE ion-irradiated cultures as “exosome-enriched,” as they are only partially purified, but contain exosomes as a major component.

### 3.2 Presence of damage-associated molecular patterns

We performed immunoblotting to investigate the presence of DAMPs in the extracellular vesicle preparations. The exosome-enriched preparations from HZE ion-irradiated cultures showed a significant, 3.9-fold increase in HSP70 levels, relative to mock-and γ-ray-irradiated cultures (Fig. 2A, P<0.01). Calreticulin levels also trended higher in the HZE ion-irradiated samples, although the difference was not significant. Results indicate that DAMPs are present in the vesicle preparations and that their levels increase roughly in proportion to TSG101.

**Fig. 2.**
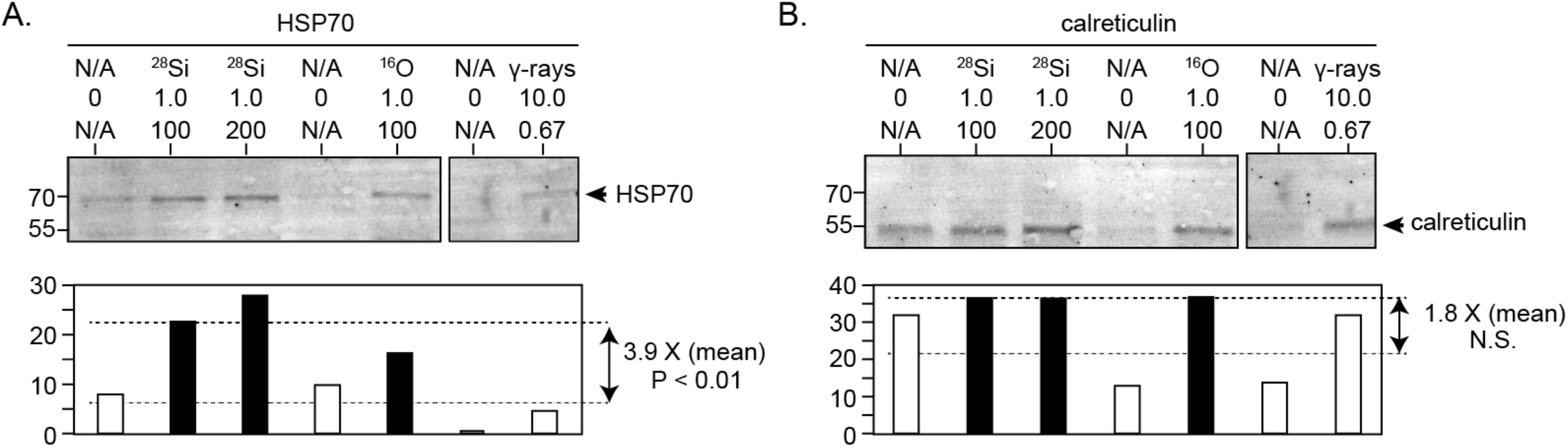
Presence of damage-associated molecular patterns in extracellular vesicle preparations. Immunoblotting analysis was performed using the same preparations as in Fig 1B. A. HSP70. Upper panel immunoblot image with molecular weight markers shown at left and arrow denoting HSP70 at right. Lower panels, quantification with dashed lines indicating mean intensity for HZE ion-irradiated groups and γ-ray/mock-irradiated groups. Statistical analysis was performed using a two-tailed Student’s t-test with P value indicated. B. calreticulin. Figure is labeled as in Panel A. N.S., not significant.

### 3.3 MiRNA profiling

We performed miRNA profiling to investigate the amount and types of miRNAs in the extracellular vesicle preparations. Small RNA was isolated as described in Materials and Methods. We obtained sufficient miRNA for analysis from ten independent samples, including four of the exosome-enriched HZE-irradiated samples, one γ-ray-irradiated sample, and five mock-irradiated control samples. Recovery of RNA was 2.0-fold greater in HZE-irradiated samples compared with the others, but the difference was not significant. Details of the RNA samples used are given in Table S1.

MiRNA profiling was performed using the NanoString platform, which is based on hybridization via bridge oligonucleotides to fluorescently tagged probes, with highly sensitive detection by single-molecule imaging. The platform provides results for 798 independent probes, representing 819 miRNA species (some probes hybridize to more than one closely related miRNA). Results are summarized in Fig. 3, and a full tabulation is presented in Table S2. The miRNA distribution reflects a few highly abundant species, together with a long tail of less abundant species (Fig. 3A). MiRNAs that have been particularly well studied in cancer are labeled. We note that just 10 of these miRNAs (miR-1246, miR-1290, miR-23a, miR-205, let-7b, miR-125b, miR-20a, miR-16, miR-21, and miR-221) together feature in more than five thousand cancer-related publications. MiRNAs of particular interest are further detailed in the Discussion.

**Fig. 3.**
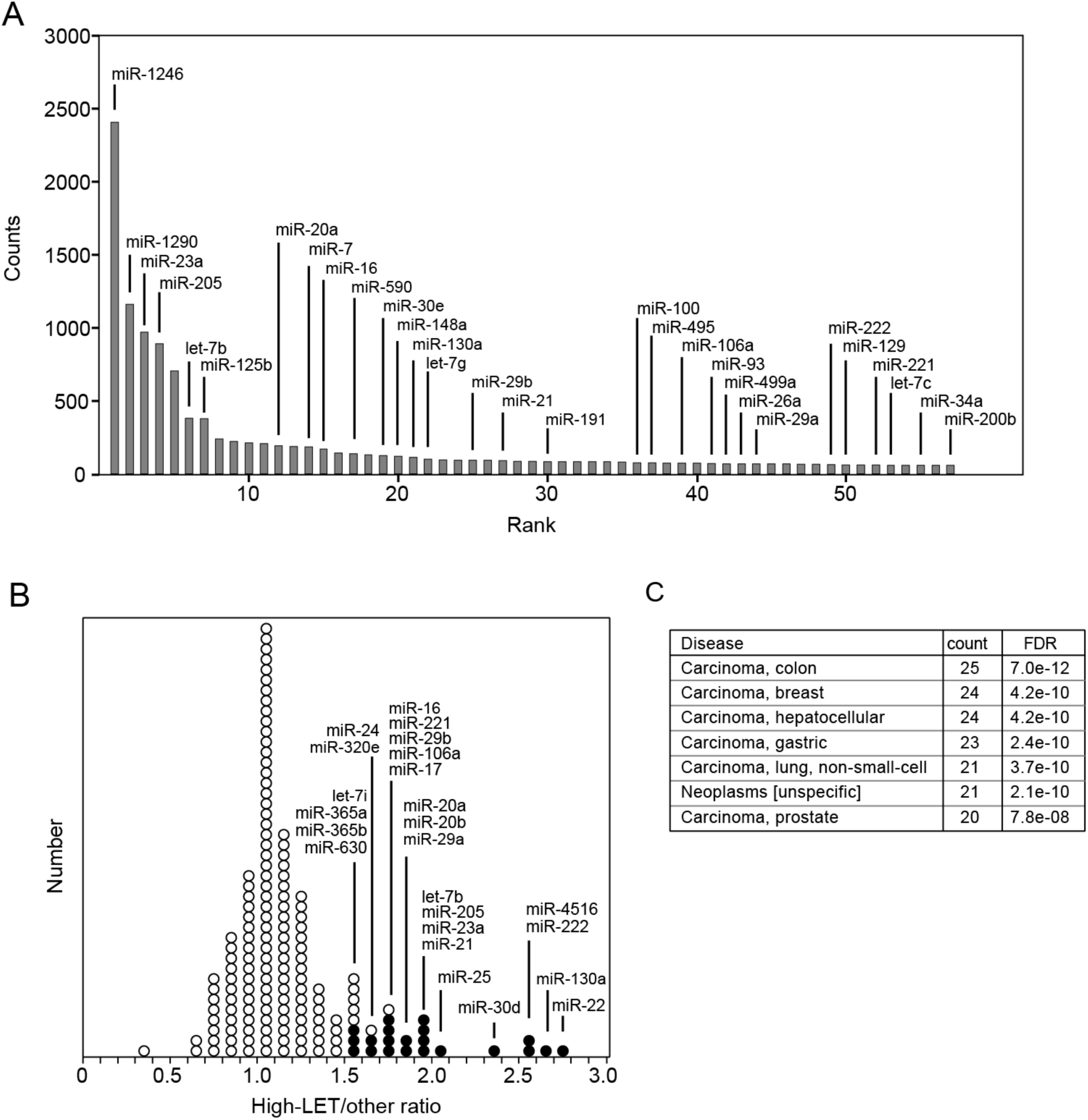
MicroRNA profiling. A. Average counts for miRNAs ranked in decreasing order of overall abundance. An arbitrary cutoff of three-fold over background was applied. Selected cancer-related miRNAs are labeled, based on a criterion of 100 or more PubMed citations retrieved using the name of the miRNA and the search term “cancer”, or 7 or more citations with the search term “lung cancer,” as of January, 2020. B. Histogram showing number of miRNAs with expression ratios in indicated ranges. Filled circles, probes that were modestly over-represented in HZE-irradiated samples based on expression ratio >1.5 and FDR <0.33 based on SAM analysis. Figure depicts the 160 most abundant miRNAs, with this cutoff chosen so as to include all of the miRNAs that met criteria as enriched in the HZE-irradiated samples. Labels indicate miRNAs corresponding to filled-circle probes (some probes hybridize to more than one closely related miRNA). C. Results of gene set enrichment analysis, using over-represented miRNAs from Fig 3B as a query set for the Tam 2.0 tool. Disease associations were ranked based on gene count and those with >20 counts are shown. FDR, false discovery rate as calculated by the TAM 2.0 tool.

We investigated whether any of the miRNAs were significantly over-represented in samples from HZE-irradiated cells using a Significance Analysis of Microarray (SAM) approach (Tusher et al., 2001), which takes into account the expression ratio, a measure of variance for individual probes, and a measure of variance for the data set as a whole. We used the SAM analysis, together with a 1.5-fold filter for expression ratio, to define a subset of 21 probes (corresponding to 24 miRNAs) that were modestly over-represented in the HZE-irradiated samples (Fig. 3B). We then used these miRNAs as a query set for gene set enrichment analysis using the TAM 2.0 tool (Li et al., 2018a). We ranked the results based on the number of miRNA genes in the query set that were related to a particular disease (noting that some miRNAs are encoded by more than one gene). Results showed a highly significant relationship between the query set and various forms of cancer, with FDRs ranging from 7.8 × 10^−8^ to 7.0 × 10^−12^ (Fig. 3C). There were also highly significant associations with cell death (FDR = 6.3 × 10^−14^) and with NF-κB regulation (FDR = 4.2 × 10^−8^). Complete results of the gene set enrichment analysis are presented in Table S3.

## 4. Discussion

Here we describe the ability of HZE ion irradiation to stimulate the release of extracellular vesicles from human bronchial epithelial cells, a cell type that is relevant to the development of lung cancer. To our knowledge, this is the first study of exosomes derived from high-LET irradiated cells. Results show that irradiation with HZE particles, but not reference γ-rays, stimulated exosome release by about 3-fold. Exosomes provide a means by which irradiated cells may communicate with their non-irradiated neighbors, which is of particular relevance at low fluences characteristic of space radiation environment, where only a fraction of cells may be directly traversed by HZE radiation tracks.

We characterized the exosome-enriched preparations for the presence of molecules that are known to modulate characteristics of the tissue microenvironment. Two DAMPs, HSP70 and calreticulin, were identified. The amount of these proteins correlated to the exosomal marker, TSG101. Prior work implicates HSP70 and calreticulin as modulators of immune and inflammatory responses (reviewed in (Boudesco et al., 2018; De Maio, 2014; Vandenabeele et al., 2016). Notably, HSP70 has been previously identified in exosomes derived from cancer cells exposed to genotoxic stress, where it activates cytokine production via an HSP70/toll-like receptor 2/NF-κB axis (Vulpis et al., 2017). The role of DAMPs in cancer has been compared to a “double-edged sword” (Hernandez et al., 2016). By triggering chronic inflammation, release of exosomal DAMPs may promote tumor development and progression. However, DAMPs may also alert the immune system to the presence of stressed or dying cells, promoting a host anti-tumor response.

Transport of miRNAs is another way in which exosomes influence their microenvironment. (reviewed in (Mills et al., 2019; Yu et al., 2016)). Exosomal miRNA transport is believed to be important for transmission of bystander effects (Jelonek et al., 2016; Xu et al., 2015), reviewed in (Du et al., 2020)). Stoichiometric analysis of exosomes prepared by standard methods suggests that only the most abundant species are likely to be present in amounts sufficient to influence gene expression in recipient cells (Chevillet et al., 2014). There were approximately a half dozen miRNAs that were detected at a much higher level than others in the exosome preparations. Among these, miR-1246 and miR-1290 have been shown to have convergent functions as drivers of nonsmall cell lung cancer initiation and progression (Zhang et al., 2016). Extracellular miR-1246, derived from irradiated lung cancer cells, promotes cancer cell proliferation and enhances radioresistance by direct targeting of the death receptor 5 gene (Yuan et al., 2016).

Two other miRNAs, miR-23a and miR-205, were abundant overall and also modestly over-represented in preparations from HZE-irradiated cells. MiR-23a, which has previously been identified in lung cancer cell-derived exosomes, is a proposed biomarker for early detection and adverse prognosis in lung cancer (Hetta et al., 2019; Qu et al., 2015). In addition, miR-23a from lung cancer-derived exosomes promotes endothelial cell proliferation and thus angiogenesis (Hsu et al., 2017) (Zheng et al., 2017). MiR-205 is pro-tumorigenic based on its ability to target expression of the PTEN tumor suppressor and other genes and is also a potential lung cancer biomarker (Bai et al., 2015; Wang et al., 2016).

One other miRNA, miR-7, is noteworthy because it has previously been detected in exosomes derived from irradiated bronchial epithelial cells and linked directly to propagation of radiation-induced bystander effect in recipient cells (Song et al., 2016). Autophagy induced by exosomal miR-7-5p was associated with EGFR/Akt/mTOR signaling pathway. The miR-7 signal ranked 14th overall among the 798 probes in the NanoString array.

There were 24 miRNAs that appeared to be modestly over-represented based on a SAM analysis, which takes into account both expression values and measures of uncertainty for those values. These included both the miR-23a and miR-205 discussed above and a number of somewhat less abundant species. Gene set enrichment analysis showed a highly significant association between the over-represented miRNAs and nonsmall cell lung and other cancers, consistent with the idea that exosomal miRNA could influence the risk of cancer development at the tissue level. Interestingly, four of the over-represented RNAs, let-7, miR-17, miR-24, and miR-21 are also part of a miRNA signature identified as a master regulator for spaceflight response, based on computational analysis of the NASA GeneLab database (Beheshti et al., 2018).

While this manuscript was in preparation, miRNA profiles were reported for exosomes prepared from low-LET irradiated cells and mice (Abramowicz et al., 2020; Zhang et al., 2020) Interestingly, several of the cancer related miRNAs described here, including miR-23a, miR-205, and let-7b, were relatively abundant in the cell-based exosome study, although their levels were unchanged following irradiation. Differences in the radiation type, radiation dose, and models make a more detailed comparison difficult.

The present studies establish a framework for understanding the potential role of exosomes in the GCR response. The limited amounts of material available for these studies did not allow for functional assays. It will be of interest in the future, however, to investigate whether exosome-enriched preparations from HZE-irradiated cells influence specific signaling pathways in bystander cells or whether they stimulate inflammatory responses at the tissue and organism level, as has been reported for exosomes derived from irradiated tumor cells (Diamond et al., 2018; Jella et al., 2020).

## ABBREVIATIONS

HZE: high charge and energy
LET: linear energy transfer
HSP70: heat shock protein, 70 kDa
DAMP: damage-associated molecular pattern
PBS: phosphate-buffered saline
miRNA: microRNA
SAM: Significance Analysis of Microarray
FDR: false discovery rate

## Acknowledgments

We thank the Emory University Robert P. Apkarian Integrated Electron Microscopy Core, the Emory Integrated Genomics Core, and the Morehouse School of Medicine Microvesicle Lab for invaluable technical services and advice. We thank the NASA Space Radiation Laboratory and Brookhaven National Laboratory staff for their support and assistance. This work was supported by grants from the US National Aeronautics and Space Administration, numbers NNX15AD63G and 80NSSC18K1116 (to WSD) and by the Winship Cancer Institute of Emory University. WSD is an Eminent Scholar of the Georgia Research Alliance.

## Appendix. Supplementary materials

Table S1. Characterization of small RNA preparations.

Table S2. NanoString nSolver export data.

Table S3. Results of TAM 2.0 gene set enrichment analysis.

