## Supplemental Tables 1, 2, and 3 for "Characterization of exosome release and extracellular vesicle-associated miRNAs for human bronchial epithelial cells irradiated with high charge and energy ions"

### TABLE LEGENDS

**Table S1. Characterization of small RNA preparations.** RNAs were prepared and characterized using an Agilent Bioanalyzer as described in the main text. NASA Space Radiation Laboratory campaign number, radiation type, calculated LET value, dose, RNA concentration, and percent miRNA are indicated. Mock irradiations were performed in parallel with the HZE ion irradiations indicated in parentheses. Two small RNA preparations that failed to pass quality control upon subsequent NanoString analysis are omitted.

**Table S2. Nanostring nSolver export data.** Table shows nSolver counts for indicated probes, ranked by mean value. Note that no normalization has been performed. Columns show rank; gene name (the name listed first in nSolver if more than one is listed); miRbase accession number; irradiation details (NASA Space Radiation Laboratory campaign, radiation type, LET value, and dose); mean counts for all samples, HZE ion-irradiated samples, and others; whether False Discovery Rate (q-value from SAM analysis) is less than 0.33; and names of additional cross-hybridizing RNAs. Two small RNA preparations that failed to pass quality control are omitted. Green shading denotes miRNAs that were modestly over-represented in HZE ion-irradiated groups, using criteria of q value <0.33 and >1.5-fold ratio of mean counts in HZE ion-irradiated groups versus others. Reference cellular mRNA probes, ligation control probes, negative control probes (with mean values indicated in bold), positive control probes, and non-human miRNA control probes (Arabidopsis thaliana, ath, Caenorhabditis elegans, cel, Oryza sativa, osa) are shown in rows 804 to 835 as indicated.

**Table S3. Results of TAM 2.0 gene set enrichment analysis.**

First page shows query set composed of miRNAs that are at least 1.5-fold over-represented in preparations from HZE-irradiated cultures and that have false discovery values <0.33 in SAM analysis. Refer to Table S2 for details. Black font indicates miRNA gene names listed first in nSolver output. Red font indicates additional miRNAs that cross-hybridize to the probe listed immediately above. Remaining pages indicate results of analysis run on indicated date. Results are ranked by number of miRNAs that meet a given criterion. Results for “Category:Disease” with count of 20 or more are also shown in Fig. 3C of the main text.

**Table S1. Characterization of small RNA preparations**

| NSRL campaign | Radiation type | LET (kev/µm) | Dose (Gy) | [RNA] (pg/µl) | miRNA (%) |
|---|---|---|---|---|---|
| 16A | 48-Ti | 200 | 1 | 1430 | 62 |
| 16C | 28-Si | 100 | 1 | 3015 | 77 |
| 16C | 28-Si | 200 | 1 | 879 | 44 |
| 16C | 16-O | 100 | 1 | 558 | 54 |
| 16C | gamma-ray | 0.67 | 10 | 418 | 51 |
| 16A | mock (48-Ti) | N/A | 0 | 982 | 59 |
| 16A | mock (48-Ti) | N/A | 0 | 1476 | 62 |
| 16C | mock (28-Si) | N/A | 0 | 405 | 46 |
| 16C | mock (16-O) | N/A | 0 | 601 | 59 |
| 16C | mock (16-O) | N/A | 0 | 533 | 64 |

**Table S2 nSolver export data**

| Rank | Gene Name | miRBase Accession | NSRL campaign | 16A | 16C | 16C | 16C | 16C | 16A | 16A | 16C | 16C | 16C | Mean | Mean((HZE) | Mean(other) | HZE:other | q<33% | Additional cross-hybridizing miRNAS | | | | |
|---|---|---|---|---|---|---|---|---|---|---|---|---|---|---|---|---|---|---|---|---|---|---|---|
| | | | Radiation type | 48-Ti | 28-Si | 28-Si | 16-O | gamma-ray | mock (48-Ti) | mock (48-Ti) | mock (28-Si) | mock (16-O) | mock (16-O) | | | | | | | | | | |
| | | | LET (keV/µm) | 200 | 100 | 200 | 100 | 0.67 | N/A | N/A | N/A | N/A | N/A | | | | | | | | | | |
| | | | Dose (Gy) | 1.0 | 1.0 | 1.0 | 1.0 | 10.0 | 0.0 | 0.0 | 0.0 | 0.0 | 0.0 | | | | | | Gene name 2 | Gene name 3 | Gene name 4 | Gene name 5 | Gene name 6 |
| 1 | hsa-miR-1246 | MIMAT0005898 | | 6775 | 1226 | 764 | 941 | 906 | 6965 | 4293 | 117 | 1817 | 273 | 2407.7 | 2427 | 2395 | 1.01 | | | | | | |
| 2 | hsa-miR-1290 | MIMAT0005880 | | 1995 | 957 | 983 | 419 | 1007 | 2330 | 2568 | 139 | 1010 | 208 | 1161.6 | 1089 | 1210 | 0.90 | | | | | | |
| 3 | hsa-miR-23a-3p | MIMAT0000078 | | 1390 | 1194 | 2522 | 379 | 312 | 1512 | 1391 | 217 | 690 | 98 | 970.5 | 1371 | 703 | 1.95 | yes | | | | | |
| 4 | hsa-miR-205-5p | MIMAT0000266 | | 1738 | 794 | 2020 | 465 | 267 | 1701 | 922 | 273 | 629 | 98 | 890.7 | 1254 | 648 | 1.93 | yes | | | | | |
| 5 | hsa-miR-4454 | MIMAT0018976 | | 1390 | 645 | 981 | 207 | 537 | 1468 | 983 | 90 | 646 | 90 | 703.7 | 806 | 636 | 1.27 | | hsa-miR-7975 | | | | |
| 6 | hsa-let-7b-5p | MIMAT0000063 | | 566 | 339 | 1081 | 149 | 184 | 585 | 463 | 107 | 264 | 61 | 379.9 | 534 | 277 | 1.92 | yes | | | | | |
| 7 | hsa-miR-125b-5p | MIMAT0000423 | | 902 | 282 | 577 | 140 | 177 | 834 | 513 | 76 | 240 | 40 | 378.1 | 475 | 313 | 1.52 | | | | | | |
| 8 | hsa-miR-644a | MIMAT0003314 | | 147 | 300 | 434 | 187 | 391 | 194 | 176 | 105 | 356 | 99 | 238.9 | 267 | 220 | 1.21 | | | | | | |
| 9 | hsa-miR-1253 | MIMAT0005904 | | 153 | 244 | 400 | 198 | 446 | 207 | 58 | 83 | 324 | 109 | 222.2 | 249 | 205 | 1.22 | | | | | | |
| 10 | hsa-miR-548n | MIMAT0005916 | | 131 | 155 | 227 | 201 | 414 | 139 | 108 | 130 | 482 | 127 | 211.4 | 179 | 233 | 0.77 | | | | | | |
| 11 | hsa-miR-549a | MIMAT0003333 | | 181 | 232 | 317 | 156 | 369 | 169 | 181 | 86 | 286 | 94 | 207.1 | 222 | 198 | 1.12 | | | | | | |
| 12 | hsa-miR-20a-5p | MIMAT0000075 | | 384 | 163 | 422 | 85 | 77 | 357 | 206 | 47 | 145 | 36 | 192.2 | 264 | 145 | 1.82 | yes | hsa-miR-20b-5p | | | | |
| 13 | hsa-miR-548aa | MIMAT0018447 | | 129 | 185 | 263 | 182 | 398 | 138 | 58 | 83 | 349 | 90 | 187.5 | 190 | 186 | 1.02 | | hsa-miR-548t-3p | | | | |
| 14 | hsa-miR-7-5p | MIMAT0000252 | | 412 | 161 | 211 | 45 | 80 | 389 | 383 | 25 | 97 | 28 | 183.1 | 207 | 167 | 1.24 | | | | | | |
| 15 | hsa-miR-16-5p | MIMAT0000069 | | 326 | 143 | 344 | 85 | 115 | 261 | 116 | 92 | 189 | 11 | 168.2 | 225 | 131 | 1.72 | yes | | | | | |
| 16 | hsa-miR-2116-5p | MIMAT0011160 | | 93 | 146 | 330 | 105 | 247 | 94 | 119 | 52 | 160 | 69 | 141.5 | 169 | 124 | 1.36 | | | | | | |
| 17 | hsa-miR-590-5p | MIMAT0003258 | | 119 | 146 | 185 | 125 | 190 | 143 | 121 | 62 | 211 | 60 | 136.2 | 144 | 131 | 1.10 | | | | | | |
| 18 | hsa-miR-520d-3p | MIMAT0002856 | | 58 | 105 | 118 | 126 | 336 | 69 | 31 | 55 | 286 | 104 | 128.8 | 102 | 147 | 0.69 | | | | | | |
| 19 | hsa-miR-30e-5p | MIMAT0000692 | | 97 | 132 | 214 | 108 | 205 | 85 | 91 | 54 | 196 | 66 | 124.8 | 138 | 116 | 1.19 | | | | | | |
| 20 | hsa-miR-148a-3p | MIMAT0000243 | | 243 | 126 | 162 | 44 | 67 | 286 | 149 | 20 | 83 | 23 | 120.3 | 144 | 105 | 1.37 | | | | | | |
| 21 | hsa-miR-130a-3p | MIMAT0000425 | | 153 | 152 | 380 | 35 | 37 | 150 | 97 | 37 | 65 | 21 | 112.7 | 180 | 68 | 2.65 | yes | | | | | |
| 22 | hsa-let-7g-5p | MIMAT0000414 | | 153 | 110 | 199 | 37 | 33 | 191 | 151 | 29 | 83 | 18 | 100.4 | 125 | 84 | 1.48 | | | | | | |
| 23 | hsa-miR-548q | MIMAT0011163 | | 83 | 88 | 142 | 81 | 151 | 90 | 67 | 45 | 165 | 35 | 94.7 | 99 | 92 | 1.07 | | | | | | |
| 24 | hsa-miR-3161 | MIMAT0015035 | | 62 | 76 | 126 | 107 | 145 | 89 | 74 | 40 | 147 | 64 | 93 | 93 | 93 | 1.00 | | | | | | |
| 25 | hsa-miR-29b-3p | MIMAT0000100 | | 116 | 112 | 232 | 40 | 63 | 137 | 108 | 24 | 74 | 16 | 92.2 | 125 | 70 | 1.78 | yes | | | | | |
| 26 | hsa-miR-365a-3p | MIMAT0000710 | | 123 | 113 | 177 | 51 | 61 | 140 | 87 | 41 | 100 | 26 | 91.9 | 116 | 76 | 1.53 | yes | hsa-miR-365b-3p | | | | |
| 27 | hsa-miR-21-5p | MIMAT0000076 | | 139 | 71 | 256 | 43 | 52 | 151 | 66 | 37 | 62 | 21 | 89.8 | 127 | 65 | 1.96 | yes | | | | | |
| 28 | hsa-miR-186-5p | MIMAT0000456 | | 78 | 67 | 142 | 70 | 137 | 84 | 67 | 39 | 116 | 45 | 84.5 | 89 | 81 | 1.10 | | | | | | |
| 29 | hsa-miR-4531 | MIMAT0019070 | | 42 | 67 | 104 | 83 | 182 | 71 | 31 | 40 | 173 | 48 | 84.1 | 74 | 91 | 0.81 | | | | | | |
| 30 | hsa-miR-191-5p | MIMAT0000440 | | 145 | 80 | 156 | 42 | 28 | 169 | 111 | 21 | 70 | 15 | 83.7 | 106 | 69 | 1.53 | | | | | | |
| 31 | hsa-miR-4455 | MIMAT0018977 | | 61 | 74 | 101 | 73 | 150 | 65 | 52 | 45 | 171 | 45 | 83.7 | 77 | 88 | 0.88 | | | | | | |
| 32 | hsa-miR-4536-5p | MIMAT0019078 | | 65 | 99 | 125 | 72 | 122 | 65 | 75 | 37 | 121 | 47 | 82.8 | 90 | 78 | 1.16 | | | | | | |
| 33 | hsa-miR-548g-3p | MIMAT0005912 | | 72 | 91 | 132 | 75 | 126 | 66 | 58 | 47 | 116 | 41 | 82.4 | 93 | 76 | 1.22 | | | | | | |
| 34 | hsa-miR-4516 | MIMAT0019053 | | 43 | 289 | 156 | 23 | 17 | 64 | 48 | 22 | 134 | 19 | 81.5 | 128 | 51 | 2.52 | yes | | | | | |
| 35 | hsa-miR-378h | MIMAT0018984 | | 54 | 83 | 131 | 76 | 116 | 81 | 78 | 43 | 98 | 40 | 80 | 86 | 76 | 1.13 | | | | | | |
| 36 | hsa-miR-100-5p | MIMAT0000098 | | 111 | 104 | 124 | 24 | 52 | 109 | 115 | 17 | 72 | 30 | 75.8 | 91 | 66 | 1.38 | | | | | | |
| 37 | hsa-miR-495-3p | MIMAT0002817 | | 63 | 79 | 96 | 65 | 108 | 63 | 85 | 41 | 116 | 40 | 75.6 | 76 | 76 | 1.00 | | | | | | |
| 38 | hsa-miR-627-5p | MIMAT0003296 | | 1 | 13 | 46 | 90 | 201 | 21 | 1 | 52 | 240 | 68 | 73.3 | 38 | 97 | 0.39 | | | | | | |
| 39 | hsa-miR-106a-5p | MIMAT0000103 | | 110 | 94 | 161 | 27 | 30 | 136 | 70 | 16 | 68 | 14 | 72.6 | 98 | 56 | 1.76 | yes | hsa-miR-17-5p | | | | |
| 40 | hsa-miR-320d | MIMAT0006764 | | 58 | 67 | 106 | 81 | 100 | 61 | 76 | 27 | 105 | 38 | 71.9 | 78 | 68 | 1.15 | | | | | | |
| 41 | hsa-miR-93-5p | MIMAT0000093 | | 134 | 59 | 125 | 39 | 32 | 118 | 77 | 28 | 70 | 21 | 70.3 | 89 | 58 | 1.55 | | | | | | |
| 42 | hsa-miR-499a-5p | MIMAT0002870 | | 62 | 77 | 92 | 51 | 72 | 93 | 65 | 42 | 70 | 55 | 67.9 | 71 | 66 | 1.07 | | | | | | |
| 43 | hsa-miR-26a-5p | MIMAT0000082 | | 79 | 74 | 147 | 34 | 75 | 105 | 57 | 28 | 56 | 21 | 67.6 | 84 | 57 | 1.46 | | | | | | |
| 44 | hsa-miR-29a-3p | MIMAT0000086 | | 74 | 56 | 215 | 32 | 45 | 57 | 47 | 62 | 65 | 22 | 67.5 | 94 | 50 | 1.90 | yes | | | | | |
| 45 | hsa-miR-1297 | MIMAT0005886 | | 52 | 77 | 85 | 65 | 108 | 57 | 47 | 31 | 113 | 37 | 67.2 | 70 | 66 | 1.06 | | | | | | |
| 46 | hsa-miR-548ah-5p | MIMAT0018972 | | 58 | 61 | 88 | 67 | 82 | 72 | 64 | 36 | 87 | 53 | 66.8 | 69 | 66 | 1.04 | | | | | | |
| 47 | hsa-miR-548ai | MIMAT0018989 | | 39 | 50 | 67 | 60 | 131 | 39 | 31 | 41 | 153 | 47 | 65.8 | 54 | 74 | 0.73 | | hsa-miR-570-5p | | | | |
| 48 | hsa-miR-363-3p | MIMAT0000707 | | 81 | 61 | 55 | 22 | 103 | 71 | 40 | 23 | 104 | 94 | 65.4 | 55 | 73 | 0.76 | | | | | | |
| 49 | hsa-miR-222-3p | MIMAT0000279 | | 57 | 47 | 270 | 24 | 43 | 48 | 40 | 38 | 48 | 13 | 62.8 | 100 | 38 | 2.60 | yes | | | | | |
| 50 | hsa-miR-129-2-3p | MIMAT0004605 | | 110 | 41 | 61 | 39 | 35 | 135 | 89 | 16 | 57 | 26 | 60.9 | 63 | 60 | 1.05 | | | | | | |
| 51 | hsa-let-7i-5p | MIMAT0000415 | | 91 | 61 | 122 | 34 | 51 | 69 | 65 | 21 | 64 | 26 | 60.4 | 77 | 49 | 1.56 | yes | | | | | |
| 52 | hsa-miR-221-3p | MIMAT0000278 | | 93 | 78 | 107 | 42 | 36 | 78 | 51 | 37 | 65 | 14 | 60.1 | 80 | 47 | 1.71 | yes | | | | | |
| 53 | hsa-let-7c-5p | MIMAT0000064 | | 96 | 45 | 110 | 27 | 35 | 99 | 76 | 20 | 51 | 19 | 57.8 | 70 | 50 | 1.39 | | | | | | |
| 54 | hsa-miR-888-5p | MIMAT0004916 | | 42 | 52 | 114 | 42 | 90 | 39 | 45 | 37 | 72 | 42 | 57.5 | 63 | 54 | 1.15 | | | | | | |
| 55 | hsa-miR-34a-5p | MIMAT0000255 | | 105 | 29 | 109 | 39 | 37 | 102 | 51 | 25 | 59 | 18 | 57.4 | 71 | 49 | 1.45 | | | | | | |
| 56 | hsa-miR-320e | MIMAT0015072 | | 86 | 87 | 101 | 27 | 43 | 67 | 56 | 18 | 64 | 21 | 57 | 75 | 45 | 1.68 | yes | | | | | |
| 57 | hsa-miR-200b-3p | MIMAT0000318 | | 123 | 48 | 88 | 26 | 29 | 117 | 67 | 15 | 39 | 17 | 56.9 | 71 | 47 | 1.51 | | | | | | |
| 58 | hsa-miR-155-5p | MIMAT0000646 | | 46 | 55 | 82 | 42 | 70 | 61 | 39 | 22 | 97 | 29 | 54.3 | 56 | 53 | 1.06 | | | | | | |
| 59 | hsa-miR-379-5p | MIMAT0000733 | | 50 | 52 | 60 | 47 | 76 | 58 | 55 | 26 | 88 | 28 | 54 | 52 | 55 | 0.95 | | | | | | |
| 60 | hsa-miR-25-3p | MIMAT0000081 | | 81 | 70 | 129 | 30 | 24 | 57 | 68 | 28 | 34 | 17 | 53.8 | 78 | 38 | 2.04 | yes | | | | | |
| 61 | hsa-miR-551a | MIMAT0003214 | | 50 | 51 | 37 | 39 | 72 | 59 | 35 | 33 | 81 | 73 | 53 | 44 | 59 | 0.75 | | | | | | |
| 62 | hsa-miR-183-5p | MIMAT0000261 | | 44 | 50 | 81 | 44 | 59 | 51 | 48 | 32 | 56 | 37 | 50.2 | 55 | 47 | 1.16 | | | | | | |
| 63 | hsa-miR-1305 | MIMAT0005893 | | 43 | 57 | 74 | 42 | 65 | 42 | 68 | 30 | 52 | 26 | 49.9 | 54 | 47 | 1.14 | | | | | | |
| 64 | hsa-miR-200c-3p | MIMAT0000617 | | 70 | 48 | 103 | 18 | 41 | 82 | 54 | 13 | 30 | 23 | 48.2 | 60 | 41 | 1.48 | | | | | | |
| 65 | hsa-miR-107 | MIMAT0000104 | | 38 | 55 | 76 | 31 | 57 | 39 | 50 | 24 | 52 | 57 | 47.9 | 50 | 47 | 1.08 | | | | | | |
| 66 | hsa-miR-548d-3p | MIMAT0003323 | | 31 | 37 | 80 | 51 | 73 | 43 | 31 | 25 | 75 | 29 | 47.5 | 50 | 46 | 1.08 | | | | | | |
| 67 | hsa-miR-30d-5p | MIMAT0000245 | | 54 | 149 | 56 | 30 | 22 | 37 | 26 | 15 | 57 | 25 | 47.1 | 72 | 30 | 2.38 | yes | | | | | |
| 68 | hsa-miR-4286 | MIMAT0016916 | | 81 | 31 | 47 | 25 | 28 | 111 | 60 | 22 | 49 | 13 | 46.7 | 46 | 47 | 0.98 | | | | | | |
| 69 | hsa-miR-548z | MIMAT0018446 | | 30 | 43 | 52 | 34 | 73 | 34 | 40 | 31 | 88 | 34 | 45.9 | 40 | 50 | 0.80 | | hsa-miR-548h-3p | | | | |
| 70 | hsa-miR-612 | MIMAT0003280 | | 40 | 39 | 59 | 43 | 64 | 45 | 48 | 25 | 61 | 26 | 45 | 45 | 45 | 1.01 | | | | | | |
| 71 | hsa-miR-520h | MIMAT0002867 | | 49 | 51 | 63 | 38 | 61 | 40 | 27 | 21 | 56 | 26 | 43.2 | 50 | 39 | 1.31 | | | | | | |
| 72 | hsa-miR-200a-3p | MIMAT0000682 | | 43 | 46 | 54 | 38 | 33 | 58 | 54 | 24 | 47 | 31 | 42.8 | 45 | 41 | 1.10 | | | | | | |
| 73 | hsa-miR-514a-3p | MIMAT0002883 | | 36 | 43 | 41 | 35 | 58 | 30 | 42 | 27 | 77 | 39 | 42.8 | 39 | 46 | 0.85 | | | | | | |
| 74 | hsa-miR-548y | MIMAT0018354 | | 38 | 39 | 62 | 39 | 52 | 36 | 47 | 20 | 66 | 24 | 42.3 | 45 | 41 | 1.09 | | | | | | |
| 75 | hsa-miR-1270 | MIMAT0005924 | | 54 | 41 | 56 | 22 | 60 | 41 | 33 | 34 | 48 | 33 | 42.2 | 43 | 42 | 1.04 | | | | | | |
| 76 | hsa-miR-1257 | MIMAT0005908 | | 38 | 42 | 61 | 33 | 54 | 42 | 39 | 23 | 60 | 29 | 42.1 | 44 | 41 | 1.06 | | | | | | |
| 77 | hsa-miR-32-5p | MIMAT0000090 | | 18 | 51 | 71 | 30 | 71 | 27 | 29 | 30 | 72 | 21 | 42 | 43 | 42 | 1.02 | | | | | | |
| 78 | hsa-miR-219b-3p | MIMAT0019748 | | 30 | 37 | 48 | 41 | 59 | 37 | 42 | 23 | 60 | 42 | 41.9 | 39 | 44 | 0.89 | | | | | | |
| 79 | hsa-miR-944 | MIMAT0004987 | | 30 | 41 | 68 | 32 | 78 | 37 | 36 | 17 | 56 | 24 | 41.9 | 43 | 41 | 1.03 | | | | | | |
| 80 | hsa-miR-26b-5p | MIMAT0000083 | | 45 | 31 | 81 | 29 | 34 | 44 | 45 | 31 | 55 | 20 | 41.5 | 47 | 38 | 1.22 | | | | | | |
| 81 | hsa-miR-630 | MIMAT0003299 | | 69 | 62 | 52 | 30 | 36 | 52 | 27 | 21 | 51 | 14 | 41.4 | 53 | 34 | 1.59 | yes | | | | | |
| 82 | hsa-miR-19b-3p | MIMAT0000074 | | 73 | 40 | 80 | 24 | 22 | 71 | 29 | 19 | 44 | 11 | 41.3 | 54 | 33 | 1.66 | | | | | | |
| 83 | hsa-miR-132-3p | MIMAT0000426 | | 69 | 36 | 46 | 18 | 15 | 100 | 58 | 15 | 29 | 26 | 41.2 | 42 | 41 | 1.04 | | | | | | |
| 84 | hsa-miR-873-3p | MIMAT0022717 | | 37 | 43 | 39 | 26 | 66 | 49 | 25 | 8 | 63 | 56 | 41.2 | 36 | 45 | 0.81 | | | | | | |
| 85 | hsa-miR-411-5p | MIMAT0003329 | | 45 | 49 | 41 | 34 | 63 | 57 | 44 | 16 | 50 | 11 | 41 | 42 | 40 | 1.05 | | | | | | |
| 86 | hsa-miR-212-3p | MIMAT0000269 | | 30 | 49 | 43 | 30 | 51 | 38 | 44 | 28 | 55 | 41 | 40.9 | 38 | 43 | 0.89 | | | | | | |
| 87 | hsa-miR-519b-5p | MIMAT0005454 | | 31 | 29 | 41 | 37 | 82 | 31 | 21 | 22 | 88 | 27 | 40.9 | 35 | 45 | 0.76 | | hsa-miR-519c-5p | hsa-miR-523-5p | hsa-miR-518e-5p | hsa-miR-522-5p | hsa-miR-519a-5p |
| 88 | hsa-miR-423-5p | MIMAT0004748 | | 39 | 40 | 100 | 27 | 24 | 56 | 30 | 18 | 46 | 28 | 40.8 | 52 | 34 | 1.53 | | | | | | |
| 89 | hsa-miR-577 | MIMAT0003242 | | 32 | 45 | 42 | 45 | 44 | 45 | 37 | 21 | 54 | 42 | 40.7 | 41 | 41 | 1.01 | | | | | | |
| 90 | hsa-miR-627-3p | MIMAT0026623 | | 45 | 31 | 49 | 39 | 39 | 37 | 46 | 24 | 60 | 36 | 40.6 | 41 | 40 | 1.02 | | | | | | |

| | | | | | | | | | | | | | | | | | | | | |
|---|---|---|---|---|---|---|---|---|---|---|---|---|---|---|---|---|---|---|---|---|
| 91 | hsa-miR-1255a | MIMAT0005906 | | 36 | 46 | 52 | 35 | 53 | 36 | 38 | 19 | 54 | 25 | 39.4 | 42 | 38 | 1.13 | | | |
| 92 | hsa-miR-1197 | MIMAT0005955 | | 31 | 40 | 53 | 22 | 55 | 28 | 51 | 27 | 45 | 40 | 39.2 | 37 | 41 | 0.89 | | | |
| 93 | hsa-miR-598-3p | MIMAT0003266 | | 43 | 39 | 36 | 32 | 47 | 37 | 32 | 22 | 52 | 51 | 39.1 | 38 | 40 | 0.93 | | | |
| 94 | hsa-miR-1-5p | MIMAT0031892 | | 34 | 53 | 51 | 35 | 55 | 37 | 33 | 21 | 53 | 18 | 39 | 43 | 36 | 1.20 | | | |
| 95 | hsa-miR-1285-5p | MIMAT0022719 | | 30 | 37 | 56 | 30 | 66 | 48 | 44 | 21 | 41 | 15 | 38.8 | 38 | 39 | 0.98 | | | |
| 96 | hsa-miR-378f | MIMAT0018932 | | 32 | 65 | 44 | 24 | 55 | 29 | 38 | 19 | 47 | 33 | 38.6 | 41 | 37 | 1.12 | | | |
| 97 | hsa-miR-548v | MIMAT0015020 | | 30 | 33 | 59 | 35 | 55 | 33 | 26 | 19 | 57 | 39 | 38.6 | 39 | 38 | 1.03 | | | |
| 98 | hsa-miR-149-5p | MIMAT0000450 | | 45 | 32 | 49 | 30 | 49 | 30 | 42 | 21 | 51 | 35 | 38.4 | 39 | 38 | 1.03 | | | |
| 99 | hsa-miR-99a-5p | MIMAT0000097 | | 62 | 46 | 57 | 17 | 28 | 73 | 48 | 8 | 35 | 10 | 38.4 | 46 | 34 | 1.35 | | | |
| 100 | hsa-miR-376a-3p | MIMAT0000729 | | 41 | 41 | 49 | 41 | 34 | 51 | 35 | 20 | 58 | 13 | 38.3 | 43 | 35 | 1.22 | | | |
| 101 | hsa-miR-574-5p | MIMAT0004795 | | 41 | 38 | 48 | 28 | 49 | 41 | 36 | 18 | 56 | 22 | 37.7 | 39 | 37 | 1.05 | | | |
| 102 | hsa-miR-1245b-5p | MIMAT0019950 | | 22 | 44 | 31 | 28 | 55 | 37 | 37 | 28 | 56 | 32 | 37 | 31 | 41 | 0.77 | | | |
| 103 | hsa-miR-548e-5p | MIMAT0026736 | | 33 | 37 | 41 | 25 | 51 | 30 | 33 | 30 | 61 | 29 | 37 | 34 | 39 | 0.87 | | | |
| 104 | hsa-miR-219a-2-3p | MIMAT0004675 | | 36 | 50 | 43 | 34 | 55 | 32 | 28 | 19 | 45 | 26 | 36.8 | 41 | 34 | 1.19 | | | |
| 105 | hsa-miR-324-3p | MIMAT0000762 | | 37 | 40 | 55 | 30 | 39 | 33 | 42 | 14 | 53 | 23 | 36.6 | 41 | 34 | 1.19 | | | |
| 106 | hsa-miR-181a-5p | MIMAT0000256 | | 52 | 38 | 92 | 15 | 25 | 54 | 39 | 13 | 24 | 12 | 36.4 | 49 | 28 | 1.77 | | | |
| 107 | hsa-miR-10b-5p | MIMAT0000254 | | 28 | 42 | 45 | 21 | 49 | 32 | 49 | 22 | 46 | 28 | 36.2 | 34 | 38 | 0.90 | | | |
| 108 | hsa-miR-335-5p | MIMAT0000765 | | 51 | 33 | 31 | 21 | 32 | 81 | 53 | 16 | 27 | 16 | 36.1 | 34 | 38 | 0.91 | | | |
| 109 | hsa-miR-651-5p | MIMAT0003321 | | 39 | 39 | 50 | 26 | 40 | 34 | 34 | 19 | 49 | 31 | 36.1 | 39 | 35 | 1.12 | | | |
| 110 | hsa-miR-374a-5p | MIMAT0000727 | | 32 | 32 | 56 | 25 | 42 | 40 | 42 | 18 | 50 | 21 | 35.8 | 36 | 36 | 1.02 | | | |
| 111 | hsa-miR-10a-5p | MIMAT0000253 | | 25 | 39 | 46 | 32 | 46 | 39 | 34 | 26 | 46 | 24 | 35.7 | 36 | 36 | 0.99 | | | |
| 112 | hsa-miR-6721-5p | MIMAT0025852 | | 38 | 34 | 23 | 8 | 61 | 36 | 28 | 18 | 58 | 53 | 35.7 | 26 | 42 | 0.61 | | | |
| 113 | hsa-miR-542-5p | MIMAT0003340 | | 52 | 44 | 51 | 14 | 29 | 56 | 36 | 17 | 32 | 24 | 35.5 | 40 | 32 | 1.24 | | | |
| 114 | hsa-miR-24-3p | MIMAT0000080 | | 51 | 41 | 72 | 19 | 30 | 37 | 27 | 20 | 29 | 28 | 35.4 | 46 | 29 | 1.61 | yes | | |
| 115 | hsa-miR-342-3p | MIMAT0000753 | | 52 | 33 | 60 | 13 | 25 | 53 | 36 | 15 | 37 | 29 | 35.3 | 40 | 33 | 1.22 | | | |
| 116 | hsa-miR-6724-5p | MIMAT0025856 | | 58 | 33 | 39 | 17 | 39 | 55 | 37 | 19 | 36 | 20 | 35.3 | 37 | 34 | 1.07 | | | |
| 117 | hsa-let-7a-5p | MIMAT0000062 | | 67 | 12 | 78 | 1 | 1 | 114 | 73 | 1 | 1 | 1 | 34.9 | 40 | 32 | 1.24 | | | |
| 118 | hsa-miR-216b-5p | MIMAT0004959 | | 31 | 34 | 45 | 33 | 53 | 41 | 28 | 22 | 47 | 15 | 34.9 | 36 | 34 | 1.04 | | | |
| 119 | hsa-miR-197-3p | MIMAT0000227 | | 56 | 40 | 48 | 19 | 29 | 48 | 32 | 15 | 37 | 23 | 34.7 | 41 | 31 | 1.33 | | | |
| 120 | hsa-miR-607 | MIMAT0003275 | | 22 | 29 | 58 | 33 | 38 | 32 | 41 | 32 | 39 | 23 | 34.7 | 36 | 34 | 1.04 | | | |
| 121 | hsa-miR-1268a | MIMAT0005922 | | 75 | 30 | 33 | 22 | 25 | 86 | 30 | 6 | 32 | 7 | 34.6 | 40 | 31 | 1.29 | | | |
| 122 | hsa-miR-299-5p | MIMAT0002890 | | 31 | 30 | 48 | 25 | 42 | 31 | 43 | 29 | 45 | 21 | 34.5 | 34 | 35 | 0.95 | | | |
| 123 | hsa-miR-503-5p | MIMAT0002874 | | 37 | 30 | 39 | 25 | 40 | 37 | 55 | 26 | 33 | 23 | 34.5 | 33 | 36 | 0.92 | | | |
| 124 | hsa-miR-376c-3p | MIMAT0000720 | | 74 | 27 | 34 | 14 | 23 | 92 | 33 | 10 | 20 | 17 | 34.4 | 37 | 33 | 1.15 | | | |
| 125 | hsa-miR-301a-5p | MIMAT0022696 | | 24 | 38 | 39 | 30 | 41 | 31 | 42 | 27 | 50 | 20 | 34.2 | 33 | 35 | 0.93 | | | |
| 126 | hsa-miR-597-5p | MIMAT0003265 | | 19 | 33 | 55 | 29 | 58 | 31 | 25 | 16 | 48 | 26 | 34 | 34 | 34 | 1.00 | | | |
| 127 | hsa-miR-378g | MIMAT0018937 | | 21 | 44 | 46 | 30 | 33 | 35 | 40 | 31 | 32 | 27 | 33.9 | 35 | 33 | 1.07 | | | |
| 128 | hsa-miR-23c | MIMAT0018000 | | 36 | 32 | 32 | 31 | 51 | 33 | 32 | 17 | 41 | 33 | 33.8 | 33 | 35 | 0.95 | | | |
| 129 | hsa-miR-127-3p | MIMAT0000446 | | 34 | 33 | 42 | 24 | 36 | 30 | 33 | 23 | 52 | 30 | 33.7 | 33 | 34 | 0.98 | | | |
| 130 | hsa-miR-1183 | MIMAT0005828 | | 35 | 36 | 37 | 16 | 50 | 42 | 15 | 12 | 45 | 47 | 33.5 | 31 | 35 | 0.88 | | | |
| 131 | hsa-miR-208b-3p | MIMAT0004960 | | 27 | 45 | 38 | 30 | 44 | 30 | 35 | 20 | 33 | 32 | 33.4 | 35 | 32 | 1.08 | | | |
| 132 | hsa-miR-525-5p | MIMAT0002838 | | 20 | 41 | 42 | 31 | 50 | 25 | 33 | 24 | 40 | 28 | 33.4 | 34 | 33 | 1.01 | | | |
| 133 | hsa-miR-371a-5p | MIMAT0004687 | | 27 | 20 | 45 | 22 | 41 | 41 | 32 | 19 | 55 | 31 | 33.3 | 29 | 37 | 0.78 | | | |
| 134 | hsa-miR-765 | MIMAT0003945 | | 33 | 28 | 34 | 35 | 50 | 24 | 35 | 28 | 41 | 25 | 33.3 | 33 | 34 | 0.96 | | | |
| 135 | hsa-miR-23b-3p | MIMAT0000418 | | 45 | 31 | 58 | 20 | 21 | 55 | 40 | 18 | 26 | 18 | 33.2 | 39 | 30 | 1.30 | | | |
| 136 | hsa-miR-346 | MIMAT0000773 | | 42 | 32 | 49 | 27 | 40 | 26 | 33 | 17 | 38 | 28 | 33.2 | 38 | 30 | 1.24 | | | |
| 137 | hsa-miR-196a-5p | MIMAT0000226 | | 22 | 42 | 46 | 34 | 42 | 17 | 33 | 14 | 60 | 20 | 33 | 36 | 31 | 1.16 | | | |
| 138 | hsa-miR-4707-5p | MIMAT0019807 | | 28 | 35 | 39 | 26 | 43 | 28 | 41 | 20 | 40 | 29 | 32.9 | 32 | 34 | 0.96 | | | |
| 139 | hsa-miR-661 | MIMAT0003324 | | 19 | 28 | 57 | 19 | 47 | 23 | 38 | 22 | 51 | 25 | 32.9 | 31 | 34 | 0.90 | | | |
| 140 | hsa-miR-648 | MIMAT0003318 | | 35 | 45 | 43 | 18 | 38 | 32 | 29 | 23 | 38 | 26 | 32.7 | 35 | 31 | 1.14 | | | |
| 141 | hsa-miR-451a | MIMAT0001631 | | 37 | 44 | 33 | 17 | 38 | 41 | 47 | 18 | 35 | 16 | 32.6 | 33 | 33 | 1.01 | | | |
| 142 | hsa-miR-585-3p | MIMAT0003250 | | 21 | 34 | 42 | 36 | 55 | 27 | 27 | 19 | 48 | 17 | 32.6 | 33 | 32 | 1.03 | | | |
| 143 | hsa-miR-548ar-3p | MIMAT0022266 | | 28 | 33 | 38 | 31 | 55 | 33 | 22 | 14 | 55 | 16 | 32.5 | 33 | 33 | 1.00 | | | |
| 144 | hsa-miR-548al | MIMAT0019024 | | 39 | 24 | 33 | 26 | 46 | 33 | 14 | 17 | 55 | 37 | 32.4 | 31 | 34 | 0.91 | | | |
| 145 | hsa-miR-1268b | MIMAT0018925 | | 30 | 35 | 33 | 33 | 31 | 36 | 31 | 18 | 42 | 32 | 32.1 | 33 | 32 | 1.03 | | | |
| 146 | hsa-miR-563 | MIMAT0003227 | | 33 | 28 | 35 | 32 | 48 | 32 | 28 | 16 | 37 | 32 | 32.1 | 32 | 32 | 0.99 | | | |
| 147 | hsa-miR-567 | MIMAT0003231 | | 36 | 39 | 38 | 33 | 48 | 23 | 26 | 16 | 41 | 20 | 32 | 37 | 29 | 1.26 | | | |
| 148 | hsa-miR-587 | MIMAT0003253 | | 28 | 36 | 51 | 31 | 32 | 38 | 23 | 17 | 41 | 23 | 32 | 37 | 29 | 1.26 | | | |
| 149 | hsa-miR-522-3p | MIMAT0002868 | | 20 | 30 | 49 | 30 | 39 | 28 | 29 | 25 | 48 | 19 | 31.7 | 32 | 31 | 1.03 | | | |
| 150 | hsa-miR-764 | MIMAT0010367 | | 25 | 31 | 56 | 25 | 28 | 23 | 35 | 16 | 47 | 31 | 31.7 | 34 | 30 | 1.14 | | | |
| 151 | hsa-miR-770-5p | MIMAT0003948 | | 39 | 35 | 48 | 25 | 30 | 30 | 26 | 17 | 52 | 15 | 31.7 | 37 | 28 | 1.30 | | | |
| 152 | hsa-miR-1295a | MIMAT0005885 | | 30 | 40 | 39 | 30 | 40 | 23 | 37 | 15 | 35 | 25 | 31.4 | 35 | 29 | 1.19 | | | |
| 153 | hsa-miR-922 | MIMAT0004972 | | 22 | 44 | 33 | 24 | 34 | 34 | 24 | 28 | 37 | 33 | 31.3 | 31 | 32 | 0.97 | | | |
| 154 | hsa-miR-22-3p | MIMAT0000077 | | 40 | 31 | 119 | 11 | 11 | 29 | 17 | 18 | 30 | 6 | 31.2 | 50 | 19 | 2.72 | yes | | |
| 155 | hsa-miR-553 | MIMAT0003216 | | 30 | 36 | 37 | 26 | 30 | 34 | 38 | 18 | 33 | 30 | 31.2 | 32 | 31 | 1.06 | | | |
| 156 | hsa-miR-3136-5p | MIMAT0015003 | | 30 | 32 | 35 | 37 | 42 | 27 | 29 | 15 | 40 | 24 | 31.1 | 34 | 30 | 1.14 | | | |
| 157 | hsa-miR-548j-3p | MIMAT0026737 | | 27 | 16 | 41 | 30 | 49 | 24 | 25 | 16 | 54 | 29 | 31.1 | 29 | 33 | 0.87 | | | |
| 158 | hsa-miR-508-3p | MIMAT0002880 | | 28 | 30 | 45 | 26 | 43 | 36 | 25 | 27 | 32 | 17 | 30.9 | 32 | 30 | 1.08 | | | |
| 159 | hsa-miR-584-3p | MIMAT0022708 | | 31 | 33 | 34 | 37 | 39 | 24 | 23 | 23 | 45 | 19 | 30.8 | 34 | 29 | 1.17 | | | |
| 160 | hsa-miR-450b-5p | MIMAT0004909 | | 14 | 34 | 59 | 24 | 37 | 29 | 24 | 18 | 41 | 26 | 30.6 | 33 | 29 | 1.12 | | | |
| 161 | hsa-miR-656-3p | MIMAT0003332 | | 25 | 36 | 47 | 27 | 25 | 29 | 32 | 16 | 41 | 28 | 30.6 | 34 | 29 | 1.18 | | | |
| 162 | hsa-miR-517a-3p | MIMAT0002852 | | 32 | 40 | 43 | 19 | 38 | 16 | 27 | 22 | 37 | 31 | 30.5 | 34 | 29 | 1.18 | | | |
| 163 | hsa-let-7f-5p | MIMAT0000067 | | 32 | 26 | 45 | 11 | 39 | 54 | 28 | 11 | 40 | 16 | 30.2 | 29 | 31 | 0.91 | | | |
| 164 | hsa-miR-224-5p | MIMAT0000281 | | 18 | 32 | 31 | 30 | 42 | 28 | 40 | 12 | 44 | 25 | 30.2 | 28 | 32 | 0.87 | | | |
| 165 | hsa-miR-325 | MIMAT0000771 | | 28 | 39 | 36 | 20 | 31 | 32 | 29 | 20 | 47 | 20 | 30.2 | 31 | 30 | 1.03 | | | |
| 166 | hsa-miR-340-5p | MIMAT0004692 | | 26 | 40 | 38 | 21 | 34 | 34 | 22 | 19 | 50 | 18 | 30.2 | 31 | 30 | 1.06 | | | |
| 167 | hsa-miR-526a | MIMAT0002845 | | 34 | 26 | 35 | 31 | 41 | 22 | 32 | 23 | 34 | 21 | 29.9 | 32 | 29 | 1.09 | | hsa-miR-518c-5p | hsa-miR-518d-5p |
| 168 | hsa-miR-28-5p | MIMAT0000085 | | 30 | 32 | 53 | 23 | 33 | 25 | 32 | 16 | 36 | 17 | 29.7 | 35 | 27 | 1.30 | | | |
| 169 | hsa-miR-301b-3p | MIMAT0004958 | | 17 | 40 | 34 | 30 | 46 | 33 | 22 | 19 | 35 | 21 | 29.7 | 30 | 29 | 1.03 | | | |
| 170 | hsa-miR-873-5p | MIMAT0004953 | | 23 | 34 | 43 | 38 | 49 | 19 | 16 | 11 | 40 | 24 | 29.7 | 35 | 27 | 1.30 | | | |
| 171 | hsa-miR-877-5p | MIMAT0004949 | | 31 | 27 | 30 | 29 | 42 | 28 | 22 | 17 | 48 | 23 | 29.7 | 29 | 30 | 0.98 | | | |
| 172 | hsa-miR-30a-5p | MIMAT0000087 | | 37 | 42 | 36 | 22 | 37 | 29 | 17 | 12 | 46 | 18 | 29.6 | 34 | 27 | 1.29 | | | |
| 173 | hsa-miR-1296-3p | MIMAT0026637 | | 27 | 31 | 40 | 23 | 30 | 23 | 42 | 13 | 43 | 23 | 29.5 | 30 | 29 | 1.04 | | | |
| 174 | hsa-miR-3147 | MIMAT0015019 | | 24 | 32 | 39 | 28 | 38 | 23 | 28 | 16 | 38 | 29 | 29.5 | 31 | 29 | 1.07 | | | |
| 175 | hsa-miR-514b-5p | MIMAT0015087 | | 26 | 27 | 29 | 28 | 41 | 31 | 25 | 21 | 41 | 26 | 29.5 | 28 | 31 | 0.89 | | | |
| 176 | hsa-miR-203a-3p | MIMAT0000264 | | 30 | 41 | 45 | 15 | 33 | 33 | 27 | 21 | 34 | 15 | 29.4 | 33 | 27 | 1.21 | | | |
| 177 | hsa-miR-761 | MIMAT0010364 | | 27 | 41 | 17 | 18 | 39 | 23 | 26 | 10 | 53 | 39 | 29.3 | 26 | 32 | 0.81 | | | |
| 178 | hsa-miR-543 | MIMAT0004954 | | 21 | 25 | 34 | 21 | 46 | 26 | 31 | 22 | 45 | 21 | 29.2 | 25 | 32 | 0.79 | | | |
| 179 | hsa-miR-423-3p | MIMAT0001340 | | 30 | 33 | 43 | 24 | 43 | 31 | 24 | 6 | 35 | 22 | 29.1 | 33 | 27 | 1.21 | | | |
| 180 | hsa-miR-548a-5p | MIMAT0004803 | | 31 | 21 | 36 | 27 | 40 | 29 | 19 | 18 | 47 | 23 | 29.1 | 29 | 29 | 0.98 | | | |
| 181 | hsa-miR-561-5p | MIMAT0022706 | | 20 | 24 | 27 | 39 | 42 | 22 | 18 | 21 | 59 | 19 | 29.1 | 28 | 30 | 0.91 | | | |
| 182 | hsa-miR-92a-3p | MIMAT0000092 | | 31 | 14 | 54 | 17 | 29 | 40 | 36 | 18 | 36 | 16 | 29.1 | 29 | 29 | 0.99 | | | |
| 183 | hsa-miR-2117 | MIMAT0011162 | | 31 | 32 | 47 | 28 | 31 | 22 | 18 | 25 | 34 | 21 | 28.9 | 35 | 25 | 1.37 | | | |
| 184 | hsa-miR-95-3p | MIMAT0000094 | | 22 | 26 | 32 | 27 | 38 | 31 | 33 | 19 | 33 | 28 | 28.9 | 27 | 30 | 0.88 | | | |
| 185 | hsa-miR-151a-3p | MIMAT0000757 | | 39 | 33 | 41 | 20 | 27 | 36 | 30 | 13 | 31 | 18 | 28.8 | 33 | 26 | 1.29 | | | |

| | | | | | | | | | | | | | | | | | | | |
|---|---|---|---|---|---|---|---|---|---|---|---|---|---|---|---|---|---|---|---|
| 186 | hsa-miR-520a-5p | MIMAT0002833 | | 30 | 24 | 33 | 29 | 42 | 21 | 29 | 22 | 38 | 19 | 28.7 | 29 | 29 | 1.02 | | |
| 187 | hsa-miR-30c-5p | MIMAT0000244 | | 35 | 27 | 37 | 28 | 29 | 40 | 30 | 11 | 34 | 15 | 28.6 | 32 | 27 | 1.20 | | |
| 188 | hsa-miR-643 | MIMAT0003313 | | 24 | 27 | 55 | 19 | 30 | 24 | 31 | 18 | 39 | 19 | 28.6 | 31 | 27 | 1.16 | | |
| 189 | hsa-miR-148b-3p | MIMAT0000759 | | 22 | 30 | 38 | 24 | 32 | 36 | 30 | 10 | 47 | 15 | 28.4 | 29 | 28 | 1.01 | | |
| 190 | hsa-miR-199a-3p | MIMAT0000232 | | 29 | 33 | 47 | 23 | 31 | 20 | 32 | 13 | 38 | 17 | 28.3 | 33 | 25 | 1.31 | | hsa-miR-199b-3p |
| 191 | hsa-miR-1258 | MIMAT0005909 | | 33 | 27 | 28 | 25 | 34 | 30 | 22 | 20 | 39 | 20 | 27.8 | 28 | 28 | 1.03 | | |
| 192 | hsa-miR-3144-5p | MIMAT0015014 | | 15 | 25 | 36 | 33 | 49 | 23 | 17 | 21 | 44 | 15 | 27.8 | 27 | 28 | 0.97 | | |
| 193 | hsa-miR-3131 | MIMAT0014996 | | 21 | 26 | 42 | 19 | 38 | 24 | 30 | 24 | 40 | 13 | 27.7 | 27 | 28 | 0.96 | | |
| 194 | hsa-miR-548j-5p | MIMAT0005875 | | 22 | 31 | 33 | 19 | 43 | 28 | 24 | 23 | 41 | 13 | 27.7 | 26 | 29 | 0.92 | | |
| 195 | hsa-miR-744-5p | MIMAT0004945 | | 29 | 25 | 41 | 19 | 31 | 34 | 28 | 17 | 29 | 24 | 27.7 | 29 | 27 | 1.05 | | |
| 196 | hsa-miR-1827 | MIMAT0006767 | | 17 | 22 | 42 | 24 | 35 | 37 | 34 | 11 | 30 | 24 | 27.6 | 26 | 29 | 0.92 | | |
| 197 | hsa-miR-887-5p | MIMAT0026720 | | 27 | 30 | 38 | 20 | 25 | 40 | 18 | 19 | 30 | 29 | 27.6 | 29 | 27 | 1.07 | | |
| 198 | hsa-miR-542-3p | MIMAT0003389 | | 21 | 23 | 36 | 19 | 37 | 23 | 28 | 18 | 45 | 25 | 27.5 | 25 | 29 | 0.84 | | |
| 199 | hsa-miR-758-3p | MIMAT0003879 | | 35 | 33 | 19 | 26 | 29 | 23 | 24 | 18 | 35 | 32 | 27.4 | 28 | 27 | 1.05 | | hsa-miR-411-3p |
| 200 | hsa-miR-1972 | MIMAT0009447 | | 17 | 22 | 29 | 25 | 28 | 34 | 38 | 14 | 39 | 27 | 27.3 | 23 | 30 | 0.78 | | |
| 201 | hsa-miR-499a-3p | MIMAT0004772 | | 24 | 26 | 31 | 22 | 32 | 19 | 17 | 16 | 65 | 21 | 27.3 | 26 | 28 | 0.91 | | |
| 202 | hsa-miR-595 | MIMAT0003263 | | 23 | 25 | 34 | 24 | 38 | 19 | 27 | 31 | 29 | 23 | 27.3 | 27 | 28 | 0.95 | | |
| 203 | hsa-miR-193a-5p | MIMAT0004614 | | 31 | 21 | 71 | 18 | 26 | 25 | 23 | 22 | 23 | 12 | 27.2 | 35 | 22 | 1.61 | | hsa-miR-193b-5p |
| 204 | hsa-miR-361-5p | MIMAT0000703 | | 33 | 32 | 56 | 17 | 22 | 38 | 24 | 14 | 26 | 9 | 27.1 | 35 | 22 | 1.56 | | |
| 205 | hsa-miR-566 | MIMAT0003230 | | 23 | 31 | 26 | 13 | 38 | 29 | 28 | 17 | 41 | 25 | 27.1 | 23 | 30 | 0.78 | | |
| 206 | hsa-miR-605-5p | MIMAT0003273 | | 37 | 29 | 30 | 18 | 36 | 24 | 17 | 14 | 41 | 24 | 27 | 29 | 26 | 1.10 | | |
| 207 | hsa-miR-1236-3p | MIMAT0005591 | | 21 | 30 | 29 | 16 | 37 | 31 | 28 | 14 | 36 | 27 | 26.9 | 24 | 29 | 0.83 | | |
| 208 | hsa-miR-125a-5p | MIMAT0000443 | | 62 | 27 | 32 | 17 | 25 | 40 | 25 | 8 | 24 | 9 | 26.9 | 35 | 22 | 1.58 | | |
| 209 | hsa-miR-15b-5p | MIMAT0000417 | | 41 | 26 | 44 | 12 | 38 | 32 | 31 | 13 | 24 | 8 | 26.9 | 31 | 24 | 1.26 | | |
| 210 | hsa-miR-421 | MIMAT0003339 | | 22 | 26 | 42 | 21 | 33 | 21 | 29 | 19 | 42 | 14 | 26.9 | 28 | 26 | 1.05 | | |
| 211 | hsa-miR-1323 | MIMAT0005795 | | 23 | 23 | 36 | 20 | 35 | 32 | 29 | 18 | 25 | 27 | 26.8 | 26 | 28 | 0.92 | | |
| 212 | hsa-miR-146b-5p | MIMAT0002809 | | 31 | 28 | 32 | 19 | 24 | 23 | 36 | 15 | 39 | 21 | 26.8 | 28 | 26 | 1.04 | | |
| 213 | hsa-miR-299-3p | MIMAT0000687 | | 25 | 28 | 42 | 17 | 27 | 28 | 26 | 23 | 29 | 23 | 26.8 | 28 | 26 | 1.08 | | |
| 214 | hsa-miR-1306-3p | MIMAT0005950 | | 25 | 34 | 36 | 19 | 38 | 27 | 22 | 17 | 24 | 24 | 26.6 | 29 | 25 | 1.13 | | |
| 215 | hsa-miR-378d | MIMAT0018926 | | 22 | 29 | 29 | 22 | 27 | 23 | 39 | 19 | 34 | 22 | 26.6 | 26 | 27 | 0.93 | | |
| 216 | hsa-miR-449b-5p | MIMAT0003327 | | 21 | 24 | 30 | 30 | 34 | 20 | 33 | 19 | 38 | 17 | 26.6 | 26 | 27 | 0.98 | | |
| 217 | hsa-miR-19a-3p | MIMAT0000073 | | 37 | 31 | 49 | 16 | 19 | 38 | 25 | 6 | 33 | 10 | 26.4 | 33 | 22 | 1.52 | | |
| 218 | hsa-miR-27b-3p | MIMAT0000419 | | 38 | 30 | 42 | 18 | 19 | 29 | 32 | 16 | 26 | 14 | 26.4 | 32 | 23 | 1.41 | | |
| 219 | hsa-miR-302b-3p | MIMAT0000715 | | 16 | 27 | 34 | 24 | 48 | 15 | 27 | 17 | 44 | 12 | 26.4 | 25 | 27 | 0.93 | | |
| 220 | hsa-miR-302a-5p | MIMAT0000683 | | 26 | 38 | 33 | 21 | 36 | 27 | 16 | 10 | 36 | 20 | 26.3 | 30 | 24 | 1.22 | | |
| 221 | hsa-miR-4443 | MIMAT0018961 | | 24 | 25 | 30 | 12 | 33 | 35 | 19 | 11 | 39 | 35 | 26.3 | 23 | 29 | 0.79 | | |
| 222 | hsa-miR-98-5p | MIMAT0000096 | | 33 | 29 | 40 | 18 | 26 | 40 | 26 | 13 | 23 | 15 | 26.3 | 30 | 24 | 1.26 | | |
| 223 | hsa-let-7d-5p | MIMAT0000065 | | 32 | 29 | 55 | 12 | 28 | 40 | 17 | 15 | 23 | 11 | 26.2 | 32 | 22 | 1.43 | | |
| 224 | hsa-miR-1262 | MIMAT0005914 | | 18 | 21 | 30 | 22 | 35 | 27 | 40 | 14 | 38 | 17 | 26.2 | 23 | 29 | 0.80 | | |
| 225 | hsa-miR-506-5p | MIMAT0022701 | | 27 | 32 | 37 | 30 | 22 | 28 | 28 | 10 | 26 | 22 | 26.2 | 32 | 23 | 1.39 | | |
| 226 | hsa-miR-891b | MIMAT0004913 | | 32 | 21 | 27 | 16 | 42 | 22 | 23 | 16 | 40 | 22 | 26.1 | 24 | 28 | 0.87 | | |
| 227 | hsa-miR-141-3p | MIMAT0000432 | | 31 | 13 | 48 | 23 | 28 | 34 | 20 | 16 | 27 | 20 | 26 | 29 | 24 | 1.19 | | |
| 228 | hsa-miR-615-3p | MIMAT0003283 | | 16 | 27 | 32 | 25 | 34 | 31 | 25 | 12 | 37 | 21 | 26 | 25 | 27 | 0.94 | | |
| 229 | hsa-miR-219a-5p | MIMAT0000276 | | 21 | 31 | 19 | 24 | 28 | 33 | 20 | 19 | 38 | 26 | 25.9 | 24 | 27 | 0.87 | | |
| 230 | hsa-miR-1272 | MIMAT0005925 | | 19 | 24 | 32 | 21 | 38 | 28 | 21 | 18 | 35 | 22 | 25.8 | 24 | 27 | 0.89 | | |
| 231 | hsa-miR-375 | MIMAT0000728 | | 44 | 22 | 24 | 22 | 27 | 45 | 23 | 11 | 25 | 15 | 25.8 | 28 | 24 | 1.15 | | |
| 232 | hsa-miR-378i | MIMAT0019074 | | 28 | 29 | 29 | 17 | 25 | 19 | 28 | 17 | 48 | 18 | 25.8 | 26 | 26 | 1.00 | | |
| 233 | hsa-miR-548a-3p | MIMAT0003251 | | 20 | 26 | 33 | 24 | 39 | 21 | 28 | 17 | 36 | 14 | 25.8 | 26 | 26 | 1.00 | | |
| 234 | hsa-miR-301a-3p | MIMAT0000688 | | 26 | 23 | 43 | 24 | 28 | 21 | 32 | 13 | 30 | 17 | 25.7 | 29 | 24 | 1.23 | | |
| 235 | hsa-miR-518b | MIMAT0002844 | | 31 | 24 | 25 | 23 | 31 | 31 | 24 | 11 | 30 | 27 | 25.7 | 26 | 26 | 1.00 | | |
| 236 | hsa-miR-766-3p | MIMAT0003888 | | 29 | 42 | 19 | 23 | 43 | 17 | 25 | 14 | 25 | 20 | 25.7 | 28 | 24 | 1.18 | | |
| 237 | hsa-miR-99b-5p | MIMAT0000689 | | 26 | 18 | 39 | 22 | 24 | 35 | 27 | 14 | 28 | 23 | 25.6 | 26 | 25 | 1.04 | | |
| 238 | hsa-miR-377-3p | MIMAT0000730 | | 24 | 26 | 31 | 21 | 43 | 23 | 25 | 17 | 26 | 19 | 25.5 | 26 | 26 | 1.00 | | |
| 239 | hsa-miR-412-3p | MIMAT0002170 | | 12 | 27 | 31 | 13 | 27 | 22 | 27 | 17 | 49 | 30 | 25.5 | 21 | 29 | 0.72 | | |
| 240 | hsa-miR-548i | MIMAT0005935 | | 18 | 29 | 25 | 27 | 34 | 23 | 28 | 14 | 33 | 24 | 25.5 | 25 | 26 | 0.95 | | |
| 241 | hsa-miR-1254 | MIMAT0005905 | | 21 | 19 | 27 | 26 | 22 | 33 | 29 | 17 | 39 | 21 | 25.4 | 23 | 27 | 0.87 | | |
| 242 | hsa-miR-2053 | MIMAT0009978 | | 25 | 20 | 34 | 19 | 33 | 24 | 31 | 13 | 30 | 25 | 25.4 | 25 | 26 | 0.94 | | |
| 243 | hsa-miR-514b-3p | MIMAT0015088 | | 22 | 22 | 50 | 28 | 32 | 13 | 29 | 14 | 26 | 17 | 25.3 | 31 | 22 | 1.40 | | |
| 244 | hsa-miR-483-3p | MIMAT0002173 | | 23 | 28 | 40 | 20 | 29 | 30 | 21 | 14 | 29 | 17 | 25.1 | 28 | 23 | 1.19 | | |
| 245 | hsa-miR-370-3p | MIMAT0000722 | | 24 | 25 | 37 | 26 | 38 | 25 | 24 | 17 | 26 | 8 | 25 | 28 | 23 | 1.22 | | |
| 246 | hsa-miR-142-3p | MIMAT0000434 | | 29 | 27 | 25 | 21 | 27 | 26 | 13 | 13 | 29 | 39 | 24.9 | 26 | 25 | 1.04 | | |
| 247 | hsa-miR-502-3p | MIMAT0004775 | | 23 | 34 | 31 | 18 | 32 | 18 | 24 | 14 | 41 | 14 | 24.9 | 27 | 24 | 1.11 | | |
| 248 | hsa-miR-1261 | MIMAT0005913 | | 22 | 24 | 27 | 19 | 40 | 22 | 18 | 13 | 44 | 19 | 24.8 | 23 | 26 | 0.88 | | |
| 249 | hsa-miR-337-3p | MIMAT0000754 | | 18 | 25 | 27 | 22 | 32 | 32 | 19 | 22 | 32 | 18 | 24.7 | 23 | 26 | 0.89 | | |
| 250 | hsa-miR-574-3p | MIMAT0003239 | | 31 | 23 | 44 | 17 | 16 | 31 | 21 | 14 | 28 | 21 | 24.6 | 29 | 22 | 1.32 | | |
| 251 | hsa-miR-1279 | MIMAT0005937 | | 16 | 23 | 30 | 22 | 30 | 21 | 24 | 15 | 36 | 27 | 24.4 | 23 | 26 | 0.89 | | |
| 252 | hsa-miR-27a-3p | MIMAT0000084 | | 15 | 26 | 36 | 15 | 31 | 18 | 24 | 24 | 34 | 21 | 24.4 | 23 | 25 | 0.91 | | |
| 253 | hsa-miR-3180 | MIMAT0018178 | | 20 | 24 | 29 | 23 | 25 | 20 | 36 | 17 | 32 | 18 | 24.4 | 24 | 25 | 0.97 | | |
| 254 | hsa-miR-4461 | MIMAT0018983 | | 22 | 21 | 27 | 29 | 32 | 13 | 29 | 17 | 41 | 13 | 24.4 | 25 | 24 | 1.02 | | |
| 255 | hsa-miR-532-3p | MIMAT0004780 | | 15 | 18 | 30 | 29 | 25 | 27 | 28 | 16 | 37 | 19 | 24.4 | 23 | 25 | 0.91 | | |
| 256 | hsa-miR-552-3p | MIMAT0003215 | | 20 | 35 | 22 | 15 | 27 | 29 | 25 | 14 | 33 | 24 | 24.4 | 23 | 25 | 0.91 | | |
| 257 | hsa-miR-582-3p | MIMAT0004797 | | 19 | 29 | 34 | 25 | 25 | 28 | 14 | 16 | 31 | 21 | 24.2 | 27 | 23 | 1.19 | | |
| 258 | hsa-miR-206 | MIMAT0000462 | | 33 | 26 | 28 | 24 | 26 | 26 | 13 | 14 | 31 | 20 | 24.1 | 28 | 22 | 1.28 | | |
| 259 | hsa-miR-380-3p | MIMAT0000735 | | 20 | 21 | 31 | 23 | 31 | 24 | 26 | 12 | 30 | 23 | 24.1 | 24 | 24 | 0.98 | | |
| 260 | hsa-miR-518f-3p | MIMAT0002842 | | 24 | 18 | 38 | 17 | 36 | 26 | 10 | 14 | 38 | 20 | 24.1 | 24 | 24 | 1.01 | | |
| 261 | hsa-miR-548ak | MIMAT0019013 | | 20 | 27 | 30 | 23 | 30 | 18 | 20 | 15 | 43 | 15 | 24.1 | 25 | 24 | 1.06 | | |
| 262 | hsa-miR-608 | MIMAT0003276 | | 22 | 19 | 36 | 22 | 25 | 25 | 28 | 8 | 38 | 18 | 24.1 | 25 | 24 | 1.05 | | |
| 263 | hsa-miR-128-1-5p | MIMAT0026477 | | 16 | 21 | 25 | 26 | 28 | 25 | 23 | 20 | 39 | 17 | 24 | 22 | 25 | 0.87 | | |
| 264 | hsa-miR-511-5p | MIMAT0002808 | | 27 | 30 | 29 | 20 | 30 | 25 | 20 | 13 | 29 | 17 | 24 | 27 | 22 | 1.19 | | |
| 265 | hsa-miR-197-5p | MIMAT0022691 | | 18 | 30 | 30 | 17 | 34 | 26 | 24 | 12 | 26 | 22 | 23.9 | 24 | 24 | 0.99 | | |
| 266 | hsa-miR-302a-3p | MIMAT0000684 | | 12 | 27 | 31 | 19 | 32 | 18 | 23 | 13 | 42 | 22 | 23.9 | 22 | 25 | 0.89 | | |
| 267 | hsa-miR-199b-5p | MIMAT0000263 | | 16 | 22 | 39 | 28 | 23 | 24 | 30 | 17 | 24 | 15 | 23.8 | 26 | 22 | 1.18 | | |
| 268 | hsa-miR-424-5p | MIMAT0001341 | | 26 | 26 | 42 | 8 | 23 | 23 | 26 | 8 | 38 | 18 | 23.8 | 26 | 23 | 1.13 | | |
| 269 | hsa-miR-454-3p | MIMAT0003885 | | 21 | 21 | 26 | 23 | 35 | 21 | 15 | 21 | 35 | 20 | 23.8 | 23 | 25 | 0.93 | | |
| 270 | hsa-miR-936 | MIMAT0004979 | | 21 | 21 | 28 | 17 | 32 | 27 | 25 | 12 | 34 | 21 | 23.8 | 22 | 25 | 0.86 | | |
| 271 | hsa-miR-3127-5p | MIMAT0014990 | | 19 | 27 | 27 | 17 | 34 | 24 | 22 | 12 | 38 | 17 | 23.7 | 23 | 25 | 0.92 | | |
| 272 | hsa-miR-498 | MIMAT0002824 | | 23 | 26 | 31 | 25 | 26 | 20 | 18 | 12 | 41 | 14 | 23.6 | 26 | 22 | 1.20 | | |
| 273 | hsa-miR-450a-5p | MIMAT0001545 | | 23 | 24 | 45 | 18 | 16 | 22 | 28 | 19 | 19 | 21 | 23.5 | 28 | 21 | 1.32 | | |
| 274 | hsa-miR-664b-3p | MIMAT0022272 | | 24 | 26 | 34 | 21 | 30 | 20 | 17 | 24 | 23 | 15 | 23.4 | 26 | 22 | 1.22 | | |
| 275 | hsa-miR-1271-5p | MIMAT0005796 | | 20 | 31 | 25 | 15 | 25 | 26 | 24 | 15 | 32 | 20 | 23.3 | 23 | 24 | 0.96 | | |
| 276 | hsa-miR-195-5p | MIMAT0000461 | | 20 | 25 | 21 | 21 | 20 | 24 | 22 | 11 | 39 | 29 | 23.2 | 22 | 24 | 0.90 | | |
| 277 | hsa-miR-526b-5p | MIMAT0002835 | | 27 | 19 | 27 | 21 | 24 | 24 | 18 | 15 | 41 | 16 | 23.2 | 24 | 23 | 1.02 | | |
| 278 | hsa-miR-147a | MIMAT0000251 | | 19 | 21 | 26 | 10 | 35 | 30 | 21 | 16 | 32 | 21 | 23.1 | 19 | 26 | 0.74 | | |
| 279 | hsa-miR-181a-3p | MIMAT0000270 | | 19 | 23 | 31 | 16 | 35 | 24 | 20 | 13 | 34 | 16 | 23.1 | 22 | 24 | 0.94 | | |
| 280 | hsa-miR-188-5p | MIMAT0000457 | | 18 | 22 | 32 | 16 | 27 | 20 | 25 | 9 | 35 | 27 | 23.1 | 22 | 24 | 0.92 | | |

| | | | | | | | | | | | | | | | | | | | | |
|---|---|---|---|---|---|---|---|---|---|---|---|---|---|---|---|---|---|---|---|---|
| 281 | hsa-miR-1973 | MIMAT0009448 | | 26 | 29 | 22 | 24 | 29 | 19 | 26 | 8 | 27 | 21 | 23.1 | 25 | 22 | 1.17 | | | |
| 282 | hsa-miR-25-5p | MIMAT0004498 | | 15 | 22 | 29 | 19 | 21 | 31 | 27 | 19 | 36 | 12 | 23.1 | 21 | 24 | 0.87 | | | |
| 283 | hsa-miR-3151-5p | MIMAT0015024 | | 20 | 27 | 34 | 9 | 33 | 22 | 21 | 16 | 28 | 21 | 23.1 | 23 | 24 | 0.96 | | | |
| 284 | hsa-miR-551b-3p | MIMAT0003233 | | 26 | 21 | 23 | 16 | 41 | 23 | 23 | 22 | 23 | 13 | 23.1 | 22 | 24 | 0.89 | | | |
| 285 | hsa-miR-592 | MIMAT0003260 | | 16 | 14 | 26 | 31 | 21 | 27 | 16 | 17 | 32 | 31 | 23.1 | 22 | 24 | 0.91 | | | |
| 286 | hsa-miR-939-5p | MIMAT0004982 | | 15 | 23 | 32 | 21 | 32 | 20 | 24 | 13 | 31 | 20 | 23.1 | 23 | 23 | 0.98 | | | |
| 287 | hsa-miR-1322 | MIMAT0005953 | | 18 | 26 | 24 | 17 | 23 | 23 | 18 | 14 | 45 | 22 | 23 | 21 | 24 | 0.88 | | | |
| 288 | hsa-miR-18a-5p | MIMAT0000072 | | 28 | 15 | 37 | 23 | 24 | 34 | 11 | 10 | 32 | 16 | 23 | 26 | 21 | 1.22 | | | |
| 289 | hsa-miR-491-3p | MIMAT0004765 | | 23 | 27 | 26 | 23 | 23 | 19 | 21 | 16 | 24 | 28 | 23 | 25 | 22 | 1.13 | | | |
| 290 | hsa-miR-642a-5p | MIMAT0003312 | | 10 | 24 | 24 | 24 | 28 | 22 | 30 | 17 | 35 | 16 | 23 | 21 | 25 | 0.83 | | | |
| 291 | hsa-miR-374b-5p | MIMAT0004955 | | 20 | 27 | 30 | 17 | 24 | 28 | 21 | 12 | 36 | 13 | 22.8 | 24 | 22 | 1.05 | | | |
| 292 | hsa-miR-548k | MIMAT0005882 | | 31 | 21 | 19 | 30 | 23 | 17 | 20 | 16 | 30 | 21 | 22.8 | 25 | 21 | 1.19 | | | |
| 293 | hsa-miR-192-5p | MIMAT0000222 | | 15 | 37 | 29 | 19 | 35 | 17 | 17 | 11 | 32 | 15 | 22.7 | 25 | 21 | 1.18 | | | |
| 294 | hsa-miR-520c-3p | MIMAT0002846 | | 25 | 24 | 26 | 18 | 30 | 20 | 26 | 13 | 27 | 18 | 22.7 | 23 | 22 | 1.04 | | | |
| 295 | hsa-miR-210-3p | MIMAT0000267 | | 26 | 20 | 44 | 14 | 19 | 25 | 27 | 13 | 30 | 8 | 22.6 | 26 | 20 | 1.28 | | | |
| 296 | hsa-miR-4536-3p | MIMAT0020959 | | 23 | 25 | 17 | 19 | 29 | 20 | 23 | 15 | 37 | 18 | 22.6 | 21 | 24 | 0.89 | | | |
| 297 | hsa-miR-497-5p | MIMAT0002820 | | 27 | 24 | 27 | 23 | 22 | 19 | 23 | 16 | 21 | 23 | 22.5 | 25 | 21 | 1.22 | | | |
| 298 | hsa-miR-519d-3p | MIMAT0002853 | | 15 | 24 | 33 | 19 | 30 | 22 | 24 | 13 | 31 | 14 | 22.5 | 23 | 22 | 1.02 | | | |
| 299 | hsa-miR-874-5p | MIMAT0026718 | | 20 | 17 | 27 | 23 | 31 | 23 | 20 | 15 | 32 | 17 | 22.5 | 22 | 23 | 0.95 | | | |
| 300 | hsa-miR-125a-3p | MIMAT0004602 | | 18 | 25 | 21 | 14 | 30 | 27 | 19 | 19 | 36 | 15 | 22.4 | 20 | 24 | 0.80 | | | |
| 301 | hsa-miR-486-3p | MIMAT0004762 | | 24 | 29 | 26 | 12 | 37 | 23 | 23 | 9 | 29 | 12 | 22.4 | 23 | 22 | 1.03 | | | |
| 302 | hsa-miR-640 | MIMAT0003310 | | 19 | 21 | 30 | 16 | 21 | 25 | 28 | 19 | 29 | 16 | 22.4 | 22 | 23 | 0.93 | | | |
| 303 | hsa-miR-181b-5p | MIMAT0000257 | | 24 | 26 | 31 | 18 | 29 | 21 | 21 | 10 | 25 | 18 | 22.3 | 25 | 21 | 1.20 | | hsa-miR-181d-5p | |
| 304 | hsa-miR-199a-5p | MIMAT0000231 | | 24 | 23 | 30 | 20 | 23 | 19 | 14 | 16 | 31 | 23 | 22.3 | 24 | 21 | 1.15 | | | |
| 305 | hsa-miR-30a-3p | MIMAT0000088 | | 21 | 28 | 30 | 15 | 26 | 19 | 28 | 13 | 28 | 15 | 22.3 | 24 | 22 | 1.09 | | | |
| 306 | hsa-miR-564 | MIMAT0003228 | | 22 | 16 | 19 | 24 | 30 | 17 | 24 | 13 | 33 | 25 | 22.3 | 20 | 24 | 0.86 | | | |
| 307 | hsa-miR-105-5p | MIMAT0000102 | | 15 | 22 | 15 | 17 | 35 | 27 | 16 | 18 | 34 | 22 | 22.1 | 17 | 25 | 0.68 | | | |
| 308 | hsa-miR-1307-5p | MIMAT0022727 | | 24 | 18 | 25 | 25 | 22 | 20 | 21 | 15 | 33 | 18 | 22.1 | 23 | 22 | 1.07 | | | |
| 309 | hsa-miR-203a-5p | MIMAT0031890 | | 24 | 25 | 10 | 25 | 26 | 19 | 22 | 15 | 29 | 26 | 22.1 | 21 | 23 | 0.92 | | | |
| 310 | hsa-miR-455-5p | MIMAT0003150 | | 18 | 23 | 35 | 16 | 25 | 25 | 21 | 18 | 21 | 19 | 22.1 | 23 | 22 | 1.07 | | | |
| 311 | hsa-miR-610 | MIMAT0003278 | | 17 | 18 | 30 | 30 | 29 | 18 | 23 | 14 | 28 | 14 | 22.1 | 24 | 21 | 1.13 | | | |
| 312 | hsa-miR-215-5p | MIMAT0000272 | | 13 | 25 | 37 | 17 | 26 | 22 | 21 | 11 | 25 | 23 | 22 | 23 | 21 | 1.08 | | | |
| 313 | hsa-miR-218-5p | MIMAT0000275 | | 21 | 22 | 27 | 13 | 23 | 22 | 29 | 16 | 28 | 19 | 22 | 21 | 23 | 0.91 | | | |
| 314 | hsa-miR-548m | MIMAT0005917 | | 24 | 17 | 32 | 25 | 21 | 19 | 19 | 12 | 28 | 23 | 22 | 25 | 20 | 1.20 | | | |
| 315 | hsa-miR-626 | MIMAT0003295 | | 18 | 23 | 21 | 26 | 18 | 18 | 21 | 15 | 32 | 28 | 22 | 22 | 22 | 1.00 | | | |
| 316 | hsa-miR-663a | MIMAT0003326 | | 22 | 22 | 34 | 15 | 31 | 19 | 15 | 18 | 29 | 15 | 22 | 23 | 21 | 1.10 | | | |
| 317 | hsa-miR-96-5p | MIMAT0000095 | | 22 | 20 | 28 | 24 | 22 | 20 | 25 | 14 | 27 | 15 | 21.7 | 24 | 21 | 1.15 | | | |
| 318 | hsa-miR-516a-3p | MIMAT0006778 | | 16 | 24 | 20 | 19 | 35 | 24 | 18 | 13 | 30 | 17 | 21.6 | 20 | 23 | 0.86 | | hsa-miR-516b-3p | |
| 319 | hsa-miR-548d-5p | MIMAT0004812 | | 16 | 28 | 27 | 14 | 28 | 20 | 27 | 8 | 33 | 15 | 21.6 | 21 | 22 | 0.97 | | | |
| 320 | hsa-miR-601 | MIMAT0003269 | | 19 | 31 | 28 | 17 | 31 | 21 | 19 | 6 | 27 | 17 | 21.6 | 24 | 20 | 1.18 | | | |
| 321 | hsa-miR-802 | MIMAT0004185 | | 15 | 18 | 20 | 16 | 27 | 26 | 25 | 20 | 27 | 22 | 21.6 | 17 | 25 | 0.70 | | | |
| 322 | hsa-miR-433-5p | MIMAT0026554 | | 24 | 29 | 18 | 25 | 23 | 20 | 18 | 20 | 21 | 17 | 21.5 | 24 | 20 | 1.21 | | | |
| 323 | hsa-let-7e-5p | MIMAT0000066 | | 40 | 23 | 35 | 10 | 18 | 31 | 21 | 2 | 24 | 9 | 21.3 | 27 | 18 | 1.54 | | | |
| 324 | hsa-miR-1180-3p | MIMAT0005825 | | 26 | 16 | 34 | 15 | 24 | 23 | 23 | 6 | 31 | 14 | 21.2 | 23 | 20 | 1.13 | | | |
| 325 | hsa-miR-133a-5p | MIMAT0026478 | | 18 | 26 | 26 | 24 | 25 | 15 | 22 | 9 | 28 | 19 | 21.2 | 24 | 20 | 1.19 | | | |
| 326 | hsa-miR-138-5p | MIMAT0000430 | | 16 | 20 | 27 | 11 | 33 | 18 | 14 | 17 | 41 | 15 | 21.2 | 19 | 23 | 0.80 | | | |
| 327 | hsa-miR-216a-5p | MIMAT0000273 | | 17 | 22 | 28 | 12 | 25 | 21 | 19 | 12 | 34 | 22 | 21.2 | 20 | 22 | 0.89 | | | |
| 328 | hsa-miR-515-5p | MIMAT0002826 | | 27 | 14 | 20 | 14 | 28 | 16 | 22 | 14 | 29 | 27 | 21.1 | 19 | 23 | 0.83 | | | |
| 329 | hsa-miR-5196-5p | MIMAT0021128 | | 14 | 30 | 24 | 23 | 23 | 28 | 12 | 14 | 26 | 17 | 21.1 | 23 | 20 | 1.14 | | | |
| 330 | hsa-miR-573 | MIMAT0003238 | | 19 | 25 | 40 | 15 | 29 | 17 | 12 | 14 | 23 | 17 | 21.1 | 25 | 19 | 1.33 | | | |
| 331 | hsa-miR-1285-3p | MIMAT0005876 | | 19 | 21 | 23 | 20 | 19 | 20 | 22 | 14 | 34 | 18 | 21 | 21 | 21 | 0.98 | | | |
| 332 | hsa-miR-301b-5p | MIMAT0032026 | | 22 | 18 | 21 | 15 | 26 | 19 | 17 | 19 | 34 | 19 | 21 | 19 | 22 | 0.85 | | | |
| 333 | hsa-miR-34c-5p | MIMAT0000686 | | 18 | 18 | 32 | 14 | 24 | 24 | 24 | 12 | 21 | 23 | 21 | 21 | 21 | 0.96 | | | |
| 334 | hsa-miR-548ad-3p | MIMAT0018946 | | 21 | 20 | 30 | 21 | 27 | 16 | 23 | 17 | 29 | 6 | 21 | 23 | 20 | 1.17 | | | |
| 335 | hsa-miR-548l | MIMAT0005889 | | 12 | 25 | 33 | 21 | 31 | 27 | 13 | 12 | 26 | 10 | 21 | 23 | 20 | 1.15 | | | |
| 336 | hsa-miR-2110 | MIMAT0010133 | | 15 | 18 | 30 | 17 | 20 | 21 | 22 | 17 | 34 | 15 | 20.9 | 20 | 22 | 0.93 | | | |
| 337 | hsa-miR-363-5p | MIMAT0003385 | | 15 | 28 | 29 | 17 | 26 | 22 | 22 | 8 | 26 | 16 | 20.9 | 22 | 20 | 1.11 | | | |
| 338 | hsa-miR-520d-5p | MIMAT0002855 | | 16 | 24 | 37 | 19 | 32 | 18 | 12 | 7 | 22 | 22 | 20.9 | 24 | 19 | 1.27 | | hsa-miR-527 | hsa-miR-518a-5p |
| 339 | hsa-miR-323b-3p | MIMAT0015050 | | 20 | 25 | 25 | 20 | 24 | 17 | 23 | 11 | 28 | 14 | 20.7 | 23 | 20 | 1.15 | | | |
| 340 | hsa-miR-660-3p | MIMAT0022711 | | 21 | 23 | 18 | 17 | 17 | 19 | 25 | 18 | 24 | 25 | 20.7 | 20 | 21 | 0.93 | | | |
| 341 | hsa-miR-1185-2-3p | MIMAT0022713 | | 13 | 23 | 25 | 23 | 37 | 20 | 20 | 12 | 22 | 11 | 20.6 | 21 | 20 | 1.03 | | | |
| 342 | hsa-miR-1228-3p | MIMAT0005583 | | 24 | 7 | 16 | 19 | 27 | 18 | 28 | 17 | 34 | 16 | 20.6 | 17 | 23 | 0.71 | | | |
| 343 | hsa-miR-320a | MIMAT0000510 | | 19 | 18 | 25 | 12 | 31 | 26 | 23 | 9 | 34 | 9 | 20.6 | 19 | 22 | 0.84 | | | |
| 344 | hsa-miR-3614-3p | MIMAT0017993 | | 15 | 24 | 25 | 18 | 27 | 20 | 23 | 9 | 31 | 14 | 20.6 | 21 | 21 | 0.99 | | | |
| 345 | hsa-miR-582-5p | MIMAT0003247 | | 21 | 14 | 28 | 20 | 22 | 17 | 16 | 15 | 30 | 23 | 20.6 | 21 | 21 | 1.01 | | | |
| 346 | hsa-miR-620 | MIMAT0003289 | | 11 | 28 | 24 | 15 | 23 | 15 | 31 | 9 | 34 | 16 | 20.6 | 20 | 21 | 0.91 | | | |
| 347 | hsa-miR-889-3p | MIMAT0004921 | | 8 | 16 | 20 | 26 | 25 | 14 | 23 | 20 | 35 | 19 | 20.6 | 18 | 23 | 0.77 | | | |
| 348 | hsa-miR-146a-5p | MIMAT0000449 | | 12 | 16 | 22 | 20 | 25 | 21 | 21 | 17 | 33 | 18 | 20.5 | 18 | 23 | 0.78 | | | |
| 349 | hsa-miR-517c-3p | MIMAT0002866 | | 12 | 18 | 22 | 22 | 28 | 18 | 21 | 14 | 33 | 17 | 20.5 | 19 | 22 | 0.85 | | hsa-miR-519a-3p | |
| 350 | hsa-miR-1255b-5p | MIMAT0005945 | | 17 | 23 | 26 | 21 | 25 | 14 | 21 | 17 | 27 | 13 | 20.4 | 22 | 20 | 1.12 | | | |
| 351 | hsa-miR-3164 | MIMAT0015038 | | 24 | 18 | 23 | 26 | 26 | 11 | 25 | 13 | 25 | 12 | 20.3 | 23 | 19 | 1.22 | | | |
| 352 | hsa-miR-671-5p | MIMAT0003880 | | 26 | 21 | 12 | 22 | 14 | 15 | 20 | 20 | 31 | 22 | 20.3 | 20 | 20 | 1.00 | | | |
| 353 | hsa-miR-382-5p | MIMAT0000737 | | 25 | 29 | 18 | 19 | 17 | 15 | 13 | 13 | 36 | 17 | 20.2 | 23 | 19 | 1.23 | | | |
| 354 | hsa-miR-504-5p | MIMAT0002875 | | 14 | 20 | 24 | 11 | 30 | 19 | 22 | 5 | 29 | 28 | 20.2 | 17 | 22 | 0.78 | | | |
| 355 | hsa-miR-769-5p | MIMAT0003886 | | 20 | 23 | 28 | 13 | 20 | 14 | 22 | 12 | 35 | 15 | 20.2 | 21 | 20 | 1.07 | | | |
| 356 | hsa-miR-198 | MIMAT0000228 | | 15 | 16 | 30 | 15 | 28 | 22 | 21 | 8 | 29 | 17 | 20.1 | 19 | 21 | 0.91 | | | |
| 357 | hsa-miR-1206 | MIMAT0005870 | | 20 | 23 | 30 | 11 | 29 | 17 | 22 | 14 | 19 | 15 | 20 | 21 | 19 | 1.09 | | | |
| 358 | hsa-miR-4451 | MIMAT0018973 | | 19 | 20 | 26 | 14 | 20 | 28 | 20 | 13 | 20 | 20 | 20 | 20 | 20 | 0.98 | | | |
| 359 | hsa-miR-4647 | MIMAT0019709 | | 17 | 25 | 18 | 26 | 26 | 21 | 18 | 16 | 19 | 14 | 20 | 22 | 19 | 1.13 | | | |
| 360 | hsa-miR-545-3p | MIMAT0003165 | | 21 | 29 | 23 | 15 | 22 | 13 | 15 | 19 | 29 | 14 | 20 | 22 | 19 | 1.18 | | | |
| 361 | hsa-miR-596 | MIMAT0003264 | | 16 | 20 | 21 | 19 | 20 | 30 | 23 | 10 | 28 | 13 | 20 | 19 | 21 | 0.92 | | | |
| 362 | hsa-miR-625-5p | MIMAT0003294 | | 19 | 17 | 31 | 16 | 28 | 22 | 11 | 10 | 29 | 17 | 20 | 21 | 20 | 1.06 | | | |
| 363 | hsa-miR-298 | MIMAT0004901 | | 20 | 22 | 25 | 14 | 38 | 19 | 11 | 11 | 24 | 15 | 19.9 | 20 | 20 | 1.03 | | | |
| 364 | hsa-miR-410-3p | MIMAT0002171 | | 19 | 32 | 26 | 16 | 12 | 25 | 14 | 15 | 28 | 12 | 19.9 | 23 | 18 | 1.32 | | | |
| 365 | hsa-miR-510-3p | MIMAT0026613 | | 12 | 21 | 19 | 14 | 27 | 17 | 20 | 20 | 38 | 11 | 19.9 | 17 | 22 | 0.74 | | | |
| 366 | hsa-miR-555 | MIMAT0003219 | | 19 | 19 | 29 | 16 | 23 | 18 | 23 | 16 | 23 | 13 | 19.9 | 21 | 19 | 1.07 | | | |
| 367 | hsa-miR-892a | MIMAT0004907 | | 18 | 20 | 22 | 17 | 23 | 22 | 17 | 12 | 30 | 18 | 19.9 | 19 | 20 | 0.95 | | | |
| 368 | hsa-miR-190b | MIMAT0004929 | | 14 | 16 | 21 | 22 | 26 | 26 | 20 | 6 | 36 | 11 | 19.8 | 18 | 21 | 0.88 | | | |
| 369 | hsa-miR-512-5p | MIMAT0002822 | | 20 | 20 | 21 | 13 | 19 | 29 | 20 | 15 | 26 | 15 | 19.8 | 19 | 21 | 0.90 | | | |
| 370 | hsa-miR-628-3p | MIMAT0003297 | | 13 | 20 | 22 | 16 | 25 | 18 | 20 | 15 | 34 | 15 | 19.8 | 18 | 21 | 0.84 | | | |
| 371 | hsa-miR-639 | MIMAT0003309 | | 15 | 20 | 33 | 11 | 29 | 10 | 21 | 10 | 27 | 22 | 19.8 | 20 | 20 | 1.00 | | | |
| 372 | hsa-miR-6503-5p | MIMAT0025462 | | 19 | 24 | 19 | 15 | 16 | 26 | 14 | 18 | 27 | 20 | 19.8 | 19 | 20 | 0.95 | | | |
| 373 | hsa-miR-144-3p | MIMAT0000436 | | 21 | 24 | 22 | 14 | 25 | 20 | 16 | 9 | 25 | 21 | 19.7 | 20 | 19 | 1.05 | | | |
| 374 | hsa-miR-147b | MIMAT0004928 | | 20 | 14 | 22 | 17 | 24 | 28 | 20 | 17 | 24 | 11 | 19.7 | 18 | 21 | 0.88 | | | |
| 375 | hsa-miR-184 | MIMAT0000454 | | 11 | 20 | 27 | 20 | 19 | 18 | 10 | 19 | 34 | 19 | 19.7 | 20 | 20 | 0.98 | | | |

| | | | | | | | | | | | | | | | | | |
|---|---|---|---|---|---|---|---|---|---|---|---|---|---|---|---|---|---|
| 376 | hsa-miR-211-5p | MIMAT0000268 | | 15 | 16 | 18 | 23 | 24 | 25 | 21 | 14 | 24 | 17 | 19.7 | 18 | 21 | 0.86 |
| 377 | hsa-miR-30e-3p | MIMAT0000693 | | 15 | 15 | 32 | 22 | 24 | 21 | 19 | 11 | 22 | 16 | 19.7 | 21 | 19 | 1.12 |
| 378 | hsa-miR-328-5p | MIMAT0026486 | | 19 | 19 | 28 | 18 | 28 | 11 | 25 | 16 | 19 | 14 | 19.7 | 21 | 19 | 1.12 |
| 379 | hsa-miR-34c-3p | MIMAT0004677 | | 21 | 20 | 28 | 21 | 23 | 17 | 14 | 14 | 19 | 20 | 19.7 | 23 | 18 | 1.26 |
| 380 | hsa-miR-4707-3p | MIMAT0019808 | | 19 | 17 | 29 | 14 | 22 | 19 | 18 | 17 | 28 | 14 | 19.7 | 20 | 20 | 1.00 |
| 381 | hsa-miR-506-3p | MIMAT0002878 | | 20 | 26 | 19 | 17 | 15 | 28 | 18 | 10 | 30 | 14 | 19.7 | 21 | 19 | 1.07 |
| 382 | hsa-miR-629-5p | MIMAT0004810 | | 16 | 23 | 18 | 15 | 31 | 16 | 16 | 15 | 28 | 19 | 19.7 | 18 | 21 | 0.86 |
| 383 | hsa-miR-941 | MIMAT0004984 | | 16 | 18 | 21 | 18 | 22 | 19 | 16 | 17 | 37 | 13 | 19.7 | 18 | 21 | 0.88 |
| 384 | hsa-miR-1286 | MIMAT0005877 | | 13 | 19 | 20 | 16 | 28 | 24 | 20 | 12 | 23 | 21 | 19.6 | 17 | 21 | 0.80 |
| 385 | hsa-miR-139-3p | MIMAT0004552 | | 15 | 17 | 27 | 16 | 17 | 25 | 16 | 8 | 28 | 27 | 19.6 | 19 | 20 | 0.93 |
| 386 | hsa-miR-382-3p | MIMAT0022697 | | 12 | 23 | 18 | 11 | 21 | 21 | 25 | 11 | 39 | 15 | 19.6 | 16 | 22 | 0.73 |
| 387 | hsa-miR-484 | MIMAT0002174 | | 20 | 14 | 22 | 13 | 27 | 18 | 15 | 16 | 29 | 22 | 19.6 | 17 | 21 | 0.81 |
| 388 | hsa-miR-593-3p | MIMAT0004802 | | 14 | 22 | 27 | 16 | 22 | 19 | 24 | 15 | 23 | 14 | 19.6 | 20 | 20 | 1.01 |
| 389 | hsa-miR-508-5p | MIMAT0004778 | | 15 | 17 | 20 | 14 | 27 | 19 | 17 | 14 | 30 | 22 | 19.5 | 17 | 22 | 0.77 |
| 390 | hsa-miR-136-5p | MIMAT0000448 | | 19 | 26 | 22 | 16 | 19 | 23 | 12 | 14 | 26 | 17 | 19.4 | 21 | 19 | 1.12 |
| 391 | hsa-miR-367-3p | MIMAT0000719 | | 14 | 16 | 11 | 17 | 43 | 16 | 15 | 14 | 34 | 14 | 19.4 | 15 | 23 | 0.64 |
| 392 | hsa-miR-556-5p | MIMAT0003220 | | 23 | 15 | 23 | 15 | 28 | 20 | 16 | 9 | 34 | 11 | 19.4 | 19 | 20 | 0.97 |
| 393 | hsa-miR-638 | MIMAT0003308 | | 25 | 15 | 22 | 14 | 24 | 26 | 19 | 8 | 30 | 11 | 19.4 | 19 | 20 | 0.97 |
| 394 | hsa-miR-1245a | MIMAT0005897 | | 20 | 21 | 23 | 23 | 20 | 16 | 18 | 11 | 26 | 15 | 19.3 | 22 | 18 | 1.23 |
| 395 | hsa-miR-1249-3p | MIMAT0005901 | | 12 | 23 | 31 | 14 | 24 | 15 | 19 | 14 | 21 | 20 | 19.3 | 20 | 19 | 1.06 |
| 396 | hsa-miR-331-3p | MIMAT0000760 | | 21 | 23 | 33 | 17 | 23 | 16 | 11 | 11 | 25 | 13 | 19.3 | 24 | 17 | 1.42 |
| 397 | hsa-miR-1269a | MIMAT0005923 | | 20 | 24 | 24 | 16 | 23 | 11 | 19 | 8 | 38 | 8 | 19.1 | 21 | 18 | 1.18 |
| 398 | hsa-miR-296-5p | MIMAT0000690 | | 19 | 23 | 24 | 19 | 19 | 19 | 23 | 7 | 16 | 22 | 19.1 | 21 | 18 | 1.20 |
| 399 | hsa-miR-3074-3p | MIMAT0015027 | | 9 | 20 | 22 | 18 | 33 | 20 | 22 | 14 | 18 | 15 | 19.1 | 17 | 20 | 0.85 |
| 400 | hsa-miR-369-3p | MIMAT0000721 | | 20 | 23 | 21 | 15 | 22 | 19 | 21 | 14 | 24 | 12 | 19.1 | 20 | 19 | 1.06 |
| 401 | hsa-miR-432-5p | MIMAT0002814 | | 16 | 21 | 26 | 13 | 23 | 23 | 24 | 8 | 23 | 14 | 19.1 | 19 | 19 | 0.99 |
| 402 | hsa-miR-499b-3p | MIMAT0019898 | | 21 | 21 | 22 | 13 | 21 | 16 | 17 | 14 | 32 | 14 | 19.1 | 19 | 19 | 1.01 |
| 403 | hsa-miR-513b-5p | MIMAT0005788 | | 14 | 28 | 22 | 13 | 21 | 19 | 19 | 15 | 25 | 15 | 19.1 | 19 | 19 | 1.01 |
| 404 | hsa-miR-519c-3p | MIMAT0002832 | | 20 | 13 | 30 | 18 | 18 | 22 | 16 | 16 | 23 | 15 | 19.1 | 20 | 18 | 1.10 |
| 405 | hsa-miR-520b | MIMAT0002843 | | 19 | 30 | 12 | 12 | 27 | 25 | 10 | 6 | 30 | 20 | 19.1 | 18 | 20 | 0.93 |
| 406 | hsa-miR-664a-3p | MIMAT0005949 | | 10 | 12 | 24 | 15 | 20 | 19 | 23 | 9 | 34 | 25 | 19.1 | 15 | 22 | 0.70 |
| 407 | hsa-miR-1537-3p | MIMAT0007399 | | 13 | 16 | 27 | 15 | 22 | 15 | 21 | 12 | 30 | 19 | 19 | 18 | 20 | 0.89 |
| 408 | hsa-miR-29c-3p | MIMAT0000681 | | 26 | 14 | 44 | 12 | 19 | 29 | 20 | 6 | 15 | 5 | 19 | 24 | 16 | 1.53 |
| 409 | hsa-miR-323b-5p | MIMAT0001630 | | 15 | 25 | 29 | 10 | 24 | 22 | 21 | 9 | 21 | 14 | 19 | 20 | 19 | 1.07 |
| 410 | hsa-miR-409-3p | MIMAT0001639 | | 22 | 15 | 22 | 13 | 22 | 22 | 16 | 16 | 21 | 21 | 19 | 18 | 20 | 0.92 |
| 411 | hsa-miR-429 | MIMAT0001536 | | 11 | 15 | 26 | 19 | 21 | 21 | 17 | 19 | 21 | 20 | 19 | 18 | 20 | 0.89 |
| 412 | hsa-miR-1178-3p | MIMAT0005823 | | 26 | 19 | 32 | 9 | 20 | 19 | 15 | 11 | 25 | 13 | 18.9 | 22 | 17 | 1.25 |
| 413 | hsa-miR-1203 | MIMAT0005866 | | 21 | 12 | 21 | 16 | 19 | 14 | 21 | 14 | 33 | 18 | 18.9 | 18 | 20 | 0.88 |
| 414 | hsa-miR-1298-5p | MIMAT0005800 | | 10 | 19 | 23 | 18 | 32 | 11 | 21 | 17 | 22 | 16 | 18.9 | 18 | 20 | 0.88 |
| 415 | hsa-miR-329-5p | MIMAT0026555 | | 17 | 17 | 26 | 18 | 29 | 16 | 15 | 7 | 27 | 17 | 18.9 | 20 | 19 | 1.05 |
| 416 | hsa-miR-448 | MIMAT0001532 | | 11 | 23 | 20 | 22 | 24 | 15 | 17 | 12 | 30 | 15 | 18.9 | 19 | 19 | 1.01 |
| 417 | hsa-miR-579-5p | MIMAT0026616 | | 19 | 20 | 25 | 9 | 26 | 17 | 15 | 12 | 27 | 19 | 18.9 | 18 | 19 | 0.94 |
| 418 | hsa-miR-185-5p | MIMAT0000455 | | 11 | 18 | 16 | 15 | 21 | 21 | 14 | 15 | 36 | 21 | 18.8 | 15 | 21 | 0.70 |
| 419 | hsa-miR-302c-3p | MIMAT0000717 | | 10 | 17 | 32 | 18 | 22 | 17 | 13 | 9 | 32 | 18 | 18.8 | 19 | 19 | 1.04 |
| 420 | hsa-miR-4485-3p | MIMAT0019019 | | 15 | 27 | 19 | 23 | 25 | 10 | 21 | 10 | 22 | 15 | 18.7 | 21 | 17 | 1.22 |
| 421 | hsa-miR-933 | MIMAT0004976 | | 18 | 21 | 23 | 29 | 21 | 18 | 16 | 10 | 19 | 12 | 18.7 | 23 | 16 | 1.42 |
| 422 | hsa-miR-1304-5p | MIMAT0005892 | | 22 | 18 | 23 | 16 | 18 | 21 | 23 | 16 | 21 | 8 | 18.6 | 20 | 18 | 1.11 |
| 423 | hsa-miR-135b-5p | MIMAT0000758 | | 19 | 14 | 42 | 12 | 20 | 10 | 16 | 13 | 26 | 14 | 18.6 | 22 | 17 | 1.32 |
| 424 | hsa-miR-328-3p | MIMAT0000752 | | 13 | 26 | 23 | 9 | 20 | 15 | 15 | 11 | 25 | 29 | 18.6 | 18 | 19 | 0.93 |
| 425 | hsa-miR-571 | MIMAT0003236 | | 19 | 21 | 23 | 20 | 19 | 16 | 15 | 14 | 20 | 19 | 18.6 | 21 | 17 | 1.21 |
| 426 | hsa-miR-1234-3p | MIMAT0005589 | | 18 | 21 | 33 | 10 | 19 | 13 | 22 | 10 | 19 | 20 | 18.5 | 21 | 17 | 1.19 |
| 427 | hsa-miR-28-3p | MIMAT0004502 | | 12 | 18 | 21 | 17 | 11 | 23 | 12 | 17 | 36 | 18 | 18.5 | 17 | 20 | 0.87 |
| 428 | hsa-miR-302e | MIMAT0005931 | | 24 | 14 | 20 | 14 | 22 | 14 | 12 | 10 | 33 | 22 | 18.5 | 18 | 19 | 0.96 |
| 429 | hsa-miR-4792 | MIMAT0019964 | | 12 | 17 | 22 | 27 | 21 | 16 | 18 | 12 | 22 | 18 | 18.5 | 20 | 18 | 1.09 |
| 430 | hsa-miR-513a-3p | MIMAT0004777 | | 19 | 16 | 19 | 12 | 14 | 23 | 21 | 17 | 24 | 20 | 18.5 | 17 | 20 | 0.83 |
| 431 | hsa-miR-642a-3p | MIMAT0020924 | | 15 | 28 | 34 | 14 | 18 | 15 | 16 | 9 | 25 | 11 | 18.5 | 23 | 16 | 1.45 |
| 432 | hsa-miR-126-3p | MIMAT0000445 | | 25 | 11 | 33 | 13 | 15 | 31 | 24 | 8 | 16 | 8 | 18.4 | 21 | 17 | 1.21 |
| 433 | hsa-miR-3615 | MIMAT0017994 | | 19 | 20 | 21 | 23 | 17 | 18 | 17 | 8 | 22 | 19 | 18.4 | 21 | 17 | 1.23 |
| 434 | hsa-miR-450b-3p | MIMAT0004910 | | 17 | 14 | 21 | 17 | 14 | 29 | 20 | 9 | 22 | 21 | 18.4 | 17 | 19 | 0.90 |
| 435 | hsa-miR-509-5p | MIMAT0004779 | | 27 | 8 | 17 | 17 | 31 | 15 | 15 | 12 | 20 | 22 | 18.4 | 17 | 19 | 0.90 |
| 436 | hsa-miR-1307-3p | MIMAT0005951 | | 18 | 24 | 17 | 16 | 27 | 13 | 18 | 12 | 21 | 17 | 18.3 | 19 | 18 | 1.04 |
| 437 | hsa-miR-137 | MIMAT0000429 | | 16 | 17 | 30 | 11 | 31 | 15 | 15 | 11 | 26 | 11 | 18.3 | 19 | 18 | 1.02 |
| 438 | hsa-miR-181b-2-3p | MIMAT0031893 | | 14 | 15 | 21 | 16 | 29 | 25 | 18 | 6 | 30 | 9 | 18.3 | 17 | 20 | 0.85 |
| 439 | hsa-miR-190a-3p | MIMAT0026482 | | 14 | 27 | 19 | 17 | 20 | 17 | 17 | 12 | 25 | 15 | 18.3 | 19 | 18 | 1.09 |
| 440 | hsa-miR-2113 | MIMAT0009206 | | 12 | 23 | 21 | 20 | 17 | 18 | 27 | 7 | 21 | 17 | 18.3 | 19 | 18 | 1.07 |
| 441 | hsa-miR-561-3p | MIMAT0003225 | | 19 | 15 | 27 | 12 | 29 | 15 | 19 | 12 | 24 | 11 | 18.3 | 18 | 18 | 1.00 |
| 442 | hsa-miR-217 | MIMAT0000274 | | 13 | 18 | 20 | 21 | 24 | 10 | 19 | 12 | 31 | 14 | 18.2 | 18 | 18 | 0.98 |
| 443 | hsa-miR-485-3p | MIMAT0002176 | | 23 | 18 | 18 | 14 | 20 | 20 | 18 | 12 | 26 | 12 | 18.1 | 18 | 18 | 1.01 |
| 444 | hsa-miR-613 | MIMAT0003281 | | 17 | 15 | 20 | 21 | 29 | 16 | 15 | 14 | 25 | 9 | 18.1 | 18 | 18 | 1.01 |
| 445 | hsa-miR-1224-5p | MIMAT0005458 | | 13 | 13 | 29 | 15 | 21 | 22 | 15 | 16 | 17 | 19 | 18 | 18 | 18 | 0.95 |
| 446 | hsa-miR-1266-5p | MIMAT0005920 | | 15 | 21 | 13 | 14 | 27 | 14 | 18 | 20 | 23 | 15 | 18 | 16 | 20 | 0.81 |
| 447 | hsa-miR-523-3p | MIMAT0002840 | | 19 | 35 | 14 | 10 | 15 | 26 | 18 | 9 | 23 | 11 | 18 | 20 | 17 | 1.15 |
| 448 | hsa-miR-1185-5p | MIMAT0005798 | | 10 | 19 | 24 | 20 | 15 | 15 | 20 | 12 | 25 | 19 | 17.9 | 18 | 18 | 1.03 |
| 449 | hsa-miR-214-3p | MIMAT0000271 | | 20 | 21 | 22 | 17 | 27 | 14 | 11 | 10 | 24 | 13 | 17.9 | 20 | 17 | 1.21 |
| 450 | hsa-miR-337-5p | MIMAT0004695 | | 16 | 16 | 19 | 10 | 22 | 21 | 20 | 15 | 32 | 8 | 17.9 | 15 | 20 | 0.78 |
| 451 | hsa-miR-376a-2-5p | MIMAT0022928 | | 22 | 20 | 28 | 12 | 14 | 14 | 22 | 7 | 22 | 18 | 17.9 | 21 | 16 | 1.27 |
| 452 | hsa-miR-4755-5p | MIMAT0019895 | | 14 | 16 | 19 | 12 | 26 | 22 | 16 | 16 | 22 | 16 | 17.9 | 15 | 20 | 0.78 |
| 453 | hsa-miR-4787-3p | MIMAT0019957 | | 17 | 21 | 13 | 16 | 25 | 21 | 20 | 12 | 23 | 11 | 17.9 | 17 | 19 | 0.90 |
| 454 | hsa-miR-493-3p | MIMAT0003161 | | 19 | 20 | 24 | 21 | 20 | 18 | 17 | 7 | 24 | 9 | 17.9 | 21 | 16 | 1.33 |
| 455 | hsa-miR-575 | MIMAT0003240 | | 18 | 26 | 28 | 13 | 16 | 17 | 15 | 9 | 26 | 11 | 17.9 | 21 | 16 | 1.36 |
| 456 | hsa-miR-591 | MIMAT0003259 | | 16 | 22 | 20 | 19 | 15 | 18 | 15 | 11 | 19 | 24 | 17.9 | 19 | 17 | 1.13 |
| 457 | hsa-miR-1287-5p | MIMAT0005878 | | 16 | 20 | 16 | 18 | 12 | 22 | 16 | 11 | 26 | 21 | 17.8 | 18 | 18 | 0.97 |
| 458 | hsa-miR-491-5p | MIMAT0002807 | | 21 | 18 | 23 | 15 | 26 | 13 | 19 | 11 | 19 | 13 | 17.8 | 19 | 17 | 1.14 |
| 459 | hsa-miR-106b-5p | MIMAT0000680 | | 23 | 17 | 27 | 14 | 17 | 17 | 15 | 12 | 22 | 13 | 17.7 | 20 | 16 | 1.27 |
| 460 | hsa-miR-208a-3p | MIMAT0000241 | | 12 | 15 | 16 | 23 | 14 | 18 | 26 | 12 | 25 | 16 | 17.7 | 17 | 19 | 0.89 |
| 461 | hsa-miR-3928-3p | MIMAT0018205 | | 16 | 14 | 26 | 10 | 19 | 15 | 20 | 13 | 25 | 19 | 17.7 | 17 | 19 | 0.89 |
| 462 | hsa-miR-374a-3p | MIMAT0004688 | | 18 | 10 | 30 | 13 | 24 | 22 | 19 | 7 | 22 | 11 | 17.6 | 18 | 18 | 1.01 |
| 463 | hsa-miR-576-5p | MIMAT0003241 | | 27 | 24 | 18 | 10 | 18 | 17 | 22 | 11 | 16 | 13 | 17.6 | 20 | 16 | 1.22 |
| 464 | hsa-miR-590-3p | MIMAT0004801 | | 17 | 15 | 30 | 12 | 19 | 22 | 15 | 6 | 27 | 13 | 17.6 | 19 | 17 | 1.09 |
| 465 | hsa-miR-767-5p | MIMAT0003882 | | 18 | 19 | 20 | 9 | 19 | 11 | 26 | 10 | 25 | 19 | 17.6 | 17 | 18 | 0.90 |
| 466 | hsa-miR-124-3p | MIMAT0000422 | | 13 | 16 | 15 | 17 | 26 | 12 | 30 | 11 | 13 | 22 | 17.5 | 15 | 19 | 0.80 |
| 467 | hsa-miR-140-5p | MIMAT0000431 | | 16 | 23 | 22 | 12 | 17 | 22 | 15 | 13 | 18 | 17 | 17.5 | 18 | 17 | 1.07 |
| 468 | hsa-miR-584-5p | MIMAT0003249 | | 21 | 14 | 21 | 9 | 24 | 17 | 16 | 7 | 28 | 18 | 17.5 | 16 | 18 | 0.89 |
| 469 | hsa-miR-532-5p | MIMAT0002888 | | 17 | 32 | 28 | 12 | 9 | 25 | 15 | 8 | 16 | 12 | 17.4 | 22 | 14 | 1.57 |
| 470 | hsa-miR-134-3p | MIMAT0026481 | | 14 | 15 | 24 | 18 | 18 | 18 | 17 | 20 | 17 | 12 | 17.3 | 18 | 17 | 1.04 |

| | | | | | | | | | | | | | | | | | | | | |
|---|---|---|---|---|---|---|---|---|---|---|---|---|---|---|---|---|---|---|---|---|
| 471 | hsa-miR-15a-5p | MIMAT0000068 | | 22 | 13 | 34 | 7 | 15 | 22 | 22 | 10 | 23 | 5 | 17.3 | 19 | 16 | 1.18 | | | |
| 472 | hsa-miR-362-5p | MIMAT0000705 | | 13 | 15 | 18 | 26 | 20 | 21 | 15 | 10 | 16 | 19 | 17.3 | 18 | 17 | 1.07 | | | |
| 473 | hsa-miR-4488 | MIMAT0019022 | | 18 | 16 | 22 | 16 | 23 | 22 | 14 | 16 | 18 | 8 | 17.3 | 18 | 17 | 1.07 | | | |
| 474 | hsa-miR-1248 | MIMAT0005900 | | 16 | 19 | 22 | 20 | 23 | 15 | 12 | 9 | 20 | 16 | 17.2 | 19 | 16 | 1.22 | | | |
| 475 | hsa-miR-1289 | MIMAT0005879 | | 17 | 15 | 19 | 12 | 26 | 21 | 16 | 8 | 18 | 20 | 17.2 | 16 | 18 | 0.87 | | | |
| 476 | hsa-miR-302f | MIMAT0005932 | | 13 | 19 | 22 | 20 | 15 | 17 | 13 | 10 | 27 | 16 | 17.2 | 19 | 16 | 1.13 | | | |
| 477 | hsa-miR-33b-5p | MIMAT0003301 | | 9 | 20 | 17 | 13 | 16 | 19 | 23 | 17 | 24 | 14 | 17.2 | 15 | 19 | 0.78 | | | |
| 478 | hsa-miR-3934-5p | MIMAT0018349 | | 12 | 15 | 21 | 9 | 16 | 23 | 22 | 16 | 28 | 10 | 17.2 | 14 | 19 | 0.74 | | | |
| 479 | hsa-miR-4787-5p | MIMAT0019956 | | 20 | 22 | 21 | 11 | 16 | 18 | 16 | 14 | 20 | 14 | 17.2 | 19 | 16 | 1.13 | | | |
| 480 | hsa-miR-1299 | MIMAT0005887 | | 16 | 18 | 22 | 15 | 25 | 12 | 15 | 14 | 21 | 13 | 17.1 | 18 | 17 | 1.07 | | | |
| 481 | hsa-miR-300 | MIMAT0004903 | | 15 | 26 | 23 | 18 | 27 | 7 | 14 | 7 | 24 | 10 | 17.1 | 21 | 15 | 1.38 | | | |
| 482 | hsa-miR-5010-3p | MIMAT0021044 | | 7 | 22 | 22 | 19 | 27 | 13 | 17 | 13 | 22 | 9 | 17.1 | 18 | 17 | 1.04 | | | |
| 483 | hsa-miR-145-5p | MIMAT0000437 | | 22 | 13 | 20 | 15 | 21 | 27 | 8 | 7 | 22 | 15 | 17 | 18 | 17 | 1.05 | | | |
| 484 | hsa-miR-568 | MIMAT0003232 | | 15 | 15 | 25 | 18 | 24 | 17 | 16 | 7 | 21 | 12 | 17 | 18 | 16 | 1.13 | | | |
| 485 | hsa-miR-98-3p | MIMAT0022842 | | 23 | 13 | 19 | 15 | 23 | 19 | 18 | 14 | 15 | 11 | 17 | 18 | 17 | 1.05 | | | |
| 486 | hsa-miR-1200 | MIMAT0005863 | | 12 | 18 | 21 | 18 | 21 | 7 | 23 | 9 | 21 | 19 | 16.9 | 17 | 17 | 1.04 | | | |
| 487 | hsa-miR-485-5p | MIMAT0002175 | | 16 | 15 | 24 | 9 | 27 | 13 | 23 | 7 | 25 | 10 | 16.9 | 16 | 18 | 0.91 | | | |
| 488 | hsa-miR-520f-3p | MIMAT0002830 | | 15 | 22 | 14 | 16 | 30 | 16 | 19 | 10 | 19 | 8 | 16.9 | 17 | 17 | 0.99 | | | |
| 489 | hsa-miR-650 | MIMAT0003320 | | 17 | 16 | 15 | 18 | 12 | 16 | 27 | 9 | 25 | 14 | 16.9 | 17 | 17 | 0.96 | | | |
| 490 | hsa-miR-937-3p | MIMAT0004980 | | 12 | 15 | 25 | 15 | 27 | 15 | 16 | 11 | 22 | 11 | 16.9 | 17 | 17 | 0.99 | | | |
| 491 | hsa-miR-1281 | MIMAT0005939 | | 18 | 14 | 26 | 10 | 18 | 14 | 15 | 10 | 27 | 16 | 16.8 | 17 | 17 | 1.02 | | | |
| 492 | hsa-miR-628-5p | MIMAT0004809 | | 20 | 14 | 18 | 17 | 29 | 14 | 13 | 10 | 25 | 8 | 16.8 | 17 | 17 | 1.05 | | | |
| 493 | hsa-miR-4521 | MIMAT0019058 | | 10 | 11 | 22 | 16 | 26 | 20 | 16 | 9 | 17 | 20 | 16.7 | 15 | 18 | 0.82 | | | |
| 494 | hsa-miR-4435 | MIMAT0018951 | | 18 | 17 | 16 | 14 | 18 | 20 | 17 | 13 | 20 | 13 | 16.6 | 16 | 17 | 0.97 | | | |
| 495 | hsa-miR-576-3p | MIMAT0004796 | | 20 | 19 | 25 | 7 | 16 | 17 | 21 | 8 | 25 | 8 | 16.6 | 18 | 16 | 1.12 | | | |
| 496 | hsa-miR-760 | MIMAT0004957 | | 13 | 12 | 23 | 8 | 26 | 13 | 24 | 12 | 23 | 12 | 16.6 | 14 | 18 | 0.76 | | | |
| 497 | hsa-miR-194-5p | MIMAT0000460 | | 20 | 22 | 14 | 16 | 16 | 23 | 16 | 9 | 11 | 18 | 16.5 | 18 | 16 | 1.16 | | | |
| 498 | hsa-miR-496 | MIMAT0002818 | | 12 | 16 | 15 | 11 | 20 | 15 | 22 | 7 | 37 | 10 | 16.5 | 14 | 19 | 0.73 | | | |
| 499 | hsa-miR-509-3-5p | MIMAT0004975 | | 7 | 26 | 24 | 5 | 24 | 18 | 12 | 7 | 24 | 18 | 16.5 | 16 | 17 | 0.90 | | | |
| 500 | hsa-miR-509-3p | MIMAT0002881 | | 18 | 12 | 13 | 16 | 19 | 17 | 18 | 17 | 19 | 16 | 16.5 | 15 | 18 | 0.83 | | | |
| 501 | hsa-miR-942-5p | MIMAT0004985 | | 14 | 17 | 29 | 11 | 17 | 18 | 13 | 8 | 25 | 13 | 16.5 | 18 | 16 | 1.13 | | | |
| 502 | hsa-miR-513c-5p | MIMAT0005789 | | 15 | 17 | 20 | 18 | 18 | 19 | 17 | 12 | 16 | 12 | 16.4 | 18 | 16 | 1.12 | | | |
| 503 | hsa-miR-589-5p | MIMAT0004799 | | 9 | 19 | 14 | 12 | 12 | 20 | 20 | 10 | 27 | 20 | 16.3 | 14 | 18 | 0.74 | | | |
| 504 | hsa-miR-641 | MIMAT0003311 | | 16 | 18 | 20 | 17 | 22 | 18 | 13 | 9 | 20 | 10 | 16.3 | 18 | 15 | 1.16 | | | |
| 505 | hsa-miR-338-5p | MIMAT0004701 | | 10 | 11 | 18 | 13 | 20 | 14 | 22 | 18 | 24 | 12 | 16.2 | 13 | 18 | 0.71 | | | |
| 506 | hsa-miR-572 | MIMAT0003237 | | 17 | 13 | 20 | 10 | 26 | 11 | 17 | 14 | 21 | 13 | 16.2 | 15 | 17 | 0.88 | | | |
| 507 | hsa-miR-140-3p | MIMAT0004597 | | 11 | 19 | 31 | 9 | 17 | 17 | 23 | 13 | 11 | 10 | 16.1 | 18 | 15 | 1.15 | | | |
| 508 | hsa-miR-3130-3p | MIMAT0014994 | | 12 | 9 | 27 | 10 | 24 | 19 | 20 | 13 | 20 | 7 | 16.1 | 15 | 17 | 0.84 | | | |
| 509 | hsa-miR-3168 | MIMAT0015043 | | 15 | 21 | 23 | 11 | 19 | 13 | 19 | 7 | 17 | 16 | 16.1 | 18 | 15 | 1.15 | | | |
| 510 | hsa-miR-3202 | MIMAT0015089 | | 17 | 26 | 15 | 6 | 14 | 19 | 20 | 8 | 18 | 18 | 16.1 | 16 | 16 | 0.99 | | | |
| 511 | hsa-miR-1908-5p | MIMAT0007881 | | 16 | 19 | 11 | 10 | 18 | 17 | 22 | 6 | 22 | 19 | 16 | 14 | 17 | 0.81 | | | |
| 512 | hsa-miR-483-5p | MIMAT0004761 | | 17 | 22 | 14 | 18 | 18 | 13 | 15 | 9 | 16 | 18 | 16 | 18 | 15 | 1.20 | | | |
| 513 | hsa-miR-495-5p | MIMAT0022924 | | 22 | 15 | 18 | 9 | 20 | 13 | 21 | 12 | 21 | 9 | 16 | 16 | 16 | 1.00 | | | |
| 514 | hsa-miR-539-5p | MIMAT0003163 | | 13 | 18 | 20 | 9 | 21 | 10 | 18 | 11 | 23 | 16 | 15.9 | 15 | 17 | 0.91 | | | |
| 515 | hsa-miR-875-3p | MIMAT0004923 | | 17 | 13 | 24 | 13 | 15 | 11 | 19 | 7 | 25 | 15 | 15.9 | 17 | 15 | 1.09 | | | |
| 516 | hsa-miR-876-3p | MIMAT0004925 | | 11 | 12 | 10 | 18 | 24 | 18 | 20 | 13 | 17 | 16 | 15.9 | 13 | 18 | 0.71 | | | |
| 517 | hsa-miR-92a-1-5p | MIMAT0004507 | | 17 | 18 | 16 | 17 | 23 | 13 | 22 | 10 | 20 | 3 | 15.9 | 17 | 15 | 1.12 | | | |
| 518 | hsa-miR-1185-1-3p | MIMAT0022838 | | 14 | 15 | 16 | 13 | 19 | 15 | 19 | 6 | 23 | 18 | 15.8 | 15 | 17 | 0.87 | | | |
| 519 | hsa-miR-31-5p | MIMAT0000089 | | 14 | 14 | 21 | 12 | 19 | 16 | 10 | 18 | 19 | 15 | 15.8 | 15 | 16 | 0.94 | | | |
| 520 | hsa-miR-3180-3p | MIMAT0015058 | | 11 | 10 | 21 | 13 | 21 | 20 | 12 | 10 | 20 | 20 | 15.8 | 14 | 17 | 0.80 | | | |
| 521 | hsa-miR-489-3p | MIMAT0002805 | | 9 | 17 | 16 | 14 | 22 | 23 | 12 | 11 | 22 | 12 | 15.8 | 14 | 17 | 0.82 | | | |
| 522 | hsa-miR-649 | MIMAT0003319 | | 10 | 14 | 23 | 16 | 23 | 19 | 16 | 8 | 16 | 13 | 15.8 | 16 | 16 | 0.99 | | | |
| 523 | hsa-miR-891a-5p | MIMAT0004902 | | 7 | 16 | 13 | 19 | 18 | 17 | 14 | 11 | 23 | 20 | 15.8 | 14 | 17 | 0.80 | | | |
| 524 | hsa-miR-1264 | MIMAT0005791 | | 12 | 14 | 15 | 17 | 20 | 10 | 21 | 10 | 21 | 17 | 15.7 | 15 | 17 | 0.88 | | | |
| 525 | hsa-miR-1278 | MIMAT0005936 | | 11 | 18 | 11 | 14 | 21 | 17 | 17 | 9 | 15 | 24 | 15.7 | 14 | 17 | 0.79 | | | |
| 526 | hsa-miR-361-3p | MIMAT0004682 | | 15 | 16 | 24 | 17 | 12 | 18 | 10 | 9 | 18 | 18 | 15.7 | 18 | 14 | 1.27 | | | |
| 527 | hsa-miR-665 | MIMAT0004952 | | 21 | 21 | 19 | 12 | 15 | 15 | 13 | 10 | 23 | 8 | 15.7 | 18 | 14 | 1.30 | | | |
| 528 | hsa-miR-133a-3p | MIMAT0000427 | | 8 | 11 | 17 | 12 | 23 | 8 | 19 | 7 | 28 | 23 | 15.6 | 12 | 18 | 0.67 | | | |
| 529 | hsa-miR-133b | MIMAT0000770 | | 19 | 14 | 16 | 14 | 24 | 17 | 10 | 14 | 17 | 11 | 15.6 | 16 | 16 | 1.02 | | | |
| 530 | hsa-miR-5196-3p | MIMAT0021129 | | 15 | 19 | 19 | 4 | 17 | 22 | 19 | 9 | 20 | 12 | 15.6 | 14 | 17 | 0.86 | | hsa-miR-6732-3p | |
| 531 | hsa-miR-520e | MIMAT0002825 | | 14 | 13 | 20 | 13 | 34 | 12 | 10 | 13 | 14 | 13 | 15.6 | 15 | 16 | 0.94 | | | |
| 532 | hsa-miR-521 | MIMAT0002854 | | 8 | 19 | 21 | 13 | 15 | 12 | 16 | 12 | 30 | 10 | 15.6 | 15 | 16 | 0.96 | | | |
| 533 | hsa-miR-1293 | MIMAT0005883 | | 17 | 8 | 19 | 10 | 21 | 20 | 14 | 7 | 21 | 18 | 15.5 | 14 | 17 | 0.80 | | | |
| 534 | hsa-miR-33a-5p | MIMAT0000091 | | 15 | 28 | 11 | 11 | 17 | 13 | 16 | 6 | 24 | 14 | 15.5 | 16 | 15 | 1.08 | | | |
| 535 | hsa-miR-4431 | MIMAT0018947 | | 14 | 11 | 19 | 17 | 16 | 19 | 17 | 10 | 22 | 10 | 15.5 | 15 | 16 | 0.97 | | | |
| 536 | hsa-miR-675-5p | MIMAT0004284 | | 10 | 11 | 19 | 13 | 15 | 20 | 14 | 15 | 23 | 14 | 15.4 | 13 | 17 | 0.79 | | | |
| 537 | hsa-miR-1302 | MIMAT0005890 | | 18 | 17 | 14 | 20 | 10 | 14 | 16 | 12 | 23 | 9 | 15.3 | 17 | 14 | 1.23 | | | |
| 538 | hsa-miR-1287-3p | MIMAT0026738 | | 14 | 21 | 13 | 21 | 13 | 16 | 22 | 4 | 14 | 14 | 15.2 | 17 | 14 | 1.25 | | | |
| 539 | hsa-miR-30b-5p | MIMAT0000420 | | 19 | 22 | 13 | 12 | 13 | 20 | 12 | 14 | 17 | 10 | 15.2 | 17 | 14 | 1.15 | | | |
| 540 | hsa-miR-3613-3p | MIMAT0017991 | | 8 | 20 | 24 | 8 | 15 | 14 | 16 | 11 | 19 | 17 | 15.2 | 15 | 15 | 0.98 | | | |
| 541 | hsa-miR-518d-3p | MIMAT0002864 | | 14 | 19 | 18 | 15 | 17 | 14 | 11 | 12 | 21 | 11 | 15.2 | 17 | 14 | 1.15 | | | |
| 542 | hsa-miR-548b-3p | MIMAT0003254 | | 13 | 15 | 12 | 17 | 16 | 14 | 16 | 13 | 22 | 14 | 15.2 | 14 | 16 | 0.90 | | | |
| 543 | hsa-miR-580-3p | MIMAT0003245 | | 18 | 9 | 14 | 12 | 14 | 17 | 20 | 11 | 23 | 14 | 15.2 | 13 | 17 | 0.80 | | | |
| 544 | hsa-miR-614 | MIMAT0003282 | | 18 | 15 | 18 | 12 | 22 | 8 | 18 | 11 | 17 | 13 | 15.2 | 16 | 15 | 1.06 | | | |
| 545 | hsa-miR-296-3p | MIMAT0004679 | | 15 | 17 | 19 | 10 | 20 | 15 | 20 | 6 | 19 | 10 | 15.1 | 15 | 15 | 1.02 | | | |
| 546 | hsa-miR-513a-5p | MIMAT0002877 | | 11 | 18 | 25 | 10 | 9 | 21 | 17 | 5 | 24 | 11 | 15.1 | 16 | 15 | 1.10 | | | |
| 547 | hsa-miR-519b-3p | MIMAT0002837 | | 12 | 19 | 17 | 11 | 17 | 14 | 20 | 7 | 18 | 16 | 15.1 | 15 | 15 | 0.96 | | | |
| 548 | hsa-miR-135a-5p | MIMAT0000428 | | 31 | 9 | 24 | 1 | 15 | 25 | 19 | 4 | 16 | 6 | 15 | 16 | 14 | 1.15 | | | |
| 549 | hsa-miR-1915-3p | MIMAT0007892 | | 14 | 12 | 24 | 10 | 23 | 28 | 15 | 4 | 17 | 3 | 15 | 15 | 15 | 1.00 | | | |
| 550 | hsa-miR-3614-5p | MIMAT0017992 | | 17 | 11 | 15 | 17 | 19 | 12 | 14 | 7 | 20 | 18 | 15 | 15 | 15 | 1.00 | | | |
| 551 | hsa-miR-374c-5p | MIMAT0018443 | | 10 | 15 | 20 | 11 | 21 | 20 | 22 | 7 | 17 | 7 | 15 | 14 | 16 | 0.89 | | | |
| 552 | hsa-miR-488-3p | MIMAT0004763 | | 14 | 8 | 19 | 10 | 22 | 21 | 10 | 10 | 25 | 11 | 15 | 13 | 17 | 0.77 | | | |
| 553 | hsa-miR-1-3p | MIMAT0000416 | | 16 | 12 | 23 | 10 | 14 | 12 | 19 | 6 | 17 | 20 | 14.9 | 15 | 15 | 1.04 | | | |
| 554 | hsa-miR-431-5p | MIMAT0001625 | | 11 | 11 | 10 | 22 | 21 | 11 | 23 | 12 | 17 | 11 | 14.9 | 14 | 16 | 0.85 | | | |
| 555 | hsa-miR-1260b | MIMAT0015041 | | 22 | 21 | 21 | 7 | 16 | 16 | 16 | 7 | 12 | 10 | 14.8 | 18 | 13 | 1.38 | | | |
| 556 | hsa-miR-1976 | MIMAT0009451 | | 8 | 12 | 15 | 19 | 22 | 11 | 15 | 7 | 19 | 20 | 14.8 | 14 | 16 | 0.86 | | | |
| 557 | hsa-miR-3180-5p | MIMAT0015057 | | 8 | 8 | 12 | 16 | 19 | 19 | 20 | 16 | 22 | 8 | 14.8 | 11 | 17 | 0.63 | | | |
| 558 | hsa-miR-450a-2-3p | MIMAT0031074 | | 19 | 15 | 18 | 9 | 18 | 18 | 16 | 13 | 17 | 5 | 14.8 | 15 | 15 | 1.05 | | | |
| 559 | hsa-miR-499b-5p | MIMAT0019897 | | 16 | 14 | 14 | 7 | 17 | 18 | 9 | 7 | 36 | 10 | 14.8 | 13 | 16 | 0.79 | | | |
| 560 | hsa-miR-548c-5p | MIMAT0004806 | | 11 | 18 | 16 | 18 | 19 | 15 | 14 | 11 | 17 | 9 | 14.8 | 16 | 14 | 1.11 | | hsa-miR-548o-5p | hsa-miR-548am-5p |
| 561 | hsa-miR-187-3p | MIMAT0000262 | | 9 | 17 | 15 | 16 | 17 | 9 | 20 | 12 | 19 | 13 | 14.7 | 14 | 15 | 0.95 | | | |
| 562 | hsa-miR-196a-3p | MIMAT0004562 | | 9 | 16 | 18 | 14 | 13 | 15 | 11 | 8 | 22 | 21 | 14.7 | 14 | 15 | 0.95 | | | |
| 563 | hsa-miR-376b-3p | MIMAT0002172 | | 21 | 11 | 17 | 9 | 16 | 23 | 12 | 13 | 13 | 12 | 14.7 | 15 | 15 | 0.98 | | | |
| 564 | hsa-miR-4421 | MIMAT0018934 | | 10 | 15 | 18 | 11 | 14 | 18 | 12 | 7 | 30 | 12 | 14.7 | 14 | 16 | 0.87 | | | |
| 565 | hsa-miR-548e-3p | MIMAT0005874 | | 10 | 18 | 21 | 17 | 17 | 11 | 17 | 10 | 13 | 13 | 14.7 | 17 | 14 | 1.22 | | | |

| | | | | | | | | | | | | | | | | | | | |
|---|---|---|---|---|---|---|---|---|---|---|---|---|---|---|---|---|---|---|---|
| 566 | hsa-miR-616-3p | MIMAT0004805 | | 13 | 17 | 15 | 19 | 13 | 11 | 21 | 5 | 24 | 9 | 14.7 | 16 | 14 | 1.16 | | |
| 567 | hsa-miR-654-5p | MIMAT0003330 | | 9 | 14 | 12 | 11 | 13 | 23 | 18 | 7 | 26 | 14 | 14.7 | 12 | 17 | 0.68 | | |
| 568 | hsa-miR-342-5p | MIMAT0004694 | | 13 | 12 | 18 | 15 | 22 | 11 | 15 | 8 | 24 | 8 | 14.6 | 15 | 15 | 0.99 | | |
| 569 | hsa-miR-520g-3p | MIMAT0002858 | | 16 | 8 | 16 | 12 | 21 | 13 | 14 | 8 | 20 | 18 | 14.6 | 13 | 16 | 0.83 | | |
| 570 | hsa-miR-654-3p | MIMAT0004814 | | 9 | 18 | 16 | 12 | 18 | 17 | 11 | 8 | 24 | 13 | 14.6 | 14 | 15 | 0.91 | | |
| 571 | hsa-miR-885-5p | MIMAT0004947 | | 11 | 13 | 15 | 12 | 19 | 12 | 19 | 7 | 23 | 15 | 14.6 | 13 | 16 | 0.81 | | |
| 572 | hsa-miR-890 | MIMAT0004912 | | 12 | 13 | 15 | 16 | 17 | 16 | 13 | 6 | 24 | 14 | 14.6 | 14 | 15 | 0.93 | | |
| 573 | hsa-miR-1193 | MIMAT0015049 | | 15 | 17 | 17 | 15 | 15 | 15 | 14 | 8 | 19 | 10 | 14.5 | 16 | 14 | 1.19 | | |
| 574 | hsa-miR-1277-3p | MIMAT0005933 | | 14 | 12 | 27 | 14 | 11 | 12 | 13 | 11 | 14 | 17 | 14.5 | 17 | 13 | 1.29 | | |
| 575 | hsa-miR-339-5p | MIMAT0000764 | | 16 | 15 | 11 | 8 | 20 | 14 | 21 | 6 | 23 | 11 | 14.5 | 13 | 16 | 0.79 | | |
| 576 | hsa-miR-487a-3p | MIMAT0002178 | | 15 | 10 | 19 | 13 | 18 | 14 | 13 | 9 | 27 | 7 | 14.5 | 14 | 15 | 0.97 | | |
| 577 | hsa-miR-487b-5p | MIMAT0026614 | | 16 | 9 | 16 | 15 | 14 | 14 | 16 | 9 | 23 | 13 | 14.5 | 14 | 15 | 0.94 | | |
| 578 | hsa-miR-1252-5p | MIMAT0005944 | | 5 | 24 | 17 | 7 | 26 | 15 | 11 | 12 | 18 | 9 | 14.4 | 13 | 15 | 0.87 | | |
| 579 | hsa-miR-219a-1-3p | MIMAT0004567 | | 11 | 13 | 19 | 14 | 14 | 11 | 15 | 8 | 26 | 13 | 14.4 | 14 | 15 | 0.98 | | |
| 580 | hsa-miR-433-3p | MIMAT0001627 | | 16 | 16 | 27 | 10 | 21 | 14 | 2 | 11 | 16 | 11 | 14.4 | 17 | 13 | 1.38 | | |
| 581 | hsa-miR-4458 | MIMAT0018980 | | 13 | 17 | 26 | 11 | 10 | 11 | 16 | 9 | 19 | 12 | 14.4 | 17 | 13 | 1.31 | | |
| 582 | hsa-miR-5001-5p | MIMAT0021021 | | 7 | 17 | 20 | 12 | 15 | 20 | 13 | 11 | 12 | 17 | 14.4 | 14 | 15 | 0.95 | | |
| 583 | hsa-miR-885-3p | MIMAT0004948 | | 7 | 21 | 15 | 14 | 17 | 10 | 21 | 7 | 15 | 17 | 14.4 | 14 | 15 | 0.98 | | |
| 584 | hsa-miR-181c-5p | MIMAT0000258 | | 6 | 16 | 23 | 10 | 13 | 12 | 14 | 8 | 25 | 16 | 14.3 | 14 | 15 | 0.94 | | |
| 585 | hsa-miR-339-3p | MIMAT0004702 | | 12 | 16 | 19 | 11 | 15 | 12 | 14 | 12 | 16 | 16 | 14.3 | 15 | 14 | 1.02 | | |
| 586 | hsa-miR-624-3p | MIMAT0004807 | | 11 | 13 | 15 | 14 | 16 | 8 | 12 | 5 | 32 | 17 | 14.3 | 13 | 15 | 0.88 | | |
| 587 | hsa-miR-876-5p | MIMAT0004924 | | 10 | 19 | 15 | 18 | 16 | 14 | 11 | 11 | 18 | 11 | 14.3 | 16 | 14 | 1.15 | | |
| 588 | hsa-miR-3192-5p | MIMAT0015076 | | 11 | 15 | 17 | 11 | 18 | 15 | 14 | 11 | 15 | 15 | 14.2 | 14 | 15 | 0.92 | | |
| 589 | hsa-miR-6720-3p | MIMAT0025851 | | 9 | 26 | 22 | 13 | 13 | 12 | 16 | 7 | 15 | 9 | 14.2 | 18 | 12 | 1.46 | | |
| 590 | hsa-miR-188-3p | MIMAT0004613 | | 12 | 13 | 19 | 12 | 12 | 10 | 22 | 10 | 22 | 9 | 14.1 | 14 | 14 | 0.99 | | |
| 591 | hsa-miR-34b-3p | MIMAT0004676 | | 11 | 13 | 13 | 8 | 16 | 17 | 22 | 4 | 17 | 20 | 14.1 | 11 | 16 | 0.70 | | |
| 592 | hsa-miR-3918 | MIMAT0018192 | | 18 | 4 | 18 | 15 | 23 | 13 | 16 | 9 | 13 | 12 | 14.1 | 14 | 14 | 0.96 | | |
| 593 | hsa-miR-518c-3p | MIMAT0002848 | | 11 | 16 | 19 | 13 | 20 | 19 | 8 | 5 | 19 | 11 | 14.1 | 15 | 14 | 1.08 | | |
| 594 | hsa-miR-1224-3p | MIMAT0005459 | | 7 | 19 | 14 | 13 | 20 | 17 | 15 | 7 | 16 | 12 | 14 | 13 | 15 | 0.91 | | |
| 595 | hsa-miR-1250-5p | MIMAT0005902 | | 14 | 20 | 15 | 4 | 6 | 20 | 17 | 8 | 25 | 11 | 14 | 13 | 15 | 0.91 | | |
| 596 | hsa-miR-127-5p | MIMAT0004604 | | 15 | 15 | 9 | 14 | 13 | 16 | 15 | 10 | 21 | 12 | 14 | 13 | 15 | 0.91 | | |
| 597 | hsa-miR-4284 | MIMAT0016915 | | 13 | 11 | 18 | 19 | 12 | 13 | 15 | 12 | 15 | 12 | 14 | 15 | 13 | 1.16 | | |
| 598 | hsa-miR-617 | MIMAT0003286 | | 10 | 17 | 21 | 13 | 9 | 7 | 15 | 10 | 21 | 17 | 14 | 15 | 13 | 1.16 | | |
| 599 | hsa-miR-181a-2-3p | MIMAT0004558 | | 6 | 12 | 20 | 15 | 16 | 11 | 16 | 11 | 21 | 11 | 13.9 | 13 | 14 | 0.92 | | |
| 600 | hsa-miR-18b-5p | MIMAT0001412 | | 9 | 7 | 25 | 11 | 11 | 15 | 17 | 6 | 17 | 21 | 13.9 | 13 | 15 | 0.90 | | |
| 601 | hsa-miR-193b-3p | MIMAT0002819 | | 9 | 16 | 13 | 11 | 15 | 17 | 19 | 5 | 22 | 12 | 13.9 | 12 | 15 | 0.82 | | |
| 602 | hsa-miR-345-3p | MIMAT0022698 | | 17 | 21 | 7 | 14 | 16 | 12 | 14 | 5 | 17 | 16 | 13.9 | 15 | 13 | 1.11 | | |
| 603 | hsa-miR-887-3p | MIMAT0004951 | | 18 | 7 | 20 | 8 | 13 | 11 | 16 | 10 | 20 | 16 | 13.9 | 13 | 14 | 0.92 | | |
| 604 | hsa-miR-3605-5p | MIMAT0017981 | | 9 | 15 | 19 | 17 | 17 | 8 | 15 | 10 | 14 | 14 | 13.8 | 15 | 13 | 1.15 | | |
| 605 | hsa-miR-381-5p | MIMAT0022862 | | 13 | 17 | 12 | 15 | 19 | 12 | 12 | 10 | 20 | 8 | 13.8 | 14 | 14 | 1.06 | | |
| 606 | hsa-miR-490-5p | MIMAT0004764 | | 15 | 18 | 18 | 10 | 13 | 16 | 10 | 7 | 23 | 8 | 13.8 | 15 | 13 | 1.19 | | |
| 607 | hsa-miR-655-3p | MIMAT0003331 | | 17 | 12 | 7 | 16 | 14 | 14 | 20 | 6 | 20 | 12 | 13.8 | 13 | 14 | 0.91 | | |
| 608 | hsa-miR-934 | MIMAT0004977 | | 14 | 14 | 12 | 9 | 19 | 11 | 12 | 9 | 22 | 16 | 13.8 | 12 | 15 | 0.83 | | |
| 609 | hsa-miR-151b | MIMAT0010214 | | 13 | 13 | 17 | 15 | 23 | 12 | 10 | 3 | 18 | 13 | 13.7 | 15 | 13 | 1.10 | | |
| 610 | hsa-miR-190a-5p | MIMAT0000458 | | 10 | 16 | 17 | 12 | 17 | 14 | 13 | 3 | 21 | 14 | 13.7 | 14 | 14 | 1.01 | | |
| 611 | hsa-miR-211-3p | MIMAT0022694 | | 13 | 18 | 16 | 6 | 8 | 22 | 18 | 5 | 23 | 8 | 13.7 | 13 | 14 | 0.95 | | |
| 612 | hsa-miR-323a-3p | MIMAT0000755 | | 7 | 12 | 17 | 8 | 24 | 16 | 12 | 9 | 20 | 12 | 13.7 | 11 | 16 | 0.71 | | |
| 613 | hsa-miR-517b-3p | MIMAT0002857 | | 11 | 5 | 14 | 9 | 24 | 7 | 19 | 12 | 20 | 16 | 13.7 | 10 | 16 | 0.60 | | |
| 614 | hsa-miR-541-3p | MIMAT0004920 | | 8 | 7 | 13 | 15 | 20 | 17 | 17 | 6 | 21 | 13 | 13.7 | 11 | 16 | 0.69 | | |
| 615 | hsa-miR-604 | MIMAT0003272 | | 11 | 11 | 17 | 16 | 17 | 15 | 13 | 13 | 17 | 7 | 13.7 | 14 | 14 | 1.01 | | |
| 616 | hsa-miR-103a-3p | MIMAT0000101 | | 13 | 12 | 10 | 12 | 19 | 10 | 20 | 15 | 18 | 7 | 13.6 | 12 | 15 | 0.79 | | |
| 617 | hsa-miR-1269b | MIMAT0019059 | | 14 | 11 | 10 | 11 | 18 | 12 | 21 | 7 | 21 | 11 | 13.6 | 12 | 15 | 0.77 | | |
| 618 | hsa-miR-150-5p | MIMAT0000451 | | 8 | 13 | 16 | 12 | 17 | 15 | 13 | 9 | 18 | 15 | 13.6 | 12 | 15 | 0.84 | | |
| 619 | hsa-miR-330-5p | MIMAT0004693 | | 13 | 19 | 12 | 6 | 15 | 14 | 15 | 8 | 20 | 14 | 13.6 | 13 | 14 | 0.87 | | |
| 620 | hsa-miR-4425 | MIMAT0018940 | | 18 | 12 | 16 | 8 | 13 | 11 | 12 | 9 | 18 | 19 | 13.6 | 14 | 14 | 0.99 | | |
| 621 | hsa-miR-371b-5p | MIMAT0019892 | | 13 | 10 | 19 | 14 | 11 | 15 | 18 | 7 | 16 | 12 | 13.5 | 14 | 13 | 1.06 | | |
| 622 | hsa-miR-874-3p | MIMAT0004911 | | 14 | 7 | 19 | 9 | 19 | 10 | 13 | 3 | 26 | 15 | 13.5 | 12 | 14 | 0.85 | | |
| 623 | hsa-miR-101-3p | MIMAT0000099 | | 11 | 11 | 16 | 14 | 14 | 6 | 10 | 9 | 29 | 14 | 13.4 | 13 | 14 | 0.95 | | |
| 624 | hsa-miR-196b-5p | MIMAT0001080 | | 17 | 14 | 15 | 11 | 10 | 17 | 14 | 8 | 20 | 8 | 13.4 | 14 | 13 | 1.11 | | |
| 625 | hsa-miR-518e-3p | MIMAT0002861 | | 7 | 8 | 16 | 13 | 18 | 14 | 23 | 11 | 13 | 11 | 13.4 | 11 | 15 | 0.73 | | |
| 626 | hsa-miR-1202 | MIMAT0005865 | | 12 | 11 | 13 | 13 | 16 | 18 | 17 | 10 | 18 | 5 | 13.3 | 12 | 14 | 0.88 | | |
| 627 | hsa-miR-3195 | MIMAT0015079 | | 12 | 15 | 14 | 14 | 16 | 13 | 14 | 10 | 14 | 11 | 13.3 | 14 | 13 | 1.06 | | |
| 628 | hsa-miR-3690 | MIMAT0018119 | | 14 | 11 | 11 | 5 | 22 | 17 | 12 | 6 | 18 | 17 | 13.3 | 10 | 15 | 0.67 | | |
| 629 | hsa-miR-452-5p | MIMAT0001635 | | 16 | 11 | 20 | 13 | 12 | 13 | 13 | 10 | 16 | 9 | 13.3 | 15 | 12 | 1.23 | | |
| 630 | hsa-miR-516b-5p | MIMAT0002859 | | 7 | 11 | 16 | 15 | 18 | 12 | 20 | 9 | 16 | 9 | 13.3 | 12 | 14 | 0.88 | | |
| 631 | hsa-miR-1226-3p | MIMAT0005577 | | 10 | 14 | 12 | 9 | 14 | 12 | 15 | 8 | 22 | 16 | 13.2 | 11 | 15 | 0.78 | | |
| 632 | hsa-miR-153-3p | MIMAT0000439 | | 7 | 18 | 12 | 13 | 15 | 10 | 17 | 9 | 14 | 17 | 13.2 | 13 | 14 | 0.91 | | |
| 633 | hsa-miR-223-3p | MIMAT0000280 | | 14 | 15 | 16 | 9 | 11 | 10 | 16 | 11 | 19 | 11 | 13.2 | 14 | 13 | 1.04 | | |
| 634 | hsa-miR-326 | MIMAT0000756 | | 10 | 15 | 12 | 22 | 9 | 10 | 12 | 12 | 23 | 7 | 13.2 | 15 | 12 | 1.21 | | |
| 635 | hsa-miR-362-3p | MIMAT0004683 | | 10 | 11 | 21 | 15 | 13 | 14 | 9 | 8 | 18 | 13 | 13.2 | 14 | 13 | 1.14 | | |
| 636 | hsa-miR-378c | MIMAT0016847 | | 6 | 13 | 15 | 10 | 8 | 13 | 17 | 7 | 23 | 20 | 13.2 | 11 | 15 | 0.75 | | |
| 637 | hsa-miR-556-3p | MIMAT0004793 | | 17 | 15 | 16 | 13 | 14 | 16 | 2 | 8 | 19 | 12 | 13.2 | 15 | 12 | 1.29 | | |
| 638 | hsa-miR-1273c | MIMAT0015017 | | 20 | 13 | 8 | 5 | 15 | 14 | 14 | 10 | 17 | 15 | 13.1 | 12 | 14 | 0.81 | | |
| 639 | hsa-miR-2682-5p | MIMAT0013517 | | 9 | 13 | 13 | 12 | 11 | 18 | 11 | 10 | 24 | 10 | 13.1 | 12 | 14 | 0.84 | | |
| 640 | hsa-miR-3916 | MIMAT0018190 | | 16 | 14 | 18 | 17 | 12 | 10 | 11 | 10 | 14 | 9 | 13.1 | 16 | 11 | 1.48 | | |
| 641 | hsa-miR-562 | MIMAT0003226 | | 9 | 16 | 22 | 5 | 15 | 10 | 17 | 10 | 15 | 12 | 13.1 | 13 | 13 | 0.99 | | |
| 642 | hsa-miR-384 | MIMAT0001075 | | 7 | 7 | 20 | 12 | 20 | 10 | 14 | 11 | 21 | 7 | 12.9 | 12 | 14 | 0.83 | | |
| 643 | hsa-miR-503-3p | MIMAT0022925 | | 15 | 11 | 19 | 8 | 17 | 10 | 11 | 7 | 21 | 10 | 12.9 | 13 | 13 | 1.05 | | |
| 644 | hsa-miR-578 | MIMAT0003243 | | 8 | 13 | 23 | 10 | 18 | 16 | 11 | 7 | 12 | 11 | 12.9 | 14 | 13 | 1.08 | | |
| 645 | hsa-miR-1244 | MIMAT0005896 | | 8 | 17 | 21 | 6 | 16 | 10 | 8 | 10 | 19 | 13 | 12.8 | 13 | 13 | 1.03 | | |
| 646 | hsa-miR-129-5p | MIMAT0000242 | | 17 | 13 | 22 | 10 | 14 | 18 | 11 | 5 | 10 | 8 | 12.8 | 16 | 11 | 1.41 | | |
| 647 | hsa-miR-450a-1-3p | MIMAT0022700 | | 9 | 19 | 13 | 9 | 14 | 18 | 18 | 2 | 17 | 9 | 12.8 | 13 | 13 | 0.96 | | |
| 648 | hsa-miR-1304-3p | MIMAT0022720 | | 11 | 14 | 15 | 11 | 15 | 14 | 11 | 6 | 16 | 14 | 12.7 | 13 | 13 | 1.01 | | |
| 649 | hsa-miR-378b | MIMAT0014999 | | 11 | 13 | 16 | 12 | 14 | 10 | 7 | 9 | 21 | 14 | 12.7 | 13 | 13 | 1.04 | | |
| 650 | hsa-miR-515-3p | MIMAT0002827 | | 9 | 15 | 19 | 8 | 13 | 14 | 13 | 9 | 15 | 12 | 12.7 | 13 | 13 | 1.01 | | |
| 651 | hsa-miR-1271-3p | MIMAT0022712 | | 13 | 14 | 15 | 6 | 14 | 13 | 9 | 6 | 17 | 19 | 12.6 | 12 | 13 | 0.92 | | |
| 652 | hsa-miR-134-5p | MIMAT0000447 | | 9 | 13 | 15 | 8 | 14 | 9 | 13 | 7 | 21 | 17 | 12.6 | 11 | 14 | 0.83 | | hsa-miR-6728-5p |
| 653 | hsa-miR-3150b-3p | MIMAT0018194 | | 13 | 12 | 18 | 9 | 16 | 18 | 12 | 6 | 16 | 6 | 12.6 | 13 | 12 | 1.05 | | |
| 654 | hsa-miR-381-3p | MIMAT0000736 | | 11 | 10 | 16 | 14 | 17 | 14 | 14 | 4 | 14 | 12 | 12.6 | 13 | 13 | 1.02 | | |
| 655 | hsa-miR-425-5p | MIMAT0003393 | | 14 | 13 | 21 | 8 | 7 | 15 | 12 | 12 | 14 | 10 | 12.6 | 14 | 12 | 1.20 | | |
| 656 | hsa-miR-449c-5p | MIMAT0010251 | | 11 | 16 | 10 | 14 | 15 | 10 | 13 | 5 | 22 | 10 | 12.6 | 13 | 13 | 1.02 | | |
| 657 | hsa-miR-487b-3p | MIMAT0003180 | | 20 | 6 | 10 | 8 | 14 | 17 | 10 | 6 | 21 | 14 | 12.6 | 11 | 14 | 0.80 | | |
| 658 | hsa-miR-514a-5p | MIMAT0022702 | | 9 | 11 | 15 | 12 | 15 | 9 | 15 | 8 | 22 | 10 | 12.6 | 12 | 13 | 0.89 | | |
| 659 | hsa-miR-208b-5p | MIMAT0026722 | | 10 | 17 | 17 | 16 | 13 | 8 | 18 | 3 | 13 | 10 | 12.5 | 15 | 11 | 1.38 | | |
| 660 | hsa-miR-320b | MIMAT0005792 | | 8 | 17 | 11 | 16 | 12 | 15 | 11 | 7 | 15 | 13 | 12.5 | 13 | 12 | 1.07 | | |

| | | | | | | | | | | | | | | | | | | | |
|---|---|---|---|---|---|---|---|---|---|---|---|---|---|---|---|---|---|---|---|
| 661 | hsa-miR-324-5p | MIMAT0000761 | | 12 | 12 | 19 | 12 | 8 | 15 | 14 | 5 | 20 | 8 | 12.5 | 14 | 12 | 1.18 | | |
| 662 | hsa-miR-4741 | MIMAT0019871 | | 16 | 15 | 18 | 10 | 16 | 9 | 11 | 9 | 12 | 9 | 12.5 | 15 | 11 | 1.34 | | |
| 663 | hsa-miR-507 | MIMAT0002879 | | 10 | 11 | 23 | 7 | 10 | 11 | 14 | 11 | 13 | 15 | 12.5 | 13 | 12 | 1.03 | | |
| 664 | hsa-miR-1275 | MIMAT0005929 | | 12 | 14 | 15 | 8 | 10 | 13 | 20 | 11 | 14 | 7 | 12.4 | 12 | 13 | 0.98 | | |
| 665 | hsa-miR-3182 | MIMAT0015062 | | 20 | 10 | 13 | 9 | 9 | 12 | 15 | 6 | 13 | 17 | 12.4 | 13 | 12 | 1.08 | | |
| 666 | hsa-miR-606 | MIMAT0003274 | | 9 | 15 | 19 | 5 | 15 | 12 | 11 | 5 | 20 | 13 | 12.4 | 12 | 13 | 0.95 | | |
| 667 | hsa-miR-1296-5p | MIMAT0005794 | | 12 | 19 | 14 | 13 | 13 | 8 | 10 | 8 | 15 | 11 | 12.3 | 15 | 11 | 1.34 | | |
| 668 | hsa-miR-154-5p | MIMAT0000452 | | 12 | 8 | 17 | 10 | 13 | 10 | 17 | 7 | 18 | 11 | 12.3 | 12 | 13 | 0.93 | | |
| 669 | hsa-miR-4532 | MIMAT0019071 | | 11 | 10 | 18 | 10 | 16 | 7 | 12 | 4 | 22 | 13 | 12.3 | 12 | 12 | 0.99 | | |
| 670 | hsa-miR-660-5p | MIMAT0003338 | | 6 | 14 | 18 | 17 | 15 | 11 | 6 | 7 | 23 | 6 | 12.3 | 14 | 11 | 1.21 | | |
| 671 | hsa-miR-151a-5p | MIMAT0004697 | | 7 | 11 | 19 | 7 | 13 | 16 | 19 | 4 | 19 | 7 | 12.2 | 11 | 13 | 0.85 | | |
| 672 | hsa-miR-370-5p | MIMAT0026483 | | 10 | 9 | 13 | 8 | 15 | 12 | 17 | 6 | 19 | 13 | 12.2 | 10 | 14 | 0.73 | | |
| 673 | hsa-miR-708-5p | MIMAT0004926 | | 8 | 15 | 10 | 12 | 21 | 13 | 8 | 14 | 15 | 6 | 12.2 | 11 | 13 | 0.88 | | |
| 674 | hsa-miR-92b-3p | MIMAT0003218 | | 11 | 12 | 10 | 12 | 10 | 11 | 17 | 10 | 17 | 12 | 12.2 | 11 | 13 | 0.88 | | |
| 675 | hsa-miR-181d-3p | MIMAT0026608 | | 9 | 14 | 20 | 11 | 13 | 10 | 19 | 3 | 13 | 9 | 12.1 | 14 | 11 | 1.21 | | |
| 676 | hsa-miR-935 | MIMAT0004978 | | 10 | 10 | 13 | 15 | 13 | 5 | 12 | 10 | 16 | 17 | 12.1 | 12 | 12 | 0.99 | | |
| 677 | hsa-miR-1276 | MIMAT0005930 | | 12 | 10 | 19 | 11 | 17 | 12 | 9 | 3 | 18 | 9 | 12 | 13 | 11 | 1.15 | | |
| 678 | hsa-miR-1291 | MIMAT0005881 | | 12 | 11 | 12 | 18 | 15 | 10 | 16 | 7 | 11 | 8 | 12 | 13 | 11 | 1.19 | | |
| 679 | hsa-miR-1908-3p | MIMAT0026916 | | 10 | 11 | 15 | 16 | 17 | 12 | 7 | 2 | 19 | 11 | 12 | 13 | 11 | 1.15 | | |
| 680 | hsa-miR-1910-5p | MIMAT0007884 | | 6 | 13 | 13 | 13 | 7 | 19 | 19 | 9 | 13 | 8 | 12 | 11 | 13 | 0.90 | | |
| 681 | hsa-miR-3065-3p | MIMAT0015378 | | 9 | 6 | 11 | 18 | 20 | 12 | 12 | 8 | 14 | 10 | 12 | 11 | 13 | 0.87 | | |
| 682 | hsa-miR-329-3p | MIMAT0001629 | | 14 | 14 | 10 | 10 | 10 | 12 | 16 | 3 | 20 | 11 | 12 | 12 | 12 | 1.00 | | |
| 683 | hsa-miR-501-3p | MIMAT0004774 | | 7 | 15 | 10 | 10 | 9 | 19 | 11 | 9 | 17 | 13 | 12 | 11 | 13 | 0.81 | | |
| 684 | hsa-miR-544a | MIMAT0003164 | | 9 | 13 | 20 | 11 | 12 | 14 | 12 | 6 | 16 | 7 | 12 | 13 | 11 | 1.19 | | |
| 685 | hsa-miR-6503-3p | MIMAT0025463 | | 6 | 18 | 11 | 9 | 10 | 11 | 8 | 9 | 23 | 15 | 12 | 11 | 13 | 0.87 | | |
| 686 | hsa-miR-345-5p | MIMAT0000772 | | 7 | 8 | 16 | 15 | 10 | 11 | 12 | 7 | 21 | 12 | 11.9 | 12 | 12 | 0.95 | | |
| 687 | hsa-miR-455-3p | MIMAT0004784 | | 6 | 17 | 14 | 7 | 13 | 9 | 15 | 10 | 19 | 9 | 11.9 | 11 | 13 | 0.88 | | |
| 688 | hsa-miR-524-3p | MIMAT0002850 | | 13 | 14 | 12 | 7 | 10 | 7 | 17 | 10 | 16 | 13 | 11.9 | 12 | 12 | 0.95 | | |
| 689 | hsa-miR-128-2-5p | MIMAT0031095 | | 7 | 10 | 15 | 8 | 11 | 8 | 16 | 9 | 22 | 11 | 11.7 | 10 | 13 | 0.78 | | |
| 690 | hsa-miR-182-3p | MIMAT0000260 | | 7 | 11 | 14 | 11 | 16 | 8 | 12 | 9 | 16 | 13 | 11.7 | 11 | 12 | 0.87 | | |
| 691 | hsa-miR-369-5p | MIMAT0001621 | | 9 | 12 | 16 | 12 | 14 | 12 | 14 | 9 | 10 | 9 | 11.7 | 12 | 11 | 1.08 | | |
| 692 | hsa-miR-490-3p | MIMAT0002806 | | 10 | 10 | 13 | 10 | 17 | 12 | 16 | 11 | 8 | 10 | 11.7 | 11 | 12 | 0.87 | | |
| 693 | hsa-miR-1469 | MIMAT0007347 | | 9 | 10 | 12 | 9 | 15 | 13 | 10 | 5 | 16 | 17 | 11.6 | 10 | 13 | 0.79 | | |
| 694 | hsa-miR-3185 | MIMAT0015065 | | 14 | 18 | 8 | 9 | 11 | 11 | 12 | 4 | 13 | 16 | 11.6 | 12 | 11 | 1.10 | | |
| 695 | hsa-miR-3196 | MIMAT0015080 | | 10 | 10 | 10 | 10 | 19 | 11 | 15 | 4 | 21 | 6 | 11.6 | 10 | 13 | 0.79 | | |
| 696 | hsa-miR-409-5p | MIMAT0001638 | | 12 | 14 | 11 | 9 | 15 | 8 | 18 | 4 | 7 | 18 | 11.6 | 12 | 12 | 0.99 | | |
| 697 | hsa-miR-516a-5p | MIMAT0004770 | | 10 | 13 | 13 | 8 | 12 | 10 | 11 | 9 | 18 | 12 | 11.6 | 11 | 12 | 0.92 | | |
| 698 | hsa-miR-520a-3p | MIMAT0002834 | | 9 | 12 | 15 | 11 | 9 | 11 | 13 | 6 | 18 | 12 | 11.6 | 12 | 12 | 1.02 | | |
| 699 | hsa-miR-130b-3p | MIMAT0000691 | | 13 | 11 | 10 | 9 | 14 | 12 | 13 | 4 | 19 | 10 | 11.5 | 11 | 12 | 0.90 | | |
| 700 | hsa-miR-210-5p | MIMAT0026475 | | 12 | 8 | 9 | 16 | 12 | 9 | 11 | 10 | 15 | 13 | 11.5 | 11 | 12 | 0.96 | | |
| 701 | hsa-miR-513c-3p | MIMAT0022728 | | 10 | 12 | 11 | 9 | 18 | 12 | 11 | 9 | 16 | 7 | 11.5 | 11 | 12 | 0.86 | | |
| 702 | hsa-miR-449a | MIMAT0001541 | | 8 | 14 | 10 | 14 | 19 | 16 | 10 | 7 | 9 | 7 | 11.4 | 12 | 11 | 1.01 | | |
| 703 | hsa-miR-600 | MIMAT0003268 | | 6 | 11 | 23 | 9 | 19 | 9 | 11 | 4 | 12 | 10 | 11.4 | 12 | 11 | 1.13 | | |
| 704 | hsa-miR-1204 | MIMAT0005868 | | 9 | 11 | 9 | 13 | 14 | 16 | 11 | 5 | 15 | 10 | 11.3 | 11 | 12 | 0.89 | | |
| 705 | hsa-miR-1288-3p | MIMAT0005942 | | 6 | 11 | 18 | 6 | 15 | 20 | 10 | 6 | 16 | 5 | 11.3 | 10 | 12 | 0.85 | | |
| 706 | hsa-miR-372-3p | MIMAT0000724 | | 6 | 11 | 16 | 9 | 17 | 7 | 9 | 7 | 18 | 13 | 11.3 | 11 | 12 | 0.89 | | |
| 707 | hsa-miR-221-5p | MIMAT0004568 | | 16 | 14 | 19 | 5 | 8 | 18 | 5 | 6 | 14 | 7 | 11.2 | 14 | 10 | 1.40 | | |
| 708 | hsa-miR-365b-5p | MIMAT0022833 | | 10 | 9 | 17 | 10 | 15 | 13 | 4 | 7 | 12 | 15 | 11.2 | 12 | 11 | 1.05 | | |
| 709 | hsa-miR-373-3p | MIMAT0000726 | | 16 | 10 | 7 | 11 | 18 | 9 | 6 | 8 | 17 | 10 | 11.2 | 11 | 11 | 0.97 | | |
| 710 | hsa-miR-651-3p | MIMAT0026624 | | 10 | 16 | 19 | 12 | 14 | 10 | 6 | 4 | 12 | 9 | 11.2 | 14 | 9 | 1.55 | | |
| 711 | hsa-miR-331-5p | MIMAT0004700 | | 8 | 9 | 15 | 6 | 15 | 10 | 9 | 8 | 16 | 15 | 11.1 | 10 | 12 | 0.78 | | |
| 712 | hsa-miR-4524a-5p | MIMAT0019062 | | 8 | 17 | 12 | 8 | 11 | 12 | 16 | 4 | 15 | 8 | 11.1 | 11 | 11 | 1.02 | | |
| 713 | hsa-miR-519e-3p | MIMAT0002829 | | 12 | 12 | 4 | 11 | 12 | 18 | 11 | 5 | 15 | 11 | 11.1 | 10 | 12 | 0.81 | | |
| 714 | hsa-miR-619-3p | MIMAT0003288 | | 10 | 11 | 13 | 12 | 16 | 13 | 7 | 7 | 15 | 7 | 11.1 | 12 | 11 | 1.06 | | |
| 715 | hsa-miR-1909-3p | MIMAT0007883 | | 10 | 13 | 9 | 10 | 17 | 11 | 11 | 11 | 10 | 8 | 11 | 11 | 11 | 0.93 | | |
| 716 | hsa-miR-505-3p | MIMAT0002876 | | 11 | 9 | 17 | 6 | 6 | 15 | 11 | 4 | 17 | 14 | 11 | 11 | 11 | 0.96 | | |
| 717 | hsa-miR-550a-5p | MIMAT0004800 | | 10 | 22 | 8 | 12 | 10 | 8 | 12 | 6 | 12 | 10 | 11 | 13 | 10 | 1.34 | | |
| 718 | hsa-miR-1205 | MIMAT0005869 | | 4 | 13 | 15 | 8 | 12 | 16 | 9 | 10 | 13 | 9 | 10.9 | 10 | 12 | 0.87 | | |
| 719 | hsa-miR-1260a | MIMAT0005911 | | 14 | 10 | 14 | 4 | 17 | 14 | 10 | 7 | 12 | 7 | 10.9 | 11 | 11 | 0.94 | | |
| 720 | hsa-miR-548h-5p | MIMAT0005928 | | 7 | 13 | 19 | 15 | 12 | 6 | 4 | 8 | 17 | 8 | 10.9 | 14 | 9 | 1.47 | | |
| 721 | hsa-miR-204-5p | MIMAT0000265 | | 19 | 9 | 10 | 10 | 11 | 14 | 10 | 6 | 12 | 7 | 10.8 | 12 | 10 | 1.20 | | |
| 722 | hsa-miR-3158-3p | MIMAT0015032 | | 11 | 12 | 11 | 2 | 10 | 20 | 13 | 4 | 12 | 13 | 10.8 | 9 | 12 | 0.75 | | |
| 723 | hsa-miR-6511a-5p | MIMAT0025478 | | 8 | 14 | 15 | 11 | 9 | 15 | 16 | 2 | 13 | 5 | 10.8 | 12 | 10 | 1.20 | | |
| 724 | hsa-miR-297 | MIMAT0004450 | | 9 | 19 | 11 | 8 | 12 | 12 | 11 | 4 | 13 | 8 | 10.7 | 12 | 10 | 1.18 | | |
| 725 | hsa-miR-320c | MIMAT0005793 | | 11 | 8 | 10 | 13 | 12 | 10 | 10 | 3 | 18 | 12 | 10.7 | 11 | 11 | 0.97 | | |
| 726 | hsa-miR-376c-5p | MIMAT0022861 | | 10 | 11 | 16 | 10 | 11 | 6 | 12 | 4 | 15 | 12 | 10.7 | 12 | 10 | 1.18 | | |
| 727 | hsa-miR-500a-5p | MIMAT0004773 | | 10 | 8 | 15 | 9 | 12 | 18 | 8 | 4 | 13 | 10 | 10.7 | 11 | 11 | 0.97 | | hsa-miR-501-5p |
| 728 | hsa-miR-940 | MIMAT0004983 | | 11 | 9 | 11 | 11 | 10 | 11 | 11 | 7 | 15 | 11 | 10.7 | 11 | 11 | 0.97 | | |
| 729 | hsa-miR-892b | MIMAT0004918 | | 6 | 11 | 18 | 7 | 14 | 6 | 12 | 9 | 15 | 8 | 10.6 | 11 | 11 | 0.98 | | |
| 730 | hsa-miR-3613-5p | MIMAT0017990 | | 5 | 11 | 6 | 12 | 15 | 8 | 14 | 9 | 16 | 9 | 10.5 | 9 | 12 | 0.72 | | |
| 731 | hsa-miR-671-3p | MIMAT0004819 | | 8 | 9 | 7 | 14 | 14 | 5 | 14 | 11 | 17 | 6 | 10.5 | 10 | 11 | 0.85 | | |
| 732 | hsa-miR-504-3p | MIMAT0026612 | | 15 | 9 | 9 | 10 | 17 | 7 | 9 | 8 | 10 | 10 | 10.4 | 11 | 10 | 1.06 | | |
| 733 | hsa-miR-539-3p | MIMAT0022705 | | 8 | 6 | 10 | 8 | 14 | 6 | 12 | 16 | 15 | 9 | 10.4 | 8 | 12 | 0.67 | | |
| 734 | hsa-miR-767-3p | MIMAT0003883 | | 8 | 12 | 9 | 9 | 15 | 12 | 8 | 7 | 13 | 11 | 10.4 | 10 | 11 | 0.86 | | |
| 735 | hsa-miR-146b-3p | MIMAT0004766 | | 13 | 15 | 7 | 10 | 17 | 10 | 8 | 6 | 11 | 6 | 10.3 | 11 | 10 | 1.16 | | |
| 736 | hsa-miR-3190-3p | MIMAT0022839 | | 13 | 15 | 11 | 9 | 12 | 10 | 6 | 4 | 14 | 9 | 10.3 | 12 | 9 | 1.31 | | |
| 737 | hsa-miR-652-5p | MIMAT0022709 | | 5 | 18 | 19 | 4 | 12 | 7 | 10 | 4 | 18 | 6 | 10.3 | 12 | 10 | 1.21 | | |
| 738 | hsa-miR-330-3p | MIMAT0000751 | | 6 | 8 | 12 | 4 | 14 | 12 | 18 | 11 | 14 | 3 | 10.2 | 8 | 12 | 0.63 | | |
| 739 | hsa-miR-4448 | MIMAT0018967 | | 13 | 8 | 11 | 16 | 15 | 4 | 6 | 7 | 16 | 6 | 10.2 | 12 | 9 | 1.33 | | |
| 740 | hsa-miR-492 | MIMAT0002812 | | 12 | 14 | 8 | 10 | 9 | 9 | 7 | 5 | 19 | 9 | 10.2 | 11 | 10 | 1.14 | | |
| 741 | hsa-miR-494-5p | MIMAT0026607 | | 10 | 12 | 11 | 11 | 11 | 9 | 10 | 5 | 14 | 9 | 10.2 | 11 | 10 | 1.14 | | |
| 742 | hsa-miR-652-3p | MIMAT0003322 | | 5 | 9 | 12 | 9 | 14 | 10 | 10 | 6 | 17 | 10 | 10.2 | 9 | 11 | 0.78 | | |
| 743 | hsa-miR-1306-5p | MIMAT0022726 | | 12 | 11 | 14 | 5 | 9 | 13 | 14 | 4 | 16 | 3 | 10.1 | 11 | 10 | 1.07 | | |
| 744 | hsa-miR-193a-3p | MIMAT0000459 | | 9 | 12 | 13 | 8 | 13 | 14 | 5 | 7 | 14 | 6 | 10.1 | 11 | 10 | 1.07 | | |
| 745 | hsa-miR-3140-5p | MIMAT0019204 | | 9 | 13 | 17 | 6 | 7 | 10 | 10 | 4 | 12 | 13 | 10.1 | 11 | 9 | 1.21 | | |
| 746 | hsa-miR-758-5p | MIMAT0022929 | | 10 | 16 | 12 | 8 | 14 | 6 | 6 | 3 | 20 | 6 | 10.1 | 12 | 9 | 1.25 | | |
| 747 | hsa-miR-512-3p | MIMAT0002823 | | 9 | 16 | 9 | 10 | 9 | 10 | 12 | 8 | 9 | 8 | 10 | 11 | 9 | 1.18 | | |
| 748 | hsa-miR-664b-5p | MIMAT0022271 | | 8 | 12 | 8 | 6 | 10 | 10 | 15 | 7 | 17 | 7 | 10 | 9 | 11 | 0.77 | | |
| 749 | hsa-miR-769-3p | MIMAT0003887 | | 12 | 9 | 9 | 8 | 13 | 12 | 11 | 6 | 8 | 12 | 10 | 10 | 10 | 0.92 | | |
| 750 | hsa-miR-942-3p | MIMAT0026734 | | 5 | 10 | 19 | 7 | 13 | 11 | 7 | 6 | 14 | 8 | 10 | 10 | 10 | 1.04 | | |
| 751 | hsa-miR-1245b-3p | MIMAT0019951 | | 15 | 8 | 12 | 7 | 13 | 7 | 17 | 3 | 10 | 7 | 9.9 | 11 | 10 | 1.11 | | |
| 752 | hsa-miR-570-3p | MIMAT0003235 | | 1 | 18 | 10 | 1 | 17 | 19 | 1 | 1 | 10 | 21 | 9.9 | 8 | 12 | 0.65 | | |
| 753 | hsa-miR-631 | MIMAT0003300 | | 5 | 12 | 7 | 8 | 16 | 10 | 9 | 6 | 13 | 13 | 9.9 | 8 | 11 | 0.72 | | |
| 754 | hsa-miR-1249-5p | MIMAT0032029 | | 7 | 13 | 18 | 8 | 6 | 6 | 13 | 7 | 13 | 7 | 9.8 | 12 | 9 | 1.33 | | |
| 755 | hsa-miR-383-5p | MIMAT0000738 | | 3 | 9 | 13 | 10 | 17 | 8 | 9 | 6 | 10 | 12 | 9.7 | 9 | 10 | 0.85 | | |

| | | | | | | | | | | | | | | | | | | | | |
|---|---|---|---|---|---|---|---|---|---|---|---|---|---|---|---|---|---|---|---|---|
| 756 | hsa-miR-1301-3p | MIMAT0005797 | | 8 | 3 | 6 | 12 | 12 | 7 | 10 | 5 | 24 | 8 | 9.5 | 7 | 11 | 0.66 | | | |
| 757 | hsa-miR-323a-5p | MIMAT0004696 | | 6 | 12 | 13 | 9 | 8 | 13 | 11 | 5 | 11 | 7 | 9.5 | 10 | 9 | 1.09 | | | |
| 758 | hsa-miR-510-5p | MIMAT0002882 | | 12 | 7 | 14 | 8 | 9 | 15 | 7 | 2 | 11 | 10 | 9.5 | 10 | 9 | 1.14 | | | |
| 759 | hsa-miR-637 | MIMAT0003307 | | 9 | 8 | 7 | 10 | 10 | 5 | 6 | 10 | 16 | 14 | 9.5 | 9 | 10 | 0.84 | | | |
| 760 | hsa-miR-1233-3p | MIMAT0005588 | | 8 | 8 | 10 | 3 | 13 | 10 | 13 | 5 | 13 | 11 | 9.4 | 7 | 11 | 0.67 | | | |
| 761 | hsa-miR-2278 | MIMAT0011778 | | 9 | 9 | 9 | 10 | 6 | 9 | 13 | 4 | 19 | 6 | 9.4 | 9 | 10 | 0.97 | | | |
| 762 | hsa-miR-142-5p | MIMAT0000433 | | 6 | 5 | 4 | 9 | 9 | 14 | 10 | 10 | 15 | 9 | 9.1 | 6 | 11 | 0.54 | | | |
| 763 | hsa-miR-182-5p | MIMAT0000259 | | 10 | 7 | 14 | 6 | 10 | 9 | 4 | 8 | 17 | 6 | 9.1 | 9 | 9 | 1.03 | | | |
| 764 | hsa-miR-1303 | MIMAT0005891 | | 10 | 15 | 7 | 7 | 14 | 7 | 4 | 4 | 13 | 9 | 9 | 10 | 9 | 1.15 | | | |
| 765 | hsa-miR-3140-3p | MIMAT0015008 | | 8 | 11 | 8 | 12 | 12 | 11 | 8 | 2 | 11 | 7 | 9 | 10 | 9 | 1.15 | | | |
| 766 | hsa-miR-152-3p | MIMAT0000438 | | 9 | 8 | 12 | 6 | 6 | 14 | 5 | 8 | 7 | 14 | 8.9 | 9 | 9 | 0.97 | | | |
| 767 | hsa-miR-5010-5p | MIMAT0021043 | | 10 | 11 | 7 | 9 | 13 | 6 | 10 | 8 | 8 | 4 | 8.6 | 9 | 8 | 1.13 | | | |
| 768 | hsa-miR-554 | MIMAT0003217 | | 7 | 14 | 10 | 8 | 10 | 6 | 9 | 5 | 9 | 8 | 8.6 | 10 | 8 | 1.24 | | | |
| 769 | hsa-miR-202-3p | MIMAT0002811 | | 6 | 9 | 10 | 3 | 10 | 4 | 13 | 10 | 11 | 9 | 8.5 | 7 | 10 | 0.74 | | | |
| 770 | hsa-miR-599 | MIMAT0003267 | | 2 | 7 | 9 | 10 | 7 | 15 | 9 | 5 | 7 | 13 | 8.4 | 7 | 9 | 0.75 | | | |
| 771 | hsa-miR-3605-3p | MIMAT0017982 | | 5 | 11 | 7 | 11 | 10 | 11 | 9 | 6 | 11 | 2 | 8.3 | 9 | 8 | 1.04 | | | |
| 772 | hsa-miR-9-5p | MIMAT0000441 | | 7 | 7 | 10 | 5 | 13 | 8 | 10 | 5 | 14 | 4 | 8.3 | 7 | 9 | 0.81 | | | |
| 773 | hsa-miR-5001-3p | MIMAT0021022 | | 9 | 6 | 8 | 3 | 14 | 8 | 10 | 3 | 12 | 9 | 8.2 | 7 | 9 | 0.70 | | | |
| 774 | hsa-miR-548o-3p | MIMAT0005919 | | 5 | 5 | 12 | 8 | 6 | 10 | 10 | 4 | 11 | 11 | 8.2 | 8 | 9 | 0.87 | | hsa-miR-548ah-3p | hsa-miR-548av-3p |
| 775 | hsa-miR-615-5p | MIMAT0004804 | | 6 | 11 | 5 | 7 | 13 | 9 | 7 | 3 | 10 | 11 | 8.2 | 7 | 9 | 0.82 | | | |
| 776 | hsa-miR-1247-5p | MIMAT0005899 | | 8 | 7 | 7 | 6 | 13 | 12 | 6 | 4 | 11 | 7 | 8.1 | 7 | 9 | 0.79 | | | |
| 777 | hsa-miR-1910-3p | MIMAT0026917 | | 4 | 7 | 11 | 6 | 12 | 12 | 7 | 7 | 10 | 4 | 8 | 7 | 9 | 0.81 | | | |
| 778 | hsa-miR-152-5p | MIMAT0026479 | | 10 | 7 | 14 | 6 | 6 | 3 | 9 | 5 | 8 | 11 | 7.9 | 9 | 7 | 1.32 | | | |
| 779 | hsa-miR-139-5p | MIMAT0000250 | | 7 | 12 | 6 | 7 | 10 | 12 | 5 | 3 | 6 | 8 | 7.6 | 8 | 7 | 1.09 | | | |
| 780 | hsa-miR-3179 | MIMAT0015056 | | 1 | 11 | 12 | 9 | 4 | 8 | 8 | 1 | 12 | 8 | 7.4 | 8 | 7 | 1.21 | | | |
| 781 | hsa-miR-766-5p | MIMAT0022714 | | 12 | 10 | 6 | 8 | 8 | 5 | 11 | 3 | 6 | 4 | 7.3 | 9 | 6 | 1.46 | | | |
| 782 | hsa-miR-6511a-3p | MIMAT0025479 | | 7 | 5 | 4 | 10 | 8 | 7 | 10 | 2 | 8 | 7 | 6.8 | 7 | 7 | 0.93 | | | |
| 783 | hsa-miR-924 | MIMAT0004974 | | 1 | 8 | 5 | 5 | 5 | 3 | 6 | 4 | 3 | 4 | 4.4 | 5 | 4 | 1.14 | | | |
| 784 | hsa-miR-3144-3p | MIMAT0015015 | | 7 | 1 | 1 | 1 | 1 | 1 | 1 | 3 | 6 | 10 | 3.2 | 3 | 4 | 0.68 | | | |
| 785 | hsa-miR-603 | MIMAT0003271 | | 1 | 1 | 1 | 1 | 1 | 1 | 1 | 1 | 1 | 15 | 2.4 | 1 | 3 | 0.30 | | | |
| 786 | hsa-miR-302d-3p | MIMAT0000718 | | 1 | 1 | 1 | 1 | 1 | 1 | 1 | 4 | 1 | 11 | 2.3 | 1 | 3 | 0.32 | | | |
| 787 | hsa-miR-525-3p | MIMAT0002839 | | 1 | 10 | 1 | 1 | 1 | 1 | 1 | 1 | 1 | 1 | 1.9 | 3 | 1 | 3.25 | | | |
| 788 | hsa-miR-122-5p | MIMAT0000421 | | 1 | 1 | 3 | 1 | 1 | 1 | 1 | 1 | 1 | 2 | 1.3 | 2 | 1 | 1.29 | | | |
| 789 | hsa-miR-128-3p | MIMAT0000424 | | 1 | 1 | 1 | 1 | 1 | 1 | 1 | 1 | 1 | 1 | 1 | 1 | 1 | 1.00 | | | |
| 790 | hsa-miR-1283 | MIMAT0005799 | | 1 | 1 | 1 | 1 | 1 | 1 | 1 | 1 | 1 | 1 | 1 | 1 | 1 | 1.00 | | | |
| 791 | hsa-miR-143-3p | MIMAT0000435 | | 1 | 1 | 1 | 1 | 1 | 1 | 1 | 1 | 1 | 1 | 1 | 1 | 1 | 1.00 | | | |
| 792 | hsa-miR-3065-5p | MIMAT0015066 | | 1 | 1 | 1 | 1 | 1 | 1 | 1 | 1 | 1 | 1 | 1 | 1 | 1 | 1.00 | | | |
| 793 | hsa-miR-378e | MIMAT0018927 | | 1 | 1 | 1 | 1 | 1 | 1 | 1 | 1 | 1 | 1 | 1 | 1 | 1 | 1.00 | | | |
| 794 | hsa-miR-422a | MIMAT0001339 | | 1 | 1 | 1 | 1 | 1 | 1 | 1 | 1 | 1 | 1 | 1 | 1 | 1 | 1.00 | | | |
| 795 | hsa-miR-494-3p | MIMAT0002816 | | 1 | 1 | 1 | 1 | 1 | 1 | 1 | 1 | 1 | 1 | 1 | 1 | 1 | 1.00 | | | |
| 796 | hsa-miR-502-5p | MIMAT0002873 | | 1 | 1 | 1 | 1 | 1 | 1 | 1 | 1 | 1 | 1 | 1 | 1 | 1 | 1.00 | | | |
| 797 | hsa-miR-548ar-5p | MIMAT0022265 | | 1 | 1 | 1 | 1 | 1 | 1 | 1 | 1 | 1 | 1 | 1 | 1 | 1 | 1.00 | | | |
| 798 | hsa-miR-579-3p | MIMAT0003244 | | 1 | 1 | 1 | 1 | 1 | 1 | 1 | 1 | 1 | 1 | 1 | 1 | 1 | 1.00 | | | |
| | ACTB | NM_001101.2 | | 29 | 27 | 26 | 17 | 37 | 23 | 43 | 10 | 52 | 20 | | | | | | | |
| | B2M | NM_004048.2 | | 21 | 14 | 26 | 11 | 26 | 25 | 23 | 13 | 29 | 17 | | | | | | | |
| | GAPDH | NM_002046.3 | | 20 | 20 | 26 | 24 | 24 | 22 | 22 | 9 | 20 | 19 | | | | | | | |
| | RPL19 | NM_000981.3 | | 23 | 33 | 43 | 16 | 44 | 22 | 33 | 21 | 29 | 15 | | | | | | | |
| | RPLP0 | NM_001002.3 | | 16 | 12 | 20 | 8 | 12 | 16 | 16 | 6 | 11 | 8 | | | | | | | |
| | LIG_NEG_A | nmiR00810.1 | | 10 | 8 | 16 | 9 | 20 | 5 | 12 | 8 | 14 | 11 | | | | | | | |
| | LIG_NEG_B | nmiR00828.1 | | 17 | 16 | 20 | 17 | 21 | 18 | 22 | 17 | 22 | 13 | | | | | | | |
| | LIG_NEG_C | nmiR00803.1 | | 10 | 9 | 12 | 9 | 8 | 11 | 17 | 9 | 19 | 6 | | | | | | | |
| | LIG_POS_A | nmiR00813.1 | | 14875 | 11728 | 20803 | 15867 | 21627 | 16761 | 11243 | 8693 | 30836 | 6108 | | | | | | | |
| | LIG_POS_B | nmiR00809.1 | | 4081 | 3832 | 6255 | 4375 | 6223 | 4667 | 3321 | 2191 | 8239 | 1782 | | | | | | | |
| | LIG_POS_C | nmiR00811.1 | | 633 | 537 | 542 | 691 | 865 | 569 | 345 | 355 | 1387 | 303 | | | | | | | |
| | NEG_A | ERCC_00096.1 | | 12 | 17 | 18 | 21 | 17 | 12 | 21 | 4 | 30 | 10 | | | | | | | |
| | NEG_B | ERCC_00041.1 | | 10 | 15 | 20 | 15 | 20 | 11 | 10 | 8 | 24 | 8 | | | | | | | |
| | NEG_C | ERCC_00019.1 | | 47 | 40 | 34 | 27 | 48 | 32 | 31 | 31 | 65 | 47 | | | | | | | |
| | NEG_D | ERCC_00076.1 | | 17 | 35 | 15 | 26 | 18 | 21 | 28 | 7 | 28 | 9 | | | | | | | |
| | NEG_E | ERCC_00098.1 | | 11 | 10 | 24 | 11 | 21 | 16 | 12 | 6 | 31 | 8 | | | | | | | |
| | NEG_F | ERCC_00126.1 | | 15 | 16 | 19 | 15 | 39 | 13 | 16 | 8 | 26 | 19 | | | | | | | |
| | NEG_G | ERCC_00144.1 | | 5 | 14 | 11 | 11 | 11 | 10 | 8 | 5 | 19 | 9 | | | | | | | |
| | NEG_H | ERCC_00154.1 | | 13 | 12 | 12 | 16 | 14 | 10 | 9 | 13 | 17 | 12 | | | | | | | |
| | | | | **16.25** | **19.875** | **19.125** | **17.75** | **23.5** | **15.625** | **16.875** | **10.25** | **30** | **15.25** | **18.4** | **mean background** | | | | | |
| | POS_A | ERCC_00117.1 | | 18305 | 20802 | 33026 | 16693 | 25521 | 20414 | 26444 | 8873 | 32703 | 7586 | | | | | | | |
| | POS_B | ERCC_00112.1 | | 6228 | 6436 | 9965 | 5356 | 8063 | 6560 | 8393 | 2897 | 10499 | 2394 | | | | | | | |
| | POS_C | ERCC_00002.1 | | 2023 | 2106 | 3275 | 1800 | 2542 | 2111 | 2603 | 1012 | 3259 | 719 | | | | | | | |
| | POS_D | ERCC_00092.1 | | 379 | 411 | 687 | 353 | 542 | 449 | 552 | 234 | 661 | 187 | | | | | | | |
| | POS_E | ERCC_00035.1 | | 89 | 110 | 164 | 78 | 125 | 108 | 118 | 46 | 162 | 42 | | | | | | | |
| | POS_F | ERCC_00034.1 | | 57 | 68 | 71 | 54 | 77 | 39 | 63 | 27 | 90 | 28 | | | | | | | |
| | ath-miR159a | MIMAT0000177 | | 31 | 42 | 65 | 33 | 62 | 47 | 44 | 22 | 59 | 22 | | | | | | | |
| | cel-miR-248 | MIMAT0000304 | | 22 | 27 | 24 | 17 | 28 | 20 | 28 | 12 | 33 | 19 | | | | | | | |
| | cel-miR-254 | MIMAT0000310 | | 20 | 17 | 28 | 25 | 23 | 23 | 19 | 9 | 38 | 16 | | | | | | | |
| | osa-miR414 | MIMAT0001330 | | 22 | 21 | 38 | 19 | 23 | 24 | 35 | 17 | 21 | 26 | | | | | | | |
| | osa-miR442 | MIMAT0001605 | | 22 | 22 | 29 | 15 | 26 | 24 | 22 | 13 | 25 | 24 | | | | | | | |

**Table S3 TAM 2.0 analysis**

| | Query set |
|---|---|
| 1 | hsa-miR-130a-3p |
| 2 | hsa-miR-25-3p |
| 3 | hsa-miR-221-3p |
| 4 | hsa-miR-30d-5p |
| 5 | hsa-miR-205-5p |
| 6 | hsa-miR-23a-3p |
| 7 | hsa-miR-320e |
| 8 | hsa-miR-4516 |
| 9 | hsa-miR-22-3p |
| 10 | hsa-let-7b-5p |
| 11 | hsa-miR-630 |
| 12 | hsa-miR-16-5p |
| 13 | hsa-miR-21-5p |
| 14 | hsa-miR-20a-5p |
| 15 | hsa-miR-20b-5p |
| 16 | hsa-miR-24-3p |
| 17 | hsa-miR-29b-3p |
| 18 | hsa-miR-222-3p |
| 19 | hsa-let-7i-5p |
| 20 | hsa-miR-365a-3p |
| 21 | hsa-miR-365b-3p |
| 22 | hsa-miR-29a-3p |
| 23 | hsa-miR-106a-5p |
| 24 | hsa-miR-365b-3p |

### Summary

The user has provided 20 miRNAs in the query list, and 23 of these miRNAs are in at least one of the following miRNA set.The default background is used.

### Result

#### Enrichment analysis results

Text file of results Results Visualization

| Term | Count | Percent | Fold | P-value | Bonferroni | FDR |
|---|---|---|---|---|---|---|
| **Category: Cluster (15 Items)** | | | | | | |
| hsa-mir-222 cluster [details] | 2 | 1 | 46.57692 | 4.44e-4 | 0.2271 | 3.68e-3 |
| hsa-mir-29b-1 cluster [details] | 2 | 1 | 46.57692 | 4.44e-4 | 0.2271 | 3.68e-3 |
| hsa-mir-181d cluster [details] | 2 | 0.4 | 18.63077 | 4.26e-3 | 1 | 0.0241 |
| hsa-mir-106a cluster [details] | 2 | 0.33333 | 15.52564 | 6.31e-3 | 1 | 0.0328 |
| hsa-mir-15a cluster [details] | 1 | 0.5 | 23.28846 | 0.0425 | 1 | 0.1655 |
| hsa-mir-15b cluster [details] | 1 | 0.5 | 23.28846 | 0.0425 | 1 | 0.1655 |
| hsa-mir-193b cluster [details] | 1 | 0.5 | 23.28846 | 0.0425 | 1 | 0.1655 |
| hsa-mir-29b-2 cluster [details] | 1 | 0.5 | 23.28846 | 0.0425 | 1 | 0.1655 |
| hsa-mir-30d cluster [details] | 1 | 0.5 | 23.28846 | 0.0425 | 1 | 0.1655 |
| hsa-mir-106b cluster [details] | 1 | 0.33333 | 15.52564 | 0.0631 | 1 | 0.2177 |
| hsa-mir-3180-5 cluster [details] | 1 | 0.25 | 11.64423 | 0.0833 | 1 | 0.2689 |
| hsa-mir-3619 cluster [details] | 1 | 0.25 | 11.64423 | 0.0833 | 1 | 0.2689 |
| hsa-mir-4724 cluster [details] | 1 | 0.25 | 11.64423 | 0.0833 | 1 | 0.2689 |
| hsa-mir-6081 cluster [details] | 1 | 0.2 | 9.31538 | 0.103 | 1 | 0.3087 |
| hsa-mir-17 cluster [details] | 1 | 0.16667 | 7.76282 | 0.1223 | 1 | 0.3519 |
| **Category: Disease (345 Items)** | | | | | | |
| Carcinoma, Colon [details] | 25 | 0.07937 | 3.69658 | 2.30e-14 | 1.18e-11 | 6.96e-12 |
| Carcinoma, Breast [details] | 24 | 0.06897 | 3.2122 | 2.76e-12 | 1.41e-9 | 4.18e-10 |
| Carcinoma, Hepatocellular [details] | 24 | 0.0708 | 3.29748 | 2.76e-12 | 1.41e-9 | 4.18e-10 |
| Carcinoma, Gastric [details] | 23 | 0.08364 | 3.89552 | 9.81e-13 | 5.02e-10 | 2.37e-10 |
| Neoplasms [unspecific] [details] | 21 | 0.10096 | 4.70248 | 1.05e-12 | 5.36e-10 | 2.11e-10 |
| Carcinoma, Lung. Non-Small-Cell [details] | 21 | 0.09589 | 4.46628 | 3.10e-12 | 1.59e-9 | 3.75e-10 |
| Carcinoma, Prostate [details] | 20 | 0.07782 | 3.62466 | 1.23e-9 | 6.28e-7 | 7.82e-8 |
| Heart Failure [details] | 19 | 0.15833 | 7.37468 | 7.86e-15 | 4.02e-12 | 3.17e-12 |
| Carcinoma, Pancreatic [details] | 19 | 0.11446 | 5.33109 | 4.34e-12 | 2.22e-9 | 4.78e-10 |
| Carcinoma, Ovarian [details] | 18 | 0.09278 | 4.32157 | 1.14e-9 | 5.82e-7 | 7.65e-8 |
| Colon Neoplasms [details] | 17 | 0.22078 | 10.28322 | 1.52e-15 | 7.78e-13 | 9.20e-13 |
| Glioma [details] | 17 | 0.08173 | 3.80677 | 4.03e-8 | 2.06e-5 | 1.16e-6 |
| Carcinoma, Bladder [details] | 16 | 0.13333 | 6.21026 | 7.56e-11 | 3.87e-8 | 7.05e-9 |
| Glioblastoma [details] | 16 | 0.09467 | 4.40965 | 1.61e-8 | 8.26e-6 | 6.10e-7 |
| Osteosarcoma [details] | 16 | 0.09357 | 4.35807 | 1.93e-8 | 9.88e-6 | 6.49e-7 |
| Atherosclerosis [details] | 15 | 0.15789 | 7.35425 | 3.37e-11 | 1.73e-8 | 3.40e-9 |
| Melanoma [details] | 15 | 0.11364 | 5.29283 | 4.66e-9 | 2.38e-6 | 2.25e-7 |
| Hepatitis C Virus Infection [details] | 14 | 0.16092 | 7.49514 | 1.66e-10 | 8.51e-8 | 1.44e-8 |
| Lung Neoplasms [details] | 14 | 0.12727 | 5.92797 | 4.48e-9 | 2.30e-6 | 2.26e-7 |
| Diabetes Mellitus, Type 2 [details] | 14 | 0.09524 | 4.4359 | 2.22e-7 | 1.14e-4 | 5.50e-6 |
| Carcinoma, Lung [details] | 14 | 0.06573 | 3.06139 | 2.42e-5 | 0.0124 | 3.25e-4 |
| Carcinoma, Nasopharyngeal [details] | 13 | 0.11404 | 5.3114 | 8.59e-8 | 4.40e-5 | 2.21e-6 |
| Leukemia, Myeloid, Acute [details] | 13 | 0.10924 | 5.08824 | 1.47e-7 | 7.51e-5 | 3.70e-6 |
| Pancreatic Neoplasms [details] | 12 | 0.1875 | 8.73317 | 9.09e-10 | 4.65e-7 | 6.47e-8 |
| Preeclampsia [details] | 12 | 0.16216 | 7.55301 | 5.42e-9 | 2.78e-6 | 2.43e-7 |
| Prostate Neoplasms [details] | 12 | 0.16 | 7.45231 | 6.38e-9 | 3.27e-6 | 2.76e-7 |
| Ovarian Neoplasms [details] | 12 | 0.11321 | 5.27286 | 3.77e-7 | 1.93e-4 | 8.45e-6 |
| Carcinoma, Renal Cell [details] | 12 | 0.10084 | 4.69683 | 1.40e-6 | 7.15e-4 | 2.77e-5 |
| Glaucoma [details] | 11 | 0.37931 | 17.66711 | 1.10e-12 | 5.65e-10 | 1.91e-10 |
| Multiple Myeloma [details] | 11 | 0.18333 | 8.5391 | 7.58e-9 | 3.88e-6 | 3.17e-7 |
| Carcinoma, Cervical [details] | 11 | 0.07971 | 3.71265 | 4.89e-5 | 0.0251 | 5.87e-4 |
| Myocardial Infarction [details] | 10 | 0.19231 | 8.9571 | 2.78e-8 | 1.42e-5 | 8.62e-7 |

| | | | | | | |
|---|---|---|---|---|---|---|
| Rheumatoid Arthritis [details] | 10 | 0.18519 | 8.62536 | 4.10e-8 | 2.10e-5 | 1.15e-6 |
| Hepatitis B Virus Infection [details] | 10 | 0.14925 | 6.95178 | 3.60e-7 | 1.85e-4 | 8.39e-6 |
| Leukemia [details] | 10 | 0.14925 | 6.95178 | 3.60e-7 | 1.85e-4 | 8.39e-6 |
| Adenocarcinoma, Lung [details] | 10 | 0.09901 | 4.61158 | 1.81e-5 | 9.25e-3 | 2.61e-4 |
| Squamous Cell Carcinoma, Esophageal [details] | 10 | 0.07042 | 3.28007 | 3.65e-4 | 0.1868 | 3.09e-3 |
| Breast Neoplasms [details] | 10 | 0.06757 | 3.14709 | 5.16e-4 | 0.2643 | 3.79e-3 |
| Pleural Mesothelioma [details] | 9 | 0.28125 | 13.09976 | 4.35e-9 | 2.23e-6 | 2.29e-7 |
| Leukemia, Lymphocytic, Chronic, B-Cell [details] | 9 | 0.26471 | 12.32919 | 7.92e-9 | 4.06e-6 | 3.20e-7 |
| Cholangiocarcinoma [details] | 9 | 0.25714 | 11.97692 | 1.05e-8 | 5.39e-6 | 4.11e-7 |
| Stroke [details] | 9 | 0.24324 | 11.32952 | 1.81e-8 | 9.25e-6 | 6.63e-7 |
| Adenocarcinoma, Gastric [details] | 9 | 0.23684 | 11.03138 | 2.34e-8 | 1.20e-5 | 7.45e-7 |
| Parkinson\'s Disease [details] | 9 | 0.21951 | 10.2242 | 4.83e-8 | 2.47e-5 | 1.33e-6 |
| Choriocarcinoma [details] | 9 | 0.15 | 6.98654 | 1.59e-6 | 8.13e-4 | 3.10e-5 |
| Lymphoma [details] | 9 | 0.15 | 6.98654 | 1.59e-6 | 8.13e-4 | 3.10e-5 |
| Cardiovascular Diseases [unspecific] [details] | 9 | 0.14063 | 6.54988 | 2.80e-6 | 1.44e-3 | 5.07e-5 |
| Gastrointestinal Neoplasms [details] | 9 | 0.10976 | 5.1121 | 2.35e-5 | 0.012 | 3.19e-4 |
| Squamous Cell Carcinoma, Oral [details] | 9 | 0.10843 | 5.05051 | 2.60e-5 | 0.0133 | 3.38e-4 |
| Malignant Neoplasms [unspecific] [details] | 8 | 0.16 | 7.45231 | 4.24e-6 | 2.17e-3 | 7.22e-5 |
| Cardiomyopathy, Hypertrophic [details] | 8 | 0.14545 | 6.77483 | 8.97e-6 | 4.59e-3 | 1.43e-4 |
| Carcinoma, Laryngeal [details] | 8 | 0.12698 | 5.91453 | 2.55e-5 | 0.0131 | 3.40e-4 |
| Leukemia, Myeloid, Chronic [details] | 8 | 0.1194 | 5.56142 | 4.07e-5 | 0.0208 | 4.93e-4 |
| Diabetes Mellitus [details] | 8 | 0.10811 | 5.03534 | 8.54e-5 | 0.0437 | 9.23e-4 |
| Spinal Cord Injuries [details] | 7 | 0.53846 | 25.07988 | 1.39e-9 | 7.14e-7 | 8.44e-8 |
| Leukemia, Lymphoblastic [details] | 7 | 0.33333 | 15.52564 | 8.45e-8 | 4.33e-5 | 2.22e-6 |
| Influenza [details] | 7 | 0.26923 | 12.53994 | 4.46e-7 | 2.28e-4 | 9.81e-6 |
| Squamous Cell Carcinoma, Tongue [details] | 7 | 0.25926 | 12.0755 | 5.93e-7 | 3.04e-4 | 1.24e-5 |
| Pulmonary Hypertension [details] | 7 | 0.19444 | 9.05662 | 4.91e-6 | 2.51e-3 | 8.26e-5 |
| Leukemia, Lymphocytic, Chronic [details] | 7 | 0.17073 | 7.95216 | 1.23e-5 | 6.31e-3 | 1.89e-4 |
| Osteoarthritis [details] | 7 | 0.17073 | 7.95216 | 1.23e-5 | 6.31e-3 | 1.89e-4 |
| Liver Cirrhosis [details] | 7 | 0.12281 | 5.71997 | 1.15e-4 | 0.0591 | 1.18e-3 |
| Neuroblastoma [details] | 7 | 0.11864 | 5.52608 | 1.45e-4 | 0.0742 | 1.44e-3 |
| Atrial Fibrillation [details] | 7 | 0.11667 | 5.43397 | 1.62e-4 | 0.0828 | 1.57e-3 |
| Pulmonary Sarcoidosis [details] | 6 | 0.54545 | 25.40559 | 2.29e-8 | 1.17e-5 | 7.49e-7 |
| Diabetic Retinopathy [details] | 6 | 0.28571 | 13.30769 | 2.33e-6 | 1.19e-3 | 4.34e-5 |
| Lymphoma, Hodgkin [details] | 6 | 0.27273 | 12.7028 | 3.15e-6 | 1.62e-3 | 5.62e-5 |
| Adenocarcinoma, Colon [details] | 6 | 0.26087 | 12.1505 | 4.21e-6 | 2.15e-3 | 7.38e-5 |
| Squamous Cell Carcinoma, Lung [details] | 6 | 0.26087 | 12.1505 | 4.21e-6 | 2.15e-3 | 7.38e-5 |
| Leukemia, Lymphoblastic, Acute [details] | 6 | 0.25 | 11.64423 | 5.53e-6 | 2.83e-3 | 9.05e-5 |
| Vascular Diseases [unspecific] [details] | 6 | 0.22222 | 10.35043 | 1.16e-5 | 5.96e-3 | 1.81e-4 |
| Muscular Dystrophy [details] | 6 | 0.2069 | 9.6366 | 1.82e-5 | 9.29e-3 | 2.59e-4 |
| Autoimmune Diseases [unspecific] [details] | 6 | 0.16667 | 7.76282 | 6.73e-5 | 0.0344 | 7.61e-4 |
| Chronic Hepatitis C [details] | 6 | 0.16667 | 7.76282 | 6.73e-5 | 0.0344 | 7.61e-4 |
| Endometriosis [details] | 6 | 0.14634 | 6.81614 | 1.44e-4 | 0.0739 | 1.45e-3 |
| Human Immunodeficiency Virus Infection [details] | 6 | 0.12766 | 5.94599 | 3.16e-4 | 0.1618 | 2.75e-3 |
| Squamous Cell Carcinoma, Head and Neck [details] | 6 | 0.125 | 5.82212 | 3.56e-4 | 0.1822 | 3.04e-3 |
| Schizophrenia [details] | 6 | 0.11321 | 5.27286 | 6.19e-4 | 0.3169 | 4.44e-3 |
| Carcinoma, Breast, Triple Negative [details] | 6 | 0.1 | 4.65769 | 1.22e-3 | 0.6239 | 8.02e-3 |
| Carcinoma, Esophageal [details] | 6 | 0.09091 | 4.23427 | 2.03e-3 | 1 | 0.0123 |
| Alzheimer\'s Disease [details] | 6 | 0.06 | 2.79462 | 0.0161 | 1 | 0.073 |
| Acute Cerebral Infarction [details] | 5 | 0.83333 | 38.8141 | 1.81e-8 | 9.25e-6 | 6.44e-7 |
| Carcinoma, Adenoid Cystic [details] | 5 | 0.71429 | 33.26923 | 6.23e-8 | 3.19e-5 | 1.68e-6 |
| Ewing\'s Sarcoma [details] | 5 | 0.41667 | 19.40705 | 2.18e-6 | 1.12e-3 | 4.13e-5 |
| Bladder Outlet Obstruction [details] | 5 | 0.35714 | 16.63462 | 5.36e-6 | 2.75e-3 | 8.90e-5 |
| Thyroid Neoplasms [details] | 5 | 0.3125 | 14.55529 | 1.14e-5 | 5.82e-3 | 1.79e-4 |
| Astrocytoma [details] | 5 | 0.29412 | 13.6991 | 1.59e-5 | 8.12e-3 | 2.34e-4 |
| Respiratory Syncytial Virus Pneumonia [details] | 5 | 0.27778 | 12.93803 | 2.16e-5 | 0.0111 | 3.01e-4 |
| Squamous Cell Carcinoma, Skin or Unspecific [details] | 5 | 0.27778 | 12.93803 | 2.16e-5 | 0.0111 | 3.01e-4 |
| Polycythemia Vera [details] | 5 | 0.25 | 11.64423 | 3.80e-5 | 0.0195 | 4.70e-4 |

| | | | | | | |
|---|---|---|---|---|---|---|
| Carcinoma, Lung, Small-Cell [details] | 5 | 0.2381 | 11.08974 | 4.92e-5 | 0.0252 | 5.84e-4 |
| Colon Adenoma [details] | 5 | 0.2381 | 11.08974 | 4.92e-5 | 0.0252 | 5.84e-4 |
| Cardiomyopathy [details] | 5 | 0.20833 | 9.70353 | 9.83e-5 | 0.0503 | 1.02e-3 |
| Lymphoma, T-Cell [details] | 5 | 0.19231 | 8.9571 | 1.48e-4 | 0.0757 | 1.44e-3 |
| Kaposi\'s Sarcoma [details] | 5 | 0.18519 | 8.62536 | 1.79e-4 | 0.0915 | 1.68e-3 |
| Chronic Obstructive Pulmonary Disease [details] | 5 | 0.17857 | 8.31731 | 2.14e-4 | 0.1098 | 1.98e-3 |
| Kidney Injury [details] | 5 | 0.17857 | 8.31731 | 2.14e-4 | 0.1098 | 1.98e-3 |
| Myelodysplastic Syndromes [details] | 5 | 0.17857 | 8.31731 | 2.14e-4 | 0.1098 | 1.98e-3 |
| Carcinoma, Adrenocortical [details] | 5 | 0.16667 | 7.76282 | 3.02e-4 | 0.1545 | 2.65e-3 |
| Chronic Kidney Disease [details] | 5 | 0.15625 | 7.27764 | 4.14e-4 | 0.212 | 3.48e-3 |
| Crohn Disease [details] | 5 | 0.15152 | 7.05711 | 4.81e-4 | 0.2462 | 3.55e-3 |
| Human Papilloma Virus Infection [details] | 5 | 0.14706 | 6.84955 | 5.56e-4 | 0.2845 | 4.01e-3 |
| Carcinoma, Thyroid [details] | 5 | 0.13889 | 6.46902 | 7.31e-4 | 0.3742 | 5.06e-3 |
| Head And Neck Neoplasms [details] | 5 | 0.13889 | 6.46902 | 7.31e-4 | 0.3742 | 5.06e-3 |
| Retinoblastoma [details] | 5 | 0.11111 | 5.17521 | 2.07e-3 | 1 | 0.0125 |
| Carcinoma, Thyroid, Papillary [details] | 5 | 0.08929 | 4.15865 | 5.51e-3 | 1 | 0.0293 |
| Coronary Heart Diseases [details] | 5 | 0.08929 | 4.15865 | 5.51e-3 | 1 | 0.0293 |
| Carcinoma, Endometrial [details] | 5 | 0.08621 | 4.01525 | 6.42e-3 | 1 | 0.0325 |
| Adenocarcinoma, Pancreatic Ductal [details] | 5 | 0.07042 | 3.28007 | 0.015 | 1 | 0.0688 |
| Ependymoma [details] | 4 | 0.66667 | 31.05128 | 2.44e-6 | 1.25e-3 | 4.48e-5 |
| Atopic Dermatitis [details] | 4 | 0.44444 | 20.70085 | 1.96e-5 | 0.0101 | 2.76e-4 |
| Cerebral Ischemia [details] | 4 | 0.36364 | 16.93706 | 4.99e-5 | 0.0256 | 5.76e-4 |
| Carcinoma, Thyroid, Anaplastic [details] | 4 | 0.33333 | 15.52564 | 7.38e-5 | 0.0378 | 8.12e-4 |
| Lymphoma, Primary Effusion [details] | 4 | 0.30769 | 14.33136 | 1.05e-4 | 0.0538 | 1.08e-3 |
| Peripheral Vascular Disease [details] | 4 | 0.28571 | 13.30769 | 1.45e-4 | 0.0742 | 1.43e-3 |
| Bronchopulmonary Dysplasia [details] | 4 | 0.25 | 11.64423 | 2.56e-4 | 0.131 | 2.31e-3 |
| Myocardial Fibrosis [details] | 4 | 0.25 | 11.64423 | 2.56e-4 | 0.131 | 2.31e-3 |
| Early-Stage Breast Carcinoma [details] | 4 | 0.23529 | 10.95928 | 3.30e-4 | 0.1688 | 2.85e-3 |
| Arteriosclerosis Obliterans [details] | 4 | 0.22222 | 10.35043 | 4.18e-4 | 0.2138 | 3.49e-3 |
| Allergy [details] | 4 | 0.19048 | 8.87179 | 7.82e-4 | 0.4002 | 5.35e-3 |
| Aortic Aneurysm, Abdominal [details] | 4 | 0.19048 | 8.87179 | 7.82e-4 | 0.4002 | 5.35e-3 |
| Ischemia-Reperfusion Injury [details] | 4 | 0.17391 | 8.10033 | 1.12e-3 | 0.5749 | 7.43e-3 |
| Carcinoma, Ovarian, Serous [details] | 4 | 0.16667 | 7.76282 | 1.33e-3 | 0.6799 | 8.33e-3 |
| Aortic Stenosis [details] | 4 | 0.16 | 7.45231 | 1.56e-3 | 0.7975 | 9.72e-3 |
| Liver Neoplasms [details] | 4 | 0.16 | 7.45231 | 1.56e-3 | 0.7975 | 9.72e-3 |
| Tuberculosis, Pulmonary [details] | 4 | 0.15385 | 7.16568 | 1.81e-3 | 0.9288 | 0.0111 |
| Epilepsy [details] | 4 | 0.14815 | 6.90028 | 2.10e-3 | 1 | 0.0126 |
| HIV-Associated Lipodystrophy [details] | 4 | 0.14286 | 6.65385 | 2.41e-3 | 1 | 0.0142 |
| Carcinoma, Urothelial, Upper Tract [details] | 4 | 0.12903 | 6.00993 | 3.55e-3 | 1 | 0.0204 |
| Hematologic Neoplasms [details] | 4 | 0.11765 | 5.47964 | 5.00e-3 | 1 | 0.0268 |
| Stroke, Ischemic [details] | 4 | 0.11765 | 5.47964 | 5.00e-3 | 1 | 0.0268 |
| Lymphoma, Large B-Cell, Diffuse [details] | 4 | 0.10526 | 4.90283 | 7.51e-3 | 1 | 0.0377 |
| Hypertension [details] | 4 | 0.10256 | 4.77712 | 8.24e-3 | 1 | 0.0408 |
| Asthma [details] | 4 | 0.09756 | 4.54409 | 9.86e-3 | 1 | 0.0472 |
| Multiple Sclerosis [details] | 4 | 0.08511 | 3.96399 | 0.0159 | 1 | 0.0724 |
| Carcinoma, Rectal [details] | 4 | 0.08333 | 3.88141 | 0.0171 | 1 | 0.0769 |
| Obesity [details] | 4 | 0.07843 | 3.65309 | 0.021 | 1 | 0.0925 |
| Systemic Lupus Erythematosus [details] | 4 | 0.07273 | 3.38741 | 0.027 | 1 | 0.1121 |
| Carcinoma, Renal Cell, Clear-Cell [details] | 4 | 0.06897 | 3.2122 | 0.0322 | 1 | 0.1312 |
| Cardiometabolic Disorders [details] | 3 | 0.75 | 34.93269 | 3.47e-5 | 0.0178 | 4.43e-4 |
| Acquired Immunodeficiency Syndrome [details] | 3 | 0.6 | 27.94615 | 8.56e-5 | 0.0438 | 9.17e-4 |
| Carcinoma, Hepatocellular, HCV-Related [details] | 3 | 0.6 | 27.94615 | 8.56e-5 | 0.0438 | 9.17e-4 |
| Central Nervous System Embryonal Tumor [details] | 3 | 0.6 | 27.94615 | 8.56e-5 | 0.0438 | 9.17e-4 |
| Cholestatic Liver Injury [details] | 3 | 0.6 | 27.94615 | 8.56e-5 | 0.0438 | 9.17e-4 |
| Chronic Heart Failure [details] | 3 | 0.5 | 23.28846 | 1.69e-4 | 0.0864 | 1.61e-3 |
| Leiomyoma [details] | 3 | 0.5 | 23.28846 | 1.69e-4 | 0.0864 | 1.61e-3 |
| Vasculitis [details] | 3 | 0.375 | 17.46635 | 4.59e-4 | 0.235 | 3.52e-3 |
| Ankylosing Spondylitis [details] | 3 | 0.33333 | 15.52564 | 6.79e-4 | 0.3474 | 4.78e-3 |

| | | | | | | |
|---|---|---|---|---|---|---|
| Carcinoma, Basal Cell [details] | 3 | 0.3 | 13.97308 | 9.56e-4 | 0.4893 | 6.43e-3 |
| Viral Myocarditis [details] | 3 | 0.3 | 13.97308 | 9.56e-4 | 0.4893 | 6.43e-3 |
| ACTH-Secreting Pituitary Adenoma [details] | 3 | 0.27273 | 12.7028 | 1.30e-3 | 0.6631 | 8.48e-3 |
| Aortic Aneurysm [details] | 3 | 0.27273 | 12.7028 | 1.30e-3 | 0.6631 | 8.48e-3 |
| Leukemia, Lymphoblastic, Acute, T-Cell [details] | 3 | 0.25 | 11.64423 | 1.70e-3 | 0.8716 | 0.0105 |
| Lofgren\'s Syndrome [details] | 3 | 0.25 | 11.64423 | 1.70e-3 | 0.8716 | 0.0105 |
| Lymphoma, Burkitt\'s [details] | 3 | 0.23077 | 10.74852 | 2.18e-3 | 1 | 0.013 |
| Tuberculosis [details] | 3 | 0.23077 | 10.74852 | 2.18e-3 | 1 | 0.013 |
| Vascular Injuries [details] | 3 | 0.23077 | 10.74852 | 2.18e-3 | 1 | 0.013 |
| Liver Injury [details] | 3 | 0.2 | 9.31538 | 3.37e-3 | 1 | 0.0195 |
| Myeloma [details] | 3 | 0.1875 | 8.73317 | 4.09e-3 | 1 | 0.0233 |
| Rhabdomyosarcoma [details] | 3 | 0.16667 | 7.76282 | 5.79e-3 | 1 | 0.0304 |
| Cutaneous Melanoma [details] | 3 | 0.15789 | 7.35425 | 6.78e-3 | 1 | 0.0342 |
| Cerebral Aneurysm [details] | 3 | 0.15 | 6.98654 | 7.87e-3 | 1 | 0.0394 |
| Vascular Hypertrophy [details] | 3 | 0.14286 | 6.65385 | 9.05e-3 | 1 | 0.0436 |
| Adenocarcinoma, Esophageal [details] | 3 | 0.13636 | 6.3514 | 0.0103 | 1 | 0.0492 |
| Lung Fibrosis [details] | 3 | 0.13636 | 6.3514 | 0.0103 | 1 | 0.0492 |
| Lymphoma, B-Cell [details] | 3 | 0.125 | 5.82212 | 0.0132 | 1 | 0.0612 |
| Hepatoblastoma [details] | 3 | 0.11538 | 5.37426 | 0.0165 | 1 | 0.0744 |
| Autism Spectrum Disorder [details] | 3 | 0.10714 | 4.99038 | 0.0202 | 1 | 0.0898 |
| Squamous Cell Carcinoma, Laryngeal or Hypopharyngeal [details] | 3 | 0.10714 | 4.99038 | 0.0202 | 1 | 0.0898 |
| Liver Diseases [unspecific] [details] | 3 | 0.10345 | 4.8183 | 0.0222 | 1 | 0.0942 |
| Polycystic Ovarian Syndrome [details] | 3 | 0.10345 | 4.8183 | 0.0222 | 1 | 0.0942 |
| Peritoneal Dialysis Failure [details] | 3 | 0.09677 | 4.50744 | 0.0265 | 1 | 0.1103 |
| Chronic Hepatitis B [details] | 3 | 0.09375 | 4.36659 | 0.0288 | 1 | 0.1192 |
| Inflammatory Bowel Diseases [details] | 3 | 0.09375 | 4.36659 | 0.0288 | 1 | 0.1192 |
| Psoriasis [details] | 3 | 0.09091 | 4.23427 | 0.0313 | 1 | 0.128 |
| Heart Diseases [unspecific] [details] | 3 | 0.07895 | 3.67713 | 0.045 | 1 | 0.1595 |
| Bladder Neoplasms [details] | 3 | 0.08108 | 3.77651 | 0.0421 | 1 | 0.1644 |
| Acute Myocardial Infarction [details] | 3 | 0.06522 | 3.03763 | 0.0724 | 1 | 0.2356 |
| Adenocarcinoma, Ampullary [details] | 2 | 1 | 46.57692 | 4.44e-4 | 0.2271 | 3.68e-3 |
| Chronic Inflammatory Pain [details] | 2 | 1 | 46.57692 | 4.44e-4 | 0.2271 | 3.68e-3 |
| Multiple Sporadic Gastrointestinal Stromal Tumor [details] | 2 | 1 | 46.57692 | 4.44e-4 | 0.2271 | 3.68e-3 |
| Sciatic Nerve Injury [details] | 2 | 1 | 46.57692 | 4.44e-4 | 0.2271 | 3.68e-3 |
| Carotid Artery Diseases [details] | 2 | 0.66667 | 31.05128 | 1.31e-3 | 0.6723 | 8.41e-3 |
| Endomyocardial Fibrosis [details] | 2 | 0.66667 | 31.05128 | 1.31e-3 | 0.6723 | 8.41e-3 |
| Erythropoiesis [details] | 2 | 0.66667 | 31.05128 | 1.31e-3 | 0.6723 | 8.41e-3 |
| Leukemia, Acute [details] | 2 | 0.66667 | 31.05128 | 1.31e-3 | 0.6723 | 8.41e-3 |
| Marek Disease [details] | 2 | 0.5 | 23.28846 | 2.59e-3 | 1 | 0.0152 |
| Plasmodium vivax Infection [details] | 2 | 0.5 | 23.28846 | 2.59e-3 | 1 | 0.0152 |
| Adenocarcinoma, Prostate [details] | 2 | 0.4 | 18.63077 | 4.26e-3 | 1 | 0.0241 |
| Hyperlipidemia [details] | 2 | 0.4 | 18.63077 | 4.26e-3 | 1 | 0.0241 |
| Pregnancy Complications [unspecific] [details] | 2 | 0.4 | 18.63077 | 4.26e-3 | 1 | 0.0241 |
| Stroke, Hemorrhagic [details] | 2 | 0.4 | 18.63077 | 4.26e-3 | 1 | 0.0241 |
| Thyroid Lymphoma [details] | 2 | 0.4 | 18.63077 | 4.26e-3 | 1 | 0.0241 |
| Uterine Leiomyoma [details] | 2 | 0.4 | 18.63077 | 4.26e-3 | 1 | 0.0241 |
| Wounds and Injuries [unspecific] [details] | 2 | 0.4 | 18.63077 | 4.26e-3 | 1 | 0.0241 |
| Irritable Bowel Syndrome [details] | 2 | 0.33333 | 15.52564 | 6.31e-3 | 1 | 0.0328 |
| Leukemia, Biphenotypic, Acute [details] | 2 | 0.33333 | 15.52564 | 6.31e-3 | 1 | 0.0328 |
| Lymphoma, Mantle-Cell [details] | 2 | 0.33333 | 15.52564 | 6.31e-3 | 1 | 0.0328 |
| Non-Traumatic Osteonecrosis [details] | 2 | 0.33333 | 15.52564 | 6.31e-3 | 1 | 0.0328 |
| Carcinoma, Biliary Tract [details] | 2 | 0.28571 | 13.30769 | 8.72e-3 | 1 | 0.0429 |
| Carcinoma, Endometrioid Endometrial [details] | 2 | 0.28571 | 13.30769 | 8.72e-3 | 1 | 0.0429 |
| Lung Injury [unspecific] [details] | 2 | 0.28571 | 13.30769 | 8.72e-3 | 1 | 0.0429 |
| Carcinoma, Skin [details] | 2 | 0.25 | 11.64423 | 0.0115 | 1 | 0.0538 |
| Metabolic Syndrome [details] | 2 | 0.22222 | 10.35043 | 0.0146 | 1 | 0.0673 |
| Systemic Sclerosis [details] | 2 | 0.22222 | 10.35043 | 0.0146 | 1 | 0.0673 |
| Barrett\'s Esophagus [details] | 2 | 0.18182 | 8.46853 | 0.0217 | 1 | 0.095 |

| | | | | | | |
|---|---|---|---|---|---|---|
| Machado-Joseph Disease [details] | 2 | 0.18182 | 8.46853 | 0.0217 | 1 | 0.095 |
| Mesothelioma [details] | 2 | 0.18182 | 8.46853 | 0.0217 | 1 | 0.095 |
| Uveal Melanoma [details] | 2 | 0.18182 | 8.46853 | 0.0217 | 1 | 0.095 |
| Adrenal Cortex Neoplasms [details] | 2 | 0.16667 | 7.76282 | 0.0256 | 1 | 0.1082 |
| Oral Leukoplakia [details] | 2 | 0.16667 | 7.76282 | 0.0256 | 1 | 0.1082 |
| Keloid [details] | 2 | 0.15385 | 7.16568 | 0.0299 | 1 | 0.1228 |
| Acute Ischemic Stroke [details] | 2 | 0.14286 | 6.65385 | 0.0344 | 1 | 0.1395 |
| Chronic Inflammation [details] | 2 | 0.14286 | 6.65385 | 0.0344 | 1 | 0.1395 |
| Coronary Atherosclerosis [details] | 2 | 0.14286 | 6.65385 | 0.0344 | 1 | 0.1395 |
| Leukemia, B-Cell [details] | 2 | 0.14286 | 6.65385 | 0.0344 | 1 | 0.1395 |
| Macular Degeneration [details] | 2 | 0.14286 | 6.65385 | 0.0344 | 1 | 0.1395 |
| Myasthenia Gravis [details] | 2 | 0.14286 | 6.65385 | 0.0344 | 1 | 0.1395 |
| Adenocarcinoma, Gastric-Esophageal Junction [details] | 2 | 0.13333 | 6.21026 | 0.0392 | 1 | 0.1557 |
| Carcinoma, Urothelial [details] | 2 | 0.13333 | 6.21026 | 0.0392 | 1 | 0.1557 |
| Idiopathic Pulmonary Fibrosis [details] | 2 | 0.13333 | 6.21026 | 0.0392 | 1 | 0.1557 |
| Maternal Obesity During Childbirth [details] | 2 | 0.13333 | 6.21026 | 0.0392 | 1 | 0.1557 |
| Cystic Fibrosis [details] | 2 | 0.125 | 5.82212 | 0.0442 | 1 | 0.1575 |
| Pituitary Neoplasms [details] | 2 | 0.125 | 5.82212 | 0.0442 | 1 | 0.1575 |
| Muscle Atrophy [details] | 2 | 0.11765 | 5.47964 | 0.0495 | 1 | 0.1747 |
| Squamous Cell Carcinoma, Cerevial [details] | 2 | 0.11765 | 5.47964 | 0.0495 | 1 | 0.1747 |
| Epstein-Barr Virus Infection [details] | 2 | 0.11111 | 5.17521 | 0.0549 | 1 | 0.1923 |
| Leukemia, Myeloid [details] | 2 | 0.11111 | 5.17521 | 0.0549 | 1 | 0.1923 |
| Cardiomyopathy, Ischemic [details] | 2 | 0.10526 | 4.90283 | 0.0606 | 1 | 0.2103 |
| Cervical Neoplasms [details] | 2 | 0.10526 | 4.90283 | 0.0606 | 1 | 0.2103 |
| Neuropsychiatric Disorders [unspecific] [details] | 2 | 0.09091 | 4.23427 | 0.0787 | 1 | 0.2556 |
| Diabetes Mellitus, Gestational [details] | 2 | 0.08696 | 4.05017 | 0.0851 | 1 | 0.2564 |
| Brain Neoplasms [details] | 2 | 0.08333 | 3.88141 | 0.0916 | 1 | 0.2754 |
| Colitis, Ulcerative [details] | 2 | 0.07692 | 3.58284 | 0.1052 | 1 | 0.3039 |
| Carcinoma, Gallbladder [details] | 2 | 0.07407 | 3.45014 | 0.1121 | 1 | 0.3232 |
| Sepsis [details] | 2 | 0.05882 | 2.73982 | 0.1637 | 1 | 0.4348 |
| Carcinoma, Oral [details] | 2 | 0.05405 | 2.51767 | 0.187 | 1 | 0.4861 |
| Atypical Teratoid Tumor [details] | 1 | 0.5 | 23.28846 | 0.0425 | 1 | 0.1655 |
| Choroidal Neovascularization [details] | 1 | 0.5 | 23.28846 | 0.0425 | 1 | 0.1655 |
| Chronic Alcohol-Induced Alveolar Macrophage Dysfunction [details] | 1 | 0.5 | 23.28846 | 0.0425 | 1 | 0.1655 |
| Emphysema [details] | 1 | 0.5 | 23.28846 | 0.0425 | 1 | 0.1655 |
| Fetal Alcohol Syndrome [details] | 1 | 0.5 | 23.28846 | 0.0425 | 1 | 0.1655 |
| Hamartoma Syndrome [details] | 1 | 0.5 | 23.28846 | 0.0425 | 1 | 0.1655 |
| In Vitro Fertilization Failure [details] | 1 | 0.5 | 23.28846 | 0.0425 | 1 | 0.1655 |
| Invasive Candida Infection [details] | 1 | 0.5 | 23.28846 | 0.0425 | 1 | 0.1655 |
| Narcolepsy [details] | 1 | 0.5 | 23.28846 | 0.0425 | 1 | 0.1655 |
| Neutropenia [details] | 1 | 0.5 | 23.28846 | 0.0425 | 1 | 0.1655 |
| Periodontitis [details] | 1 | 0.5 | 23.28846 | 0.0425 | 1 | 0.1655 |
| Silicosis [details] | 1 | 0.5 | 23.28846 | 0.0425 | 1 | 0.1655 |
| Thyroid-Associated Ophthalmopathy [details] | 1 | 0.5 | 23.28846 | 0.0425 | 1 | 0.1655 |
| Type 2 Diabetes Complications [details] | 1 | 0.5 | 23.28846 | 0.0425 | 1 | 0.1655 |
| Bronchiolitis Obliterans Syndrome [details] | 1 | 0.33333 | 15.52564 | 0.0631 | 1 | 0.2177 |
| Carcinoma, Non-Functioning Pituitary [details] | 1 | 0.33333 | 15.52564 | 0.0631 | 1 | 0.2177 |
| Chronic Atrial Fibrillation [details] | 1 | 0.33333 | 15.52564 | 0.0631 | 1 | 0.2177 |
| Granular Corneal Dystrophy [details] | 1 | 0.33333 | 15.52564 | 0.0631 | 1 | 0.2177 |
| Growth Disorders [details] | 1 | 0.33333 | 15.52564 | 0.0631 | 1 | 0.2177 |
| Hereditary Breast Carcinoma [details] | 1 | 0.33333 | 15.52564 | 0.0631 | 1 | 0.2177 |
| Liver Failure [details] | 1 | 0.33333 | 15.52564 | 0.0631 | 1 | 0.2177 |
| Malignant Peripheral Nerve Sheath Tumor [details] | 1 | 0.33333 | 15.52564 | 0.0631 | 1 | 0.2177 |
| Primary Biliary Cirrhosis [details] | 1 | 0.33333 | 15.52564 | 0.0631 | 1 | 0.2177 |
| Recurrent Wheezing [details] | 1 | 0.33333 | 15.52564 | 0.0631 | 1 | 0.2177 |
| Scleroderma, Systemic [details] | 1 | 0.33333 | 15.52564 | 0.0631 | 1 | 0.2177 |
| Carcinoma, Oropharyngeal [details] | 1 | 0.25 | 11.64423 | 0.0833 | 1 | 0.2689 |
| Carcinoma, Ovarian, Clear Cell [details] | 1 | 0.25 | 11.64423 | 0.0833 | 1 | 0.2689 |

| | | | | | | |
|---|---|---|---|---|---|---|
| Carcinoma, Salivary Adenoid Cystic [details] | 1 | 0.25 | 11.64423 | 0.0833 | 1 | 0.2689 |
| Carcinoma, Spindle Cell [details] | 1 | 0.25 | 11.64423 | 0.0833 | 1 | 0.2689 |
| Cholesteatoma [details] | 1 | 0.25 | 11.64423 | 0.0833 | 1 | 0.2689 |
| Chronic Pancreatitis [details] | 1 | 0.25 | 11.64423 | 0.0833 | 1 | 0.2689 |
| Cryptosporidium infection [details] | 1 | 0.25 | 11.64423 | 0.0833 | 1 | 0.2689 |
| Diabetic Vasculopathy [details] | 1 | 0.25 | 11.64423 | 0.0833 | 1 | 0.2689 |
| Encephalomyelitis [details] | 1 | 0.25 | 11.64423 | 0.0833 | 1 | 0.2689 |
| Fatty Liver [unspecific] [details] | 1 | 0.25 | 11.64423 | 0.0833 | 1 | 0.2689 |
| Meningioma [details] | 1 | 0.25 | 11.64423 | 0.0833 | 1 | 0.2689 |
| Pheochromocytoma [details] | 1 | 0.25 | 11.64423 | 0.0833 | 1 | 0.2689 |
| Carcinoma, Supraglottic [details] | 1 | 0.2 | 9.31538 | 0.103 | 1 | 0.3087 |
| Cerebral Cavernous Malformations [details] | 1 | 0.2 | 9.31538 | 0.103 | 1 | 0.3087 |
| Early-Stage Colon Carcinoma [details] | 1 | 0.2 | 9.31538 | 0.103 | 1 | 0.3087 |
| IgA Nephropathy [details] | 1 | 0.2 | 9.31538 | 0.103 | 1 | 0.3087 |
| Panic Disorder [details] | 1 | 0.2 | 9.31538 | 0.103 | 1 | 0.3087 |
| Pediatric Astrocytoma [details] | 1 | 0.2 | 9.31538 | 0.103 | 1 | 0.3087 |
| Tumor Metastasis, to Brain [details] | 1 | 0.2 | 9.31538 | 0.103 | 1 | 0.3087 |
| Biliary Atresia [details] | 1 | 0.16667 | 7.76282 | 0.1223 | 1 | 0.3519 |
| Carcinoma, Gastrointestinal [details] | 1 | 0.16667 | 7.76282 | 0.1223 | 1 | 0.3519 |
| Digestive System Neoplasms [details] | 1 | 0.16667 | 7.76282 | 0.1223 | 1 | 0.3519 |
| Early-Stage Cervical Squamous Cell Carcinoma [details] | 1 | 0.16667 | 7.76282 | 0.1223 | 1 | 0.3519 |
| Essential Thrombocythemia [details] | 1 | 0.16667 | 7.76282 | 0.1223 | 1 | 0.3519 |
| Fibromyalgia [details] | 1 | 0.16667 | 7.76282 | 0.1223 | 1 | 0.3519 |
| Nephrosclerosis [details] | 1 | 0.16667 | 7.76282 | 0.1223 | 1 | 0.3519 |
| Pancreatic Intraductal Papillary Mucinous Neoplasms [details] | 1 | 0.16667 | 7.76282 | 0.1223 | 1 | 0.3519 |
| Toxoplasma gondii Infection [details] | 1 | 0.16667 | 7.76282 | 0.1223 | 1 | 0.3519 |
| Acute Pancreatitis [details] | 1 | 0.14286 | 6.65385 | 0.1413 | 1 | 0.3924 |
| Kidney Diseases [unspecific] [details] | 1 | 0.14286 | 6.65385 | 0.1413 | 1 | 0.3924 |
| Lymphoma, Large B-Cell [details] | 1 | 0.14286 | 6.65385 | 0.1413 | 1 | 0.3924 |
| Oral Lichen Planus [details] | 1 | 0.14286 | 6.65385 | 0.1413 | 1 | 0.3924 |
| Retinal Neovascularization [details] | 1 | 0.14286 | 6.65385 | 0.1413 | 1 | 0.3924 |
| Arrhythmia [details] | 1 | 0.125 | 5.82212 | 0.1598 | 1 | 0.434 |
| Hemophagocytic Lymphohistiocytosis [details] | 1 | 0.125 | 5.82212 | 0.1598 | 1 | 0.434 |
| Muscle Diseases [unspecific] [details] | 1 | 0.125 | 5.82212 | 0.1598 | 1 | 0.434 |
| Tumor Metastasis, to Liver [details] | 1 | 0.125 | 5.82212 | 0.1598 | 1 | 0.434 |
| Wound Inflammation [details] | 1 | 0.125 | 5.82212 | 0.1598 | 1 | 0.434 |
| Alopecia [details] | 1 | 0.11111 | 5.17521 | 0.178 | 1 | 0.4716 |
| Carcinoma, Thyroid, Follicular [details] | 1 | 0.11111 | 5.17521 | 0.178 | 1 | 0.4716 |
| Cardiac Myocyte Injury [details] | 1 | 0.11111 | 5.17521 | 0.178 | 1 | 0.4716 |
| Esophageal Neoplasms [details] | 1 | 0.11111 | 5.17521 | 0.178 | 1 | 0.4716 |
| Human Cytomegalovirus Infection [details] | 1 | 0.11111 | 5.17521 | 0.178 | 1 | 0.4716 |
| Leukemia-Lymphoma, Adult T-Cell [details] | 1 | 0.11111 | 5.17521 | 0.178 | 1 | 0.4716 |
| Leukemia-Lymphoma, Precursor B-Cell Lymphoblastic [details] | 1 | 0.11111 | 5.17521 | 0.178 | 1 | 0.4716 |
| Salivary Gland Neoplasms [details] | 1 | 0.11111 | 5.17521 | 0.178 | 1 | 0.4716 |
| Alcoholic Hepatitis [details] | 1 | 0.1 | 4.65769 | 0.1958 | 1 | 0.5076 |
| Carotid Atherosclerosis [details] | 1 | 0.1 | 4.65769 | 0.1958 | 1 | 0.5076 |
| Early-Stage Gastric Carcinoma [details] | 1 | 0.1 | 4.65769 | 0.1958 | 1 | 0.5076 |
| Graves\' Disease [details] | 1 | 0.1 | 4.65769 | 0.1958 | 1 | 0.5076 |
| Carcinoma, Thyroid, Medullary [details] | 1 | 0.09091 | 4.23427 | 0.2132 | 1 | 0.5469 |
| Gastric Neoplasms [details] | 1 | 0.09091 | 4.23427 | 0.2132 | 1 | 0.5469 |
| Kidney Transplant Rejection [details] | 1 | 0.09091 | 4.23427 | 0.2132 | 1 | 0.5469 |
| Necrotizing Enterocolitis [details] | 1 | 0.09091 | 4.23427 | 0.2132 | 1 | 0.5469 |
| Colitis [details] | 1 | 0.08333 | 3.88141 | 0.2302 | 1 | 0.5845 |
| Hepatitis [unspecific] [details] | 1 | 0.08333 | 3.88141 | 0.2302 | 1 | 0.5845 |
| Leukemia, Lymphoblastic, Acute, Childhood [details] | 1 | 0.08333 | 3.88141 | 0.2302 | 1 | 0.5845 |
| Renal Fibrosis [details] | 1 | 0.08333 | 3.88141 | 0.2302 | 1 | 0.5845 |
| Traumatic Brain Injury [details] | 1 | 0.08333 | 3.88141 | 0.2302 | 1 | 0.5845 |
| Hemoglobin Diseases [details] | 1 | 0.07692 | 3.58284 | 0.2469 | 1 | 0.6191 |

| | | | | | | |
|---|---|---|---|---|---|---|
| Intervertebral Disc Degeneration [details] | 1 | 0.07692 | 3.58284 | 0.2469 | 1 | 0.6191 |
| Leukemia, Lymphoblastic, Acute, B-Cell [details] | 1 | 0.07692 | 3.58284 | 0.2469 | 1 | 0.6191 |
| Mycobacterium Tuberculosis Infection [details] | 1 | 0.07692 | 3.58284 | 0.2469 | 1 | 0.6191 |
| Chordoma [details] | 1 | 0.07143 | 3.32692 | 0.2633 | 1 | 0.6546 |
| Mandibular Prognathism [details] | 1 | 0.07143 | 3.32692 | 0.2633 | 1 | 0.6546 |
| Myotonic Muscular Dystrophy [details] | 1 | 0.07143 | 3.32692 | 0.2633 | 1 | 0.6546 |
| Polycystic Kidney Disease [details] | 1 | 0.07143 | 3.32692 | 0.2633 | 1 | 0.6546 |
| Fatty Liver, Non-Alcoholic [details] | 1 | 0.06667 | 3.10513 | 0.2793 | 1 | 0.686 |
| Prolactinoma [details] | 1 | 0.06667 | 3.10513 | 0.2793 | 1 | 0.686 |
| Myocardial Ischemic-Reperfusion Injury [details] | 1 | 0.0625 | 2.91106 | 0.2949 | 1 | 0.7215 |
| Acute Kidney Injury [details] | 1 | 0.05882 | 2.73982 | 0.3103 | 1 | 0.7575 |
| Focal Segmental Glomerulosclerosis [details] | 1 | 0.05882 | 2.73982 | 0.3103 | 1 | 0.7575 |
| Graft-Versus-Host Disease [details] | 1 | 0.05556 | 2.58761 | 0.3253 | 1 | 0.7894 |
| Mesial Temporal Lobe Epilepsy [details] | 1 | 0.05556 | 2.58761 | 0.3253 | 1 | 0.7894 |
| Muscular Dystrophy, Facioscapulohumeral [details] | 1 | 0.05556 | 2.58761 | 0.3253 | 1 | 0.7894 |
| Essential Hypertension [details] | 1 | 0.05263 | 2.45142 | 0.34 | 1 | 0.8185 |
| Neurodegenerative Diseases [unspecific] [details] | 1 | 0.05263 | 2.45142 | 0.34 | 1 | 0.8185 |
| Allergic Rhinitis [details] | 1 | 0.04762 | 2.21795 | 0.3685 | 1 | 0.8836 |
| Acute Coronary Syndrome [details] | 1 | 0.04545 | 2.11713 | 0.3823 | 1 | 0.9149 |
| Hirschsprung\'s Disease [details] | 1 | 0.04545 | 2.11713 | 0.3823 | 1 | 0.9149 |
| Intrahepatic Cholangiocarcinoma [details] | 1 | 0.04545 | 2.11713 | 0.3823 | 1 | 0.9149 |
| Diabetic Nephropathy [details] | 1 | 0.04167 | 1.94071 | 0.409 | 1 | 0.9712 |
| Medulloblastoma [details] | 1 | 0.03333 | 1.55256 | 0.4827 | 1 | 1 |

**Category: Family (12 Items)**

| | | | | | | |
|---|---|---|---|---|---|---|
| mir-29 family [details] | 3 | 0.75 | 34.93269 | 3.47e-5 | 0.0178 | 4.43e-4 |
| mir-17 family [details] | 3 | 0.375 | 17.46635 | 4.59e-4 | 0.235 | 3.52e-3 |
| mir-221 family [details] | 2 | 1 | 46.57692 | 4.44e-4 | 0.2271 | 3.68e-3 |
| mir-24 family [details] | 2 | 1 | 46.57692 | 4.44e-4 | 0.2271 | 3.68e-3 |
| mir-365 family [details] | 2 | 1 | 46.57692 | 4.44e-4 | 0.2271 | 3.68e-3 |
| mir-15 family [details] | 2 | 0.4 | 18.63077 | 4.26e-3 | 1 | 0.0241 |
| let-7 family [details] | 2 | 0.16667 | 7.76282 | 0.0256 | 1 | 0.1082 |
| mir-23 family [details] | 1 | 0.5 | 23.28846 | 0.0425 | 1 | 0.1655 |
| mir-130 family [details] | 1 | 0.25 | 11.64423 | 0.0833 | 1 | 0.2689 |
| mir-25 family [details] | 1 | 0.25 | 11.64423 | 0.0833 | 1 | 0.2689 |
| mir-30 family [details] | 1 | 0.16667 | 7.76282 | 0.1223 | 1 | 0.3519 |
| mir-320 family [details] | 1 | 0.125 | 5.82212 | 0.1598 | 1 | 0.434 |

**Category: Function (95 Items)**

| | | | | | | |
|---|---|---|---|---|---|---|
| Cell Death [details] | 18 | 0.23077 | 10.74852 | 5.20e-17 | 2.66e-14 | 6.30e-14 |
| Apoptosis [details] | 14 | 0.13208 | 6.15167 | 2.69e-9 | 1.38e-6 | 1.48e-7 |
| Cell Proliferation [details] | 13 | 0.1625 | 7.56875 | 9.02e-10 | 4.62e-7 | 6.82e-8 |
| Cell Cycle [details] | 13 | 0.15663 | 7.29518 | 1.47e-9 | 7.51e-7 | 8.45e-8 |
| Inflammation [details] | 12 | 0.10714 | 4.99038 | 7.05e-7 | 3.61e-4 | 1.45e-5 |
| Hormone-mediated Signaling Pathway [details] | 11 | 0.18966 | 8.83355 | 5.16e-9 | 2.64e-6 | 2.40e-7 |
| Angiogenesis [details] | 11 | 0.16923 | 7.88225 | 1.87e-8 | 9.55e-6 | 6.46e-7 |
| Onco-MiRNAs [details] | 9 | 0.23077 | 10.74852 | 3.00e-8 | 1.54e-5 | 9.08e-7 |
| Regulation of Stem Cell [details] | 9 | 0.11392 | 5.30623 | 1.72e-5 | 8.79e-3 | 2.50e-4 |
| Immune Response [details] | 9 | 0.09783 | 4.55644 | 6.08e-5 | 0.0312 | 6.95e-4 |
| Aging [details] | 8 | 0.12698 | 5.91453 | 2.55e-5 | 0.0131 | 3.40e-4 |
| Epithelial-to-Mesenchymal Transition [details] | 8 | 0.09639 | 4.48934 | 1.97e-4 | 0.1009 | 1.84e-3 |
| Bone Regeneration [details] | 7 | 0.23333 | 10.86795 | 1.30e-6 | 6.67e-4 | 2.63e-5 |
| Adipocyte Differentiation [details] | 7 | 0.17073 | 7.95216 | 1.23e-5 | 6.31e-3 | 1.89e-4 |
| Hematopoiesis [details] | 7 | 0.12281 | 5.71997 | 1.15e-4 | 0.0591 | 1.18e-3 |
| Neuron Apoptosis [details] | 6 | 0.4 | 18.63077 | 2.34e-7 | 1.20e-4 | 5.67e-6 |
| Nephrotoxicity [details] | 6 | 0.35294 | 16.43891 | 5.62e-7 | 2.88e-4 | 1.19e-5 |
| Toxicity [details] | 6 | 0.16667 | 7.76282 | 6.73e-5 | 0.0344 | 7.61e-4 |
| Innate Immunity [details] | 6 | 0.14286 | 6.65385 | 1.66e-4 | 0.085 | 1.60e-3 |
| Muscle Development [details] | 5 | 0.2381 | 11.08974 | 4.92e-5 | 0.0252 | 5.84e-4 |

| | | | | | | |
|---|---|---|---|---|---|---|
| Wound Healing [details] | 5 | 0.21739 | 10.12542 | 7.90e-5 | 0.0404 | 8.62e-4 |
| Cell Differentiation [details] | 5 | 0.08929 | 4.15865 | 5.51e-3 | 1 | 0.0293 |
| Tumor Suppressor MiRNAs [details] | 5 | 0.07692 | 3.58284 | 0.0104 | 1 | 0.0492 |
| DNA Damage Response [details] | 4 | 0.25 | 11.64423 | 2.56e-4 | 0.131 | 2.31e-3 |
| T-Cell Differentiation [details] | 4 | 0.25 | 11.64423 | 2.56e-4 | 0.131 | 2.31e-3 |
| Cell Division [details] | 4 | 0.23529 | 10.95928 | 3.30e-4 | 0.1688 | 2.85e-3 |
| DNA Damage Repair [details] | 4 | 0.21053 | 9.80567 | 5.21e-4 | 0.2669 | 3.80e-3 |
| T-helper 17 Cell Differentiation [details] | 4 | 0.21053 | 9.80567 | 5.21e-4 | 0.2669 | 3.80e-3 |
| Neurotoxicity [details] | 4 | 0.2 | 9.31538 | 6.42e-4 | 0.3288 | 4.57e-3 |
| Pluripotent Stem Cells Reprogramming [details] | 4 | 0.2 | 9.31538 | 6.42e-4 | 0.3288 | 4.57e-3 |
| Immune System(Xiao\'s Cell2010) [details] | 4 | 0.19048 | 8.87179 | 7.82e-4 | 0.4002 | 5.35e-3 |
| Regulation of Akt Pathway [details] | 4 | 0.15385 | 7.16568 | 1.81e-3 | 0.9288 | 0.0111 |
| Embryonic Stem Cell Differentiation [details] | 4 | 0.12903 | 6.00993 | 3.55e-3 | 1 | 0.0204 |
| Brain Development [details] | 4 | 0.11111 | 5.17521 | 6.17e-3 | 1 | 0.0322 |
| Osteogenesis [details] | 4 | 0.0678 | 3.15776 | 0.034 | 1 | 0.1382 |
| Cell Adhesion [details] | 3 | 0.375 | 17.46635 | 4.59e-4 | 0.235 | 3.52e-3 |
| Chemosensitivity Of Tumor Cells [details] | 3 | 0.375 | 17.46635 | 4.59e-4 | 0.235 | 3.52e-3 |
| Myogensis [details] | 3 | 0.375 | 17.46635 | 4.59e-4 | 0.235 | 3.52e-3 |
| Cardiac Remodeling [details] | 3 | 0.33333 | 15.52564 | 6.79e-4 | 0.3474 | 4.78e-3 |
| Folliculogenesis [details] | 3 | 0.27273 | 12.7028 | 1.30e-3 | 0.6631 | 8.48e-3 |
| Vascular Inflammation [details] | 3 | 0.2 | 9.31538 | 3.37e-3 | 1 | 0.0195 |
| Latent Virus Replication [details] | 3 | 0.17647 | 8.21946 | 4.90e-3 | 1 | 0.0264 |
| Adipogenesis [details] | 3 | 0.15 | 6.98654 | 7.87e-3 | 1 | 0.0394 |
| Cell Motility [details] | 3 | 0.14286 | 6.65385 | 9.05e-3 | 1 | 0.0436 |
| Circadian Rhythm [details] | 3 | 0.13636 | 6.3514 | 0.0103 | 1 | 0.0492 |
| Osteoblast Differentiation [details] | 3 | 0.12 | 5.58923 | 0.0148 | 1 | 0.0678 |
| Glucose Metabolism [details] | 3 | 0.10714 | 4.99038 | 0.0202 | 1 | 0.0898 |
| Collagen Formation [details] | 2 | 1 | 46.57692 | 4.44e-4 | 0.2271 | 3.68e-3 |
| Extracellular Matrix Remodeling [details] | 2 | 0.4 | 18.63077 | 4.26e-3 | 1 | 0.0241 |
| Cardiomyocyte Apoptosis [details] | 2 | 0.28571 | 13.30769 | 8.72e-3 | 1 | 0.0429 |
| Anti-Cell Proliferation(Hwang Etal Bjc2007) [details] | 2 | 0.25 | 11.64423 | 0.0115 | 1 | 0.0538 |
| Transdifferentiation [details] | 2 | 0.25 | 11.64423 | 0.0115 | 1 | 0.0538 |
| Granulopoiesis [details] | 2 | 0.2 | 9.31538 | 0.018 | 1 | 0.0805 |
| Oxidative Stress [details] | 2 | 0.2 | 9.31538 | 0.018 | 1 | 0.0805 |
| Vascular Homeostasis [details] | 2 | 0.16667 | 7.76282 | 0.0256 | 1 | 0.1082 |
| Chondrocyte Development [details] | 2 | 0.13333 | 6.21026 | 0.0392 | 1 | 0.1557 |
| Embryonic Development [details] | 2 | 0.11765 | 5.47964 | 0.0495 | 1 | 0.1747 |
| Smooth Muscle Cell Proliferation [details] | 2 | 0.11111 | 5.17521 | 0.0549 | 1 | 0.1923 |
| Skeletal Muscle Cell Differentiation [details] | 2 | 0.09091 | 4.23427 | 0.0787 | 1 | 0.2556 |
| Antiviral Immunity [details] | 1 | 0.5 | 23.28846 | 0.0425 | 1 | 0.1655 |
| Cardiomyocyte Migration [details] | 1 | 0.5 | 23.28846 | 0.0425 | 1 | 0.1655 |
| DNA Synthesis [details] | 1 | 0.5 | 23.28846 | 0.0425 | 1 | 0.1655 |
| Muscle Regeneration [details] | 1 | 0.5 | 23.28846 | 0.0425 | 1 | 0.1655 |
| Osteoblast Apoptosis [details] | 1 | 0.5 | 23.28846 | 0.0425 | 1 | 0.1655 |
| Cardiomyocyte Differentiation [details] | 1 | 0.33333 | 15.52564 | 0.0631 | 1 | 0.2177 |
| Genomic Instability [details] | 1 | 0.33333 | 15.52564 | 0.0631 | 1 | 0.2177 |
| Myoblast Differentiation [details] | 1 | 0.33333 | 15.52564 | 0.0631 | 1 | 0.2177 |
| Pancreas Development [details] | 1 | 0.33333 | 15.52564 | 0.0631 | 1 | 0.2177 |
| Radiation-Induced Bystander Effects [details] | 1 | 0.33333 | 15.52564 | 0.0631 | 1 | 0.2177 |
| Smooth Muscle Cell Differentiation [details] | 1 | 0.33333 | 15.52564 | 0.0631 | 1 | 0.2177 |
| Tumor Cell Radiation Sensitivity [details] | 1 | 0.33333 | 15.52564 | 0.0631 | 1 | 0.2177 |
| Cardiomyocyte Proliferation [details] | 1 | 0.25 | 11.64423 | 0.0833 | 1 | 0.2689 |
| Cell Proliferation(Hwang Etal Bjc2007) [details] | 1 | 0.25 | 11.64423 | 0.0833 | 1 | 0.2689 |
| Cleavage Stage Development [details] | 1 | 0.25 | 11.64423 | 0.0833 | 1 | 0.2689 |
| Dendritic Cell Differentiation [details] | 1 | 0.25 | 11.64423 | 0.0833 | 1 | 0.2689 |
| Mesenchymal-to-Epithelial Transition [details] | 1 | 0.25 | 11.64423 | 0.0833 | 1 | 0.2689 |
| Type II Pneumocyte Differentiation [details] | 1 | 0.25 | 11.64423 | 0.0833 | 1 | 0.2689 |
| Hepatic Stellate Cell Differentiation [details] | 1 | 0.2 | 9.31538 | 0.103 | 1 | 0.3087 |

| | | | | | | |
|---|---|---|---|---|---|---|
| Natural Killer Cell Activation [details] | 1 | 0.2 | 9.31538 | 0.103 | 1 | 0.3087 |
| Chondrogenic Differentiation [details] | 1 | 0.16667 | 7.76282 | 0.1223 | 1 | 0.3519 |
| Hepatotoxicity [details] | 1 | 0.16667 | 7.76282 | 0.1223 | 1 | 0.3519 |
| Cellular Senescence [details] | 1 | 0.14286 | 6.65385 | 0.1413 | 1 | 0.3924 |
| Chromatin Remodeling [details] | 1 | 0.14286 | 6.65385 | 0.1413 | 1 | 0.3924 |
| Regulation of Nf-Kb Pathway [details] | 1 | 0.14286 | 6.65385 | 0.1413 | 1 | 0.3924 |
| Autophagy [details] | 1 | 0.125 | 5.82212 | 0.1598 | 1 | 0.434 |
| Mesenchymal Stem Cell Proliferation [details] | 1 | 0.125 | 5.82212 | 0.1598 | 1 | 0.434 |
| Response to Estrogen [details] | 1 | 0.125 | 5.82212 | 0.1598 | 1 | 0.434 |
| T-Cell Activation [details] | 1 | 0.125 | 5.82212 | 0.1598 | 1 | 0.434 |
| Carbohydrate Metabolism [details] | 1 | 0.11111 | 5.17521 | 0.178 | 1 | 0.4716 |
| Heart Development [details] | 1 | 0.09091 | 4.23427 | 0.2132 | 1 | 0.5469 |
| Osteoclast Differentiation [details] | 1 | 0.07143 | 3.32692 | 0.2633 | 1 | 0.6546 |
| Cardiotoxicity [details] | 1 | 0.05882 | 2.73982 | 0.3103 | 1 | 0.7575 |
| Insulin Resistance [details] | 1 | 0.05556 | 2.58761 | 0.3253 | 1 | 0.7894 |
| Peritoneal Cavity Homeostasis(26495316) [details] | 1 | 0.04348 | 2.02508 | 0.3958 | 1 | 0.9416 |
| Lipid Metabolism [details] | 1 | 0.02222 | 1.03504 | 0.6303 | 1 | 1 |

**Category: Transcription Factor (45 Items)**

| | | | | | | |
|---|---|---|---|---|---|---|
| NFKB1 [details] | 9 | 0.34615 | 16.12278 | 5.24e-10 | 2.68e-7 | 4.23e-8 |
| MYC [details] | 9 | 0.225 | 10.47981 | 3.82e-8 | 1.96e-5 | 1.13e-6 |
| E2F1 [details] | 7 | 0.26923 | 12.53994 | 4.46e-7 | 2.28e-4 | 9.81e-6 |
| ERS1 [details] | 6 | 0.4 | 18.63077 | 2.34e-7 | 1.20e-4 | 5.67e-6 |
| TGFB1 [details] | 5 | 0.25 | 11.64423 | 3.80e-5 | 0.0195 | 4.70e-4 |
| SP1 [details] | 4 | 0.4 | 18.63077 | 3.22e-5 | 0.0165 | 4.15e-4 |
| PDGF [details] | 3 | 1 | 46.57692 | 8.81e-6 | 4.51e-3 | 1.42e-4 |
| AP-1 [details] | 3 | 0.75 | 34.93269 | 3.47e-5 | 0.0178 | 4.43e-4 |
| YY1 [details] | 3 | 0.375 | 17.46635 | 4.59e-4 | 0.235 | 3.52e-3 |
| PTEN [details] | 3 | 0.33333 | 15.52564 | 6.79e-4 | 0.3474 | 4.78e-3 |
| MYCN [details] | 3 | 0.3 | 13.97308 | 9.56e-4 | 0.4893 | 6.43e-3 |
| CEBPA [details] | 3 | 0.27273 | 12.7028 | 1.30e-3 | 0.6631 | 8.48e-3 |
| TP53 [details] | 3 | 0.15 | 6.98654 | 7.87e-3 | 1 | 0.0394 |
| FOSL1 [details] | 2 | 1 | 46.57692 | 4.44e-4 | 0.2271 | 3.68e-3 |
| P27 [details] | 2 | 1 | 46.57692 | 4.44e-4 | 0.2271 | 3.68e-3 |
| SF2/ASF [details] | 2 | 0.4 | 18.63077 | 4.26e-3 | 1 | 0.0241 |
| HMGA1 [details] | 2 | 0.33333 | 15.52564 | 6.31e-3 | 1 | 0.0328 |
| E2F3 [details] | 2 | 0.28571 | 13.30769 | 8.72e-3 | 1 | 0.0429 |
| EIF2C2 [details] | 2 | 0.18182 | 8.46853 | 0.0217 | 1 | 0.095 |
| LIN28 [details] | 2 | 0.18182 | 8.46853 | 0.0217 | 1 | 0.095 |
| LIN28B [details] | 2 | 0.18182 | 8.46853 | 0.0217 | 1 | 0.095 |
| SPI1 [details] | 2 | 0.18182 | 8.46853 | 0.0217 | 1 | 0.095 |
| TRIM32 [details] | 2 | 0.18182 | 8.46853 | 0.0217 | 1 | 0.095 |
| EGR1 [details] | 2 | 0.06897 | 3.2122 | 0.1264 | 1 | 0.3518 |
| BMP2 [details] | 1 | 0.5 | 23.28846 | 0.0425 | 1 | 0.1655 |
| CCND1 [details] | 1 | 0.5 | 23.28846 | 0.0425 | 1 | 0.1655 |
| DDX5 [details] | 1 | 0.5 | 23.28846 | 0.0425 | 1 | 0.1655 |
| GFI1 [details] | 1 | 0.5 | 23.28846 | 0.0425 | 1 | 0.1655 |
| ETS1 [details] | 1 | 0.33333 | 15.52564 | 0.0631 | 1 | 0.2177 |
| TCF4 [details] | 1 | 0.33333 | 15.52564 | 0.0631 | 1 | 0.2177 |
| NANOG [details] | 1 | 0.25 | 11.64423 | 0.0833 | 1 | 0.2689 |
| NKX2-5 [details] | 1 | 0.25 | 11.64423 | 0.0833 | 1 | 0.2689 |
| TLX1 [details] | 1 | 0.25 | 11.64423 | 0.0833 | 1 | 0.2689 |
| TLX3 [details] | 1 | 0.25 | 11.64423 | 0.0833 | 1 | 0.2689 |
| BMP4 [details] | 1 | 0.2 | 9.31538 | 0.103 | 1 | 0.3087 |
| KLF4 [details] | 1 | 0.2 | 9.31538 | 0.103 | 1 | 0.3087 |
| REST [details] | 1 | 0.2 | 9.31538 | 0.103 | 1 | 0.3087 |
| RUNX2 [details] | 1 | 0.2 | 9.31538 | 0.103 | 1 | 0.3087 |
| STAT3 [details] | 1 | 0.2 | 9.31538 | 0.103 | 1 | 0.3087 |

| | | | | | | |
|---|---|---|---|---|---|---|
| AKT1 [details] | 1 | 0.16667 | 7.76282 | 0.1223 | 1 | 0.3519 |
| STAT5 [details] | 1 | 0.14286 | 6.65385 | 0.1413 | 1 | 0.3924 |
| TP63 [details] | 1 | 0.14286 | 6.65385 | 0.1413 | 1 | 0.3924 |
| TWIST1 [details] | 1 | 0.1 | 4.65769 | 0.1958 | 1 | 0.5076 |
| SOX2 [details] | 1 | 0.08333 | 3.88141 | 0.2302 | 1 | 0.5845 |
| ZEB1 [details] | 1 | 0.07143 | 3.32692 | 0.2633 | 1 | 0.6546 |
